# Theory on the rate equations of Michaelis-Menten type enzyme kinetics with competitive inhibition

**DOI:** 10.1101/2022.11.28.518182

**Authors:** R. Murugan

## Abstract

We derive approximate expressions under various conditions of validity over both pre- and post-steady state regimes of the velocity-substrate-inhibitor spaces of the Michaelis-Menten enzyme kinetic schemes with fully and partial competitive inhibition. Our refinement over the currently available standard quasi steady state approximations (sQSSA) seems to be valid over wide range of enzyme to substrate and enzyme to inhibitor ratios. Further, we show that under certain conditions the enzyme-inhibitor-substrate system can exhibit temporally well-separated two different steady states with respect to both enzyme-substrate and enzyme-inhibitor complexes. We define the ratios *f*_*S*_ = *v*_*max*_⁄(*K*_*MS*_ + *e*_0_) and *f*_*I*_ = *u*_*max*_⁄(*K*_*MI*_ + *e*_0_) as the acceleration factors with respect to the catalytic conversion of substrate and inhibitor into their respective products. Here *K*_*MS*_ and *K*_*MI*_ are the Michaelis-Menten parameters associated with the binding of substrate and inhibitor with the enzyme, *v*_*max*_ and *u*_*ma*x_ are the respective maximum reaction velocities and e_0_, s_0_, and i_0_ are total enzyme, substrate and inhibitor levels. When (*f*_*S*_ ⁄ *f*_*I*_) < 1, then enzyme-substrate complex will show multiple steady states subsequently reaches the full-fledged steady state only after the depletion of enzyme-inhibitor complex. When (*f*_*S*_ ⁄ *f*_*I*_) > 1, then the enzyme-inhibitor complex will show multiple steady states and subsequently reaches the full-fledged steady state only after the depletion of enzyme-substrate complex. This complex behavior exclusively when (*f*_*S*_ ⁄ *f*_*I*_) ≠ 1 is the root cause of large amount of error in the estimation of various kinetic parameters both in the cases of fully and partial competitive inhibition schemes using the sQSSA methods. Remarkably, we show that our refined expressions for the reaction velocities over enzyme-substrate-inhibitor space can control this error more significantly than the currently available sQSSA velocity expressions.

## 1. Introduction

Enzymes catalyze various reactions of the biochemical pathways [1-4]. The **M**ichaelis-**M**enten (MM) kinetics [5, 6] is the fundamental mechanistic description of the biological catalysis of enzyme reactions [3, 7-9]. In this kinetics scheme, the enzyme reversibly binds its substrate to form the enzyme-substrate complex which subsequently decompose into free enzyme and product of the substrate. Integral solution to the rate equations associated with the Michaelis-Menten scheme (**MMS**) is not expressible in terms of elementary functions. Several analytical methods were tried to obtain the approximate solution of MMS in terms of ordinary [10] and singular perturbation series [11-13] and perturbation expansions over slow manifolds [14, 15]. In general, the singular perturbation expansions yield a combination of inner and outer solutions which were then combined via proper matching at the boundary layer [11, 16-21].

Several steady state approximations were proposed in the light of experimental characterization of a single substrate MM enzyme. The **standard quasi steady state approximation** (sQSSA) is widely used across several fields of biochemical research to obtain the enzyme kinetic parameters such as v_max_ and K_M_ from the experimental datasets on the reaction velocity versus initial substrate concentrations. This approximation works well when the product formation step is a rate limiting one apart from the condition that the substrate concentration is much higher than the enzyme concentration. In general, sQSSA yields expressions which can be directly used by the experimentalists to obtain various enzyme kinetic parameters [22]. Recently, explicit closed form expressions of the integrated rate equation corresponding to sQSSA were obtained in terms of **Lambert’s W** functions [23-27]. The **total QSSA** (tQSSA) assumes that the amount of product formed near the steady state is much negligible compared to the total substrate concentration [28, 29]. The **reverse QSSA** (rQSSA) works very well [30, 31] when the substrate concentration is much lesser than the enzyme concentration.

Several linearization techniques such as Lineweaver-Burk representation were also proposed [32, 33] to obtain the kinetic parameters from the experimental data. Although sQSSA, rQSSA and tQSSA methods work well under *in vitro* conditions, there are several situations such as single molecule enzyme kinetics [34] and other *in vivo* experimental conditions where one cannot manipulate the ratio of substrate to enzyme concentrations much. Further, successfulness of various QSSAs in accurately obtaining the kinetic parameters is strongly dependent on the timescale separation between the pre- and post-steady state regimes of MMS [35, 36]. Particularly, when the timescale separation between pre- and post-steady states of MMS is high enough, then the sQSSA along with **stationary reactant assumption** where one replaces the unknown steady state substrate concentration with the total substrate concentration [27] can be used to directly obtain the kinetic parameters.

The catalytic properties of an enzyme can be manipulated by an inhibitor. Inhibitors can be competitive or allosteric in nature [2]. Competitive inhibitors (**Fig. 1**) are substrate like molecules which reversibly bind the active site of the same enzyme and hence block the further binding of substrate. This in turn deceases the catalytic efficiency of the enzyme over substrate. In a fully competitive inhibition (**Scheme A** in **Fig. 1**), the inhibitor competes with the substrate to bind the active site of the enzyme and subsequently gets converted into the respective product. In this case, both substrate and inhibitor will be converted into their respective products by the same enzyme. In case of partial competitive inhibition (**Scheme B** in **Fig. 1**), the reversibly formed enzyme-inhibitor complex will be a dead-end one. Several drugs have been designed to strongly inhibit the pathogenic or metabolic enzymes. Understanding the dynamical behavior of the fully and partial competitive inhibition of MM enzymes is critical to understand the pharmacokinetic and efficiency aspects of such enzyme inhibiting drugs. Variation of v_max_ and K_M_ of the enzyme with respect to the concentration of an inhibitor decides the efficiency of a given drug molecule in targeting that enzyme. In the steady state experiments on the single substrate enzymes, the total substrate concentration will be iterated to obtain the respective substrate conversion velocities. This substrate concentration versus reaction velocity dataset will be then used to obtain the kinetic parameters such as K_M_ and v_max_. To obtain the kinetic parameters related to the enzyme inhibition, one needs to conduct a series of velocity versus substrate type steady state experiments at different concentrations of the inhibitor. Using this dataset on the steady state substrate, inhibitor versus reaction velocities, one obtains the kinetic parameters related to the enzyme-substrate-inhibitor system.

**FIGURE 1.**
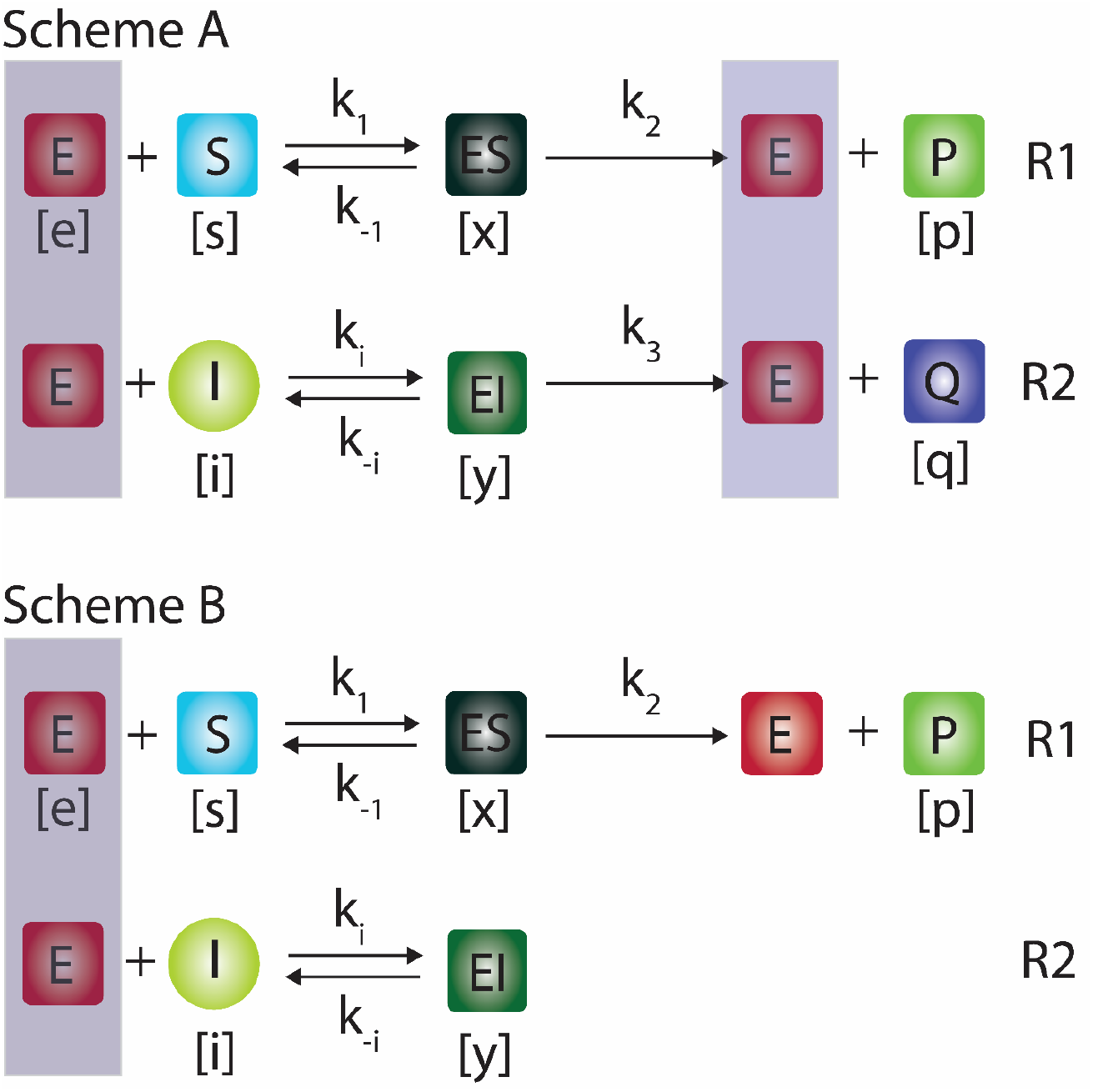
Fully (Scheme A) and partial (Scheme B) competitive inhibition schemes of Michaelis-Menten type enzyme kinetics. In fully competitive inhibition, both substrate (S) and inhibitor (I) compete for the same active site of enzyme (E) to bind and form reversible complexes (ES, EI) which subsequently get converted into their respective products (P, Q). Whereas, in partial competitive inhibition, the reversibly formed enzyme-inhibitor (EI) is a dead-end complex. Here (e, s, i, x, y, p, q) are respectively the concentrations of enzyme, substrate, inhibitor, enzyme-substrate, enzyme-inhibitor, product of substrate and product of inhibitor. Further, k_1_ and k_i_ are the respective forward rate constants, k_-1_ and k_-i_ are the reverse rate constants and, k_2_ and k_3_ are the respective product formation rates.

The successfulness of various steady state approximations in obtaining the kinetic parameters of enzymes from the experimental datasets strongly depends on the occurrence of a common steady state with respect to both the substrate and inhibitor binding dynamics in case of fully and partial competitive inhibition schemes. Mismatch in the steady state timescales can be resolved by setting higher substrate and inhibitor concentrations than the enzyme concentration. This condition drives the steady state reaction velocities as well as the timescales corresponding to both the binding of substrate and inhibitor with the same enzyme close to zero. However, under *in vivo* conditions, one cannot manipulate the relative concentrations of substrate, inhibitor and enzyme much. All the quasi steady state type approximations will fail when the concentration of the enzyme is comparable with that of the substrate and inhibitor which is generally true under *in vivo* conditions. In this article, we will address this issue in detail and derive accurate expressions for the steady state reaction velocities when the concentrations of enzyme, substrate and inhibitor are comparable with each other.

## 2. Theory

The competitive inhibition of Michaelis-Menten enzymes can be via fully or partial mode as depicted in **Scheme A** and **B** respectively in **Fig. 1**. In fully competitive inhibition given in Scheme A, both the substrate and inhibitor molecules compete for the same active site of the target enzyme to bind and subsequently get converted into their respective products in a parallel manner. In case of partial competitive inhibition, the reversibly formed enzyme-inhibitor complex will not be converted into any product and it will be a dead-end one. Particularly, several drug molecules are partial competitive inhibitors. Fully competitive inhibition plays important roles in the regulation of metabolic reaction pathways. In the following sections we will analyze various kinetic aspects of fully and partial competitive inhibition schemes in detail. We use the equation numbering as the section number followed by the respective equation number within that section e.g., in the notation **Eq. <u>x.y.z</u>.k**, <u>x.y.z</u> is the section number and k is the equation number in that section.

### 2.1. Fully competitive inhibition

The fully competitive inhibition of Michaelis-Menten enzymes that is depicted in **Scheme A** of **Fig. 1** can be quantitatively described by the following set of differential rate equations.

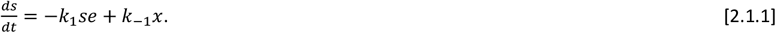

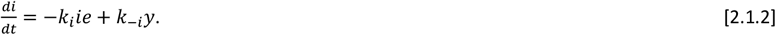

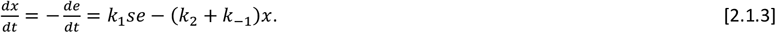

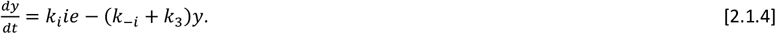

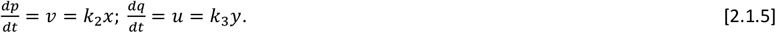

In **Eqs. 2.1.1-5**, (s, i, e, x, y, p, q) are respectively the concentrations (mol/lit, M) of substrate, inhibitor, enzyme, enzyme-substrate complex, enzyme-inhibitor complex, product of substrate and product of inhibitor. Here k_1_ and k_i_ are the respective forward bimolecular rate constants (1/M/second), k_-1_, k_-i_ (1/second) are the respective reverse unimolecular rate constants, u and v (M/second) are the respective reaction velocities and k_2_, k_3_ (1/second) are the respective unimolecular product formation rate constants along with the mass conservation laws: *e* = *e*_0_ − *x* − *y* ; *s* = *s*_0_ − *x* − *p* ; *i* = *i*_0_ − *y* − *q*. The initial conditions are (s, i, e, x, y, p, q, v, u) = (s_0_, i_0_, e_0_, 0, 0, 0, 0, 0, 0) at t = 0. When *t* → ∞, then the reaction ends at (s, i, e, x, y, p, q, v, u) = (0, 0, e_0_, 0, 0, s_0_, i_0_, 0, 0). The steady states occur when 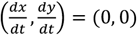 especially under the condition that *t*< ∞ since 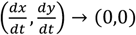 when *t* → ∞. However, the timescale 0 < *t*_*CS*_ < ∞ at which 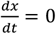 can be different from the timescale 0 < *t*_*CI*_ < ∞ at which 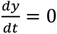. When there is a mismatch in the steady state timescales (*t*_*CS*_ ≠ *t*_*CI*_), then one cannot obtain a common steady state solution to **Eqs. 2.1.1-5** by simultaneously equating all of them to zero. This means that there exist two different steady states with respect to enzyme-substrate and enzyme-inhibitor complexes at two different time points along with four different timescales viz. two different pre-steady state timescales and two different post-steady state timescales. Various definitions and symbols used in the theory section are summarized in **Table 1**.

**TABLE 1.** Summary of variables and parameters used in the theory section.

| Parameters / Variables | Definition | Remarks |
| --- | --- | --- |
| $e, s, i, p, q, x, y, v, u$ | Concentration of enzyme, substrate, inhibitor, product of substrate, product of inhibitor, enzyme-substrate complex, enzyme-inhibitor complex, velocity of substrate-product formation, velocity of inhibitor-product formation. | mol/lit, M |
| $e_0, s_0, i_0$ | Initial enzyme, substrate and inhibitor concentrations. | M |
| $E, S, I, P, Q, X, Y, V, U$ | $S = \frac{s}{s_0}, E = \frac{e}{e_0}, P = \frac{p}{s_0}, Y = \frac{y}{e_0}, X = \frac{x}{e_0}, I = \frac{i}{i_0}, Q = \frac{q}{i_0}$ are the normalized concentrations of substrate, enzyme, product of substrate, enzyme-inhibitor complex, enzyme-substrate complex, inhibitor and product of inhibitor, velocity of product of substrate formation, and velocity of product of inhibitor formation. | dimensionless |
| $E_C, S_C, I_C, P_C, Q_C, X_C, Y_C, V_C, U_C$ | Overall steady state values with respect to both X as well as Y. | dimensionless |
| $S_{CP}, I_{CP}, P_{CP}, Q_{CP}, X_{CP}, Y_{CP}, V_{CP}, U_{CP}$ | Values at the steady state with respect to only X. | dimensionless |
| $S_{CQ}, I_{CQ}, P_{CQ}, Q_{CQ}, X_{CQ}, Y_{CQ}, V_{CQ}, U_{CQ}$ | Values at the steady state with respect to only Y. | dimensionless |
| $k_2, k_3$ | Unimolecular product formation rate constants. | 1/second |
| $\tau$ | $= k_2 t$ | dimensionless |
| $v_{max}, u_{max}$ | $v_{max} = k_2 e_0, u_{max} = k_3 e_0$ | M/second |
| $k_1, k_i$ | Bimolecular rate constants associated with the binding of substrate and inhibitor respectively with the same enzyme. | 1 / (M second) |
| $k_{-1}, k_{-i}$ | Dissociation rate constants associated with the enzyme-substrate and enzyme-inhibitor complexes. | 1/second |
| $K_{RS}, K_{RI}$ | $K_{RS} = k_2 / k_1, K_{RI} = k_3 / k_i$ | M |
| $K_{DS}, K_{DI}$ | $K_{DS} = k_{-1} / k_1, K_{DI} = k_{-i} / k_i$ | M |
| $K_{MS}, K_{MI}$ | $K_{MS} = K_{RS} + K_{DS}, K_{MI} = K_{RI} + K_{DI}$ | M |
| $\eta_S, \eta_I$ | $\eta_S = k_2 / k_1 s_0, \eta_I = k_3 / k_i i_0$ | dimensionless |
| $\epsilon_S, \epsilon_I, \epsilon_{IS}$ | $\epsilon_S = e_0 / s_0, \epsilon_I = e_0 / i_0, \epsilon_{IS} = \epsilon_I / \epsilon_S = s_0 / i_0$ | dimensionless |
| $\kappa_S, \kappa_I$ | $\kappa_S = k_{-1} / k_1 s_0, \kappa_I = k_{-i} / k_i i_0$ | dimensionless |
| $\chi_I$ | $= k_2 / k_i i_0$ defined for the partial competitive inhibition scheme. | dimensionless |
| $\rho$ | $= k_3 / k_2 = u_{max} / v_{max}$ | dimensionless |
| $\sigma$ | $= k_1 / k_i$ | dimensionless |
| $\mu_S, \mu_I$ | $\mu_S = \eta_S + \kappa_S, \mu_I = \eta_I + \kappa_I$ | dimensionless |
| $\Upsilon$ | $= k_{-1} / k_{-i}$ | dimensionless |
| $\delta$ | $= \tilde{\mu}_I \varepsilon_S / \rho \tilde{\mu}_S \varepsilon_I = (v_{max}/\tilde{\mu}_S)/(u_{max}/\tilde{\mu}_I)$ | dimensionless |
| $\tilde{\mu}_I$ | $= \varepsilon_I + \kappa_I + \eta_I = \varepsilon_I + \mu_I$ | dimensionless |
| $\tilde{\mu}_S$ | $= \varepsilon_S + \kappa_S + \eta_S = \varepsilon_S + \mu_S$ | dimensionless |
| $\tilde{\kappa}_I$ | $= \varepsilon_I + \kappa_I$ | dimensionless |
| $\alpha_S$ | $= 1 + \varepsilon_S + \kappa_S + \eta_S$ | dimensionless |
| $\alpha_I$ | $= 1 + \varepsilon_I + \kappa_I + \eta_I$ | dimensionless |
| $\beta_I$ | $= 1 + \varepsilon_I + \kappa_I$ | dimensionless |
| $\phi_S$ | $= \eta_S \varepsilon_S$ | dimensionless |
| $\phi_I$ | $= \eta_I \varepsilon_I$ | dimensionless |
| $\tau_{CS}$ | $\cong \frac{\alpha_I \eta_S}{(\alpha_S \alpha_I - 1)}$, steady state timescale corresponding to the enzyme-substrate complex (fully competitive).<br>$\cong \frac{\beta_I \eta_S}{(\alpha_S \beta_I - 1)}$, steady state timescale corresponding to the enzyme-substrate complex (partial competitive). | dimensionless<br>**Eq. 2.6.1.1.** |
| $\tau_{CI}$ | $\cong \frac{\alpha_S \eta_I}{(\alpha_S \alpha_I - 1)}$, steady state timescale corresponding to the enzyme-inhibitor complex (fully competitive) | dimensionless<br>**Eq. 2.6.1.1.** |
| $\tau_{CY}$ | $\cong 2\chi_I / \sqrt{\beta_I^2 - 4\varepsilon_I}$, (pesudo) steady state timescale corresponding to enzyme-inhibitor complex (partial competitive). | Dimensionless<br>**Eq. 2.9.3.2.** |
| $f_S, f_I$ | $f_S = v_{max}/\tilde{\mu}_S, f_I = u_{max}/\tilde{\mu}_I$ are the reaction acceleration factors associated with the conversion of substrate and inhibitor into their products.<br>$\delta = f_S/f_I$. | M/second. |

### 2.2. Scaling and non-dimensionalization

To simplify the system of **Eqs. 2.1.1-5**, we introduce the following set of scaling transformations.

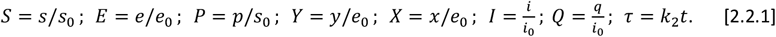

We further define the following parameters.

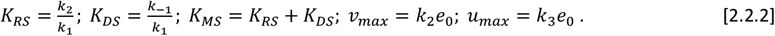

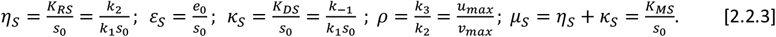

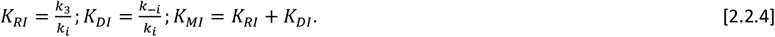

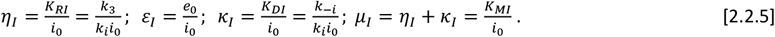

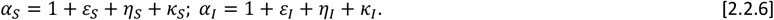

Here (S, I, E, X, Y, P, Q) ∈ [0,1] are the dimensionless time dependent dynamical variables along with the mass conservation laws: *E* = 1 − *X* − *Y* ; *S* = 1 − *ε*_*S*_*X* − *P* ; *I* = 1 − *ε*_*I*_ *Y* − *Q*. With these scaling transformations, one can reduce **Eqs. 2.1.1-5** into the following set of equations.

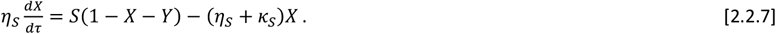

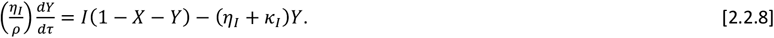

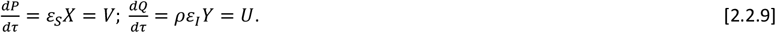

Upon expanding S and I with their definition in the right-hand side of **Eqs. 2.2.7-8** and rearranging the linear and nonlinear terms, we arrive at the following form.

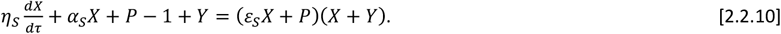

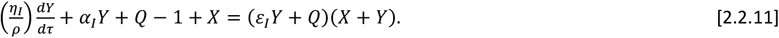

Here *α*_*S*_ and *α*_*I*_ are defined as in **Eqs. 2.2.6**. The coupled first order nonlinear ODEs given in **Eqs. 2.2.9-11** completely characterize the dynamics of fully competitive inhibition scheme over (P, Q, X, Y, τ) space. Here the initial conditions are (S, I, E, X, Y, P, Q, V, U) = (1, 1, 1, 0, 0, 0, 0, 0, 0) at τ = 0. When *τ* → ∞, then the reaction trajectory ends at (S, I, E, X, Y, P, Q, V, U) = (0, 0, 1, 0, 0, 1, 1, 0, 0). The steady state with respect to X occurs at 0 < *τ*_*CS*_ < ∞ where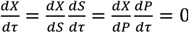. Since S and P are a monotonically decreasing and increasing functions of *τ* so that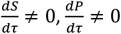 throughout the reaction timescale except at *τ*→ 0 where 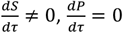 and at *τ*→ ∞ where 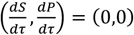, one implicitly finds at the steady state that 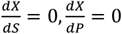. Using the same arguments, one can show that 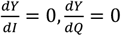 at the steady state with respect to Y at 0 < *τ*_*CI*_ < ∞ since I and Q are monotonically decreasing and increasing functions of *τ* so that 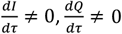 throughout the entire timescale regime except at τ = 0 where 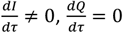 and at *τ*→ ∞ where 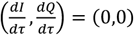. Here (*τ*_*CS*_, *τ*_*CL*_) = *k*_2_ (*t*_*CS*_, *t*_*CI*_). When *τ*_*CS*_ = *τ*_*CL*_ =*τ*_*C*_, then we represent the common steady state values of the dynamical variables as (S_C_, I_C_, E_C_, X_C_, Y_C_, P_C_, Q_C_, V_C_, U_C_).

In **Eqs. 2.2.7-9**, V and U are the dimensionless reaction velocities associated with the substrate and inhibitor conversions into their respective products (P, Q) and the mass conservation laws can be rewritten in the dimensionless velocity-substrate-product spaces as 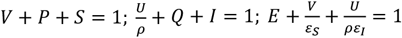. Further, the transformation rules for the reaction velocities (V, U) are 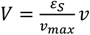 and 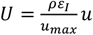. Numerically integrated sample trajectories of **Eqs. 2.2.7-9** are shown in **Fig. 2A-E**. Clearly, all the reaction trajectories in the (V, P, S) space fall on the plane V + P + S = 1, and all the reaction trajectories in the (U, Q, I) space fall on the plane 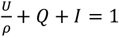 as demonstrated in **Figs. 2B** and **2C**. Parameters associated with the nonlinear system of **Eqs. 2.2.7-9** as defined in **Eqs. 2.2.2-6** can be grouped into ordinary and singular ones. Here (*ε*_*S*_, *κ*_*S*_, *ε*_*I*_, *κ*_*I*_) are the ordinary perturbation parameters. Further, (*η*_*S*_, *η*_*I*_, *ρ*) are the singular perturbation parameters since they multiply or divide the highest derivative terms. Particularly, (*η*_*S*_, *η*_*I*_) decide how fast the system of **Eqs. 2.2.7-9** attains the steady state, (*κ*_*S*_, *κ*_*I*_) decide how fast the enzyme-substrate / inhibitor complexes dissociate and (*μ*_*S*_, *μ*_*I*_) are the dimensionless Michaelis-Menten type constants which describe the summary of the effects of (*η*_*S*_, *κ*_*S*_, *η*_*I*_, *κ*_*I*_). The relative fastness of the conversion of substrate and inhibitor into their respective products as given in **Scheme A** of **Figs. 1** can be characterized by the following critical ratios of the reactions rates.

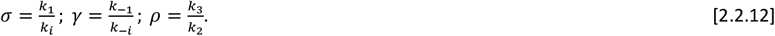

**FIGURE 2.**
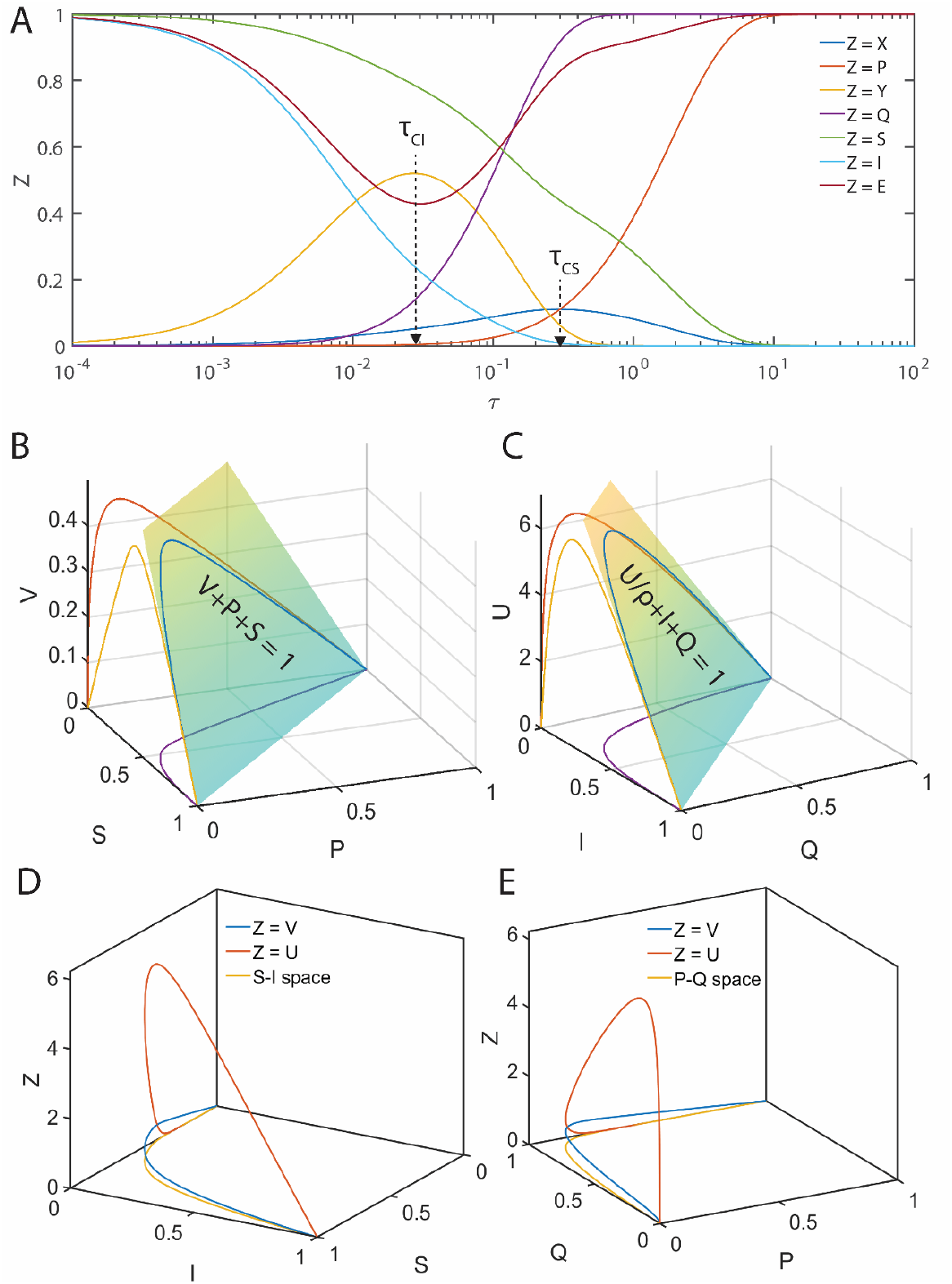
Occurrence of distinct steady state timescales with respect to enzyme-substrate (X) and enzyme-inhibitor (Y) complexes. Here S, I, E, X, Y, P, Q are the dimensionless concentrations of substrate, inhibitor, enzyme, enzyme-substrate, enzyme-inhibitor, product of substrate and product of inhibitor. Trajectories are from numerical integration of **Eqs. 2.2.7-9** with the parameters *η*_*S*_ = 0.2, *ε*_*S*_ = 4.1, *κ*_*S*_ = 3.1, *ρ* = 10, *η*_*I*_ = 0.1, *ε*_*I*_ = 1.2, *κ*_*I*_ = 0.1 along with the initial conditions (*S, I, E, X, Y, P, Q*) = (1,1,1,0,0,0,0) at τ = 0. Further, upon fixing ρ, one finds that *δ* = 0.05, *γ* = 1.55 and *σ* = 0.17. Here V = *ε*_*S*_*X* and U = *ρε*_*I*_ *Y* are the dimensionless reaction velocities corresponding to the conversion of the substrate and inhibitor into their respective products P and Q. **A**. The steady states corresponding to the enzyme-inhibitor and enzyme-substrate complexes occur at τ_CI_ = 0.03, τ_CS_ = 0.31 respectively. We should note that τ_CI_ is the time at which 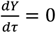 and τ_CS_ is the time at which 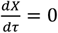. Since τ_CS_ ≠ τ_CI_ with the current parameter settings, one cannot obtain a common steady state solution to **Eqs. 2.2.7-9. B**. All the trajectories in the velocity-substrate-product (VPS) space fall within the plane V + P + S = 1. **C**. All the trajectories in the velocity-inhibitor-product (UQI) space fall within the plane U/ρ + Q + I = 1. **D**. Sample trajectories in the velocities-inhibitor-substrate (VIS, UIS) and velocities-products spaces (U, P, Q) and (V, P, Q).

When (*σ, γ, ρ*) = (1,1,1), then the dynamical aspects of the enzyme-substrate and enzyme-inhibitor complexes will be similar. Here one should note that the parameters (*η*_*S*_, *η*_*I*_, *ε*_*S*_, *ε*_*I*_, *ρ, σ*) are connected via 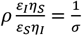 so that one finds the connection 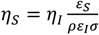. Similarly, the set of arameters (*κ*_*S*_, *κ*_*I*_, *ε*_*S*_, *ε*_*I*_, *γ, σ*) are connected via 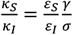 so that 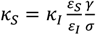. In general, the parameters (*σ, γ ρ*) are connected as follows.

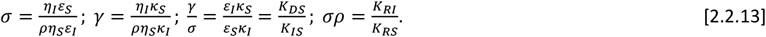

When the parameters (*η*_*S*_, *η*_*I*_, *ε*_*S*_, *ε*_*I*_, *κ*_*S*_, *κ*_*I*_) are varied independently, then fixing one parameter in (*σ, γ, ρ*) eventually fixes the other two parameters. For example, when we fix *σ* = *σ*_*f*_ then the corresponding 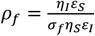 and 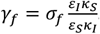. The fully competitive enzyme kinetics scheme can exhibit a complex behavior depending on the relative values of the parameters (*σ, γ, ρ*).

### 2.3. Variable transformations

Using the substitutions 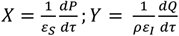 and noting the relations that *V* = *ε*_*S*_*X* and *U* = *ρε*_*I*_ *Y*, the system of **Eqs. 2.2.7-9** can be reduced to the following set of coupled nonlinear second order ODEs in the (P, Q, τ) and first order ODEs in the (V, P, Q) and (U, P, Q) spaces [37].

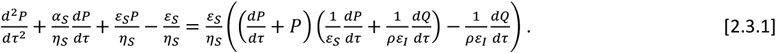

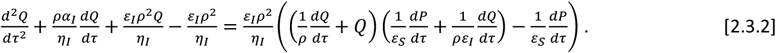

Here the initial conditions are 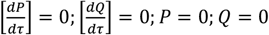.

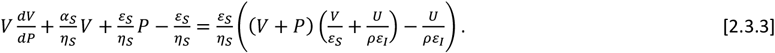

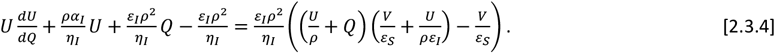

Here the initial conditions are *V* = 0; *U* = 0 at P = 0 and Q = 0. When *ρ* ≠ 1, *η*_*S*_ ≠ *η*_*I*_ and *ε*_*S*_ ≠ *ε*_*I*_, then the system of **Eqs. 2.2.7-9** will have distinct and temporally well separated steady states corresponding to the enzyme-substrate and enzyme-inhibitor complexes. Under such conditions, the system of equations given in **Eqs. 2.2.7-9** will not have common steady state solutions both in the (V, S, I) and (U, S, I) spaces (as demonstrated in **Fig. 2A**) as given by most of the currently proposed standard QSSAs.

### 2.4. Standard quasi steady state solutions

**Case I**: When (*η*_*S*_, *η*_*I*_) → (0,0) simultaneously on **Eqs. 2.2.7-8**, then upon noting the fact that *V* = *ε*_*S*_*X* and *U* = *ρε*_*I*_ *Y* one can obtain the following set of well-known quasi steady state velocity equations in the (V, S, I) and (U, S, I) spaces.

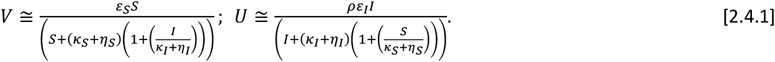

Particularly, these equations are the approximations of the post steady state dynamics of the competitive inhibition **scheme A** in the (V, S, I) and (U, S, I) spaces. When (*ε*_*S*_, *ε*_*I*_) → (0,0) along with (*η*_*S*_, *η*_*I*_) → (0,0), then one finds that (*V, U*) ≅ (0,0) along with (*P, Q*) ≅ (0,0) in the pre-steady state regime. This results in the ***reactants stationary assumption*** where we set S ≅1 and I ≅ 1 in **Eqs. 2.4.1** and the quasi-steady state velocities become as follows.

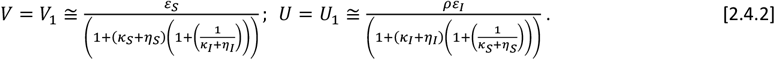

We denote the approximations given in **Eqs. 2.4.2** as V_1_ and U_1_. In terms of the original velocity variables (v, u), **Eqs. 2.4.2** can be written as follows.

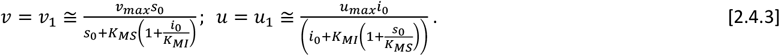

**Eqs. 2.4.3** are generally used to obtain the enzyme kinetic parameters such as (K_MS_, K_MI_, v_max_, u_max_) from the steady state based fully competitive inhibition experiments via reciprocal plotting methods under the assumptions that (*ε*_*S*_, *ε*_*I*_) → (0,0) and *ρ* = 1. Similarly, when the conditions (*η*_*S*_, *η*_*I*_) → (0,0) applied on **Eqs. 2.2.10-11**, one can arrive at the following quasi steady state velocities in the (V, P, Q) and (U, P, Q) spaces.

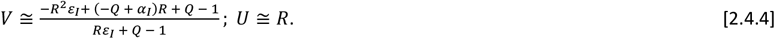

In these equations, R is the appropriate real root of the cubic equation *aR*^3^ + *bR*^2^ + *cR* + *d* = 0 where the coefficients are defined as follows.

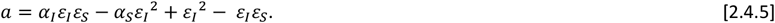

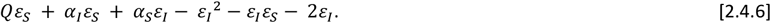

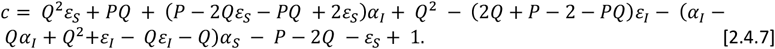

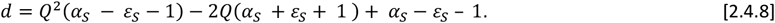

**Case II**: (*η*_*S*_, *η*_*I*_, *Q*) → (0,0,0). When only *Q* ≅ 0 which can be achieved by setting *ε*_*I*_ → 0 in the pre-steady state regime along with the conditions that (*η*_*S*_, *η*_*I*_) → (0,0), then **Eqs. 2.2.10-11** can be approximated in the (X, P, S) and (Y, P, S) spaces as follows.

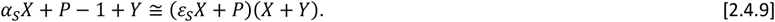

Upon the substitution of *P* = 1 − *εS X* − *S* in this equation,

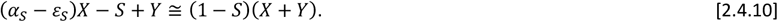

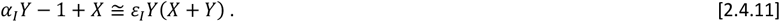

**Eq. 2.4.10** can be derived from **Eq. 2.4.9**, by using the conservation relationship V + P + S = 1 where *V* = *ε*_*S*_*X*. Upon solving **Eqs. 2.4.10-11** for (X, Y) and then converting X into V using **Eqs. 2.2.9**, one finds the following expressions for the post-steady state reaction velocity in the (V, S) space under the conditions that (*η*_*S*_, *η*_*I*_, *Q*) → (0,0,0).

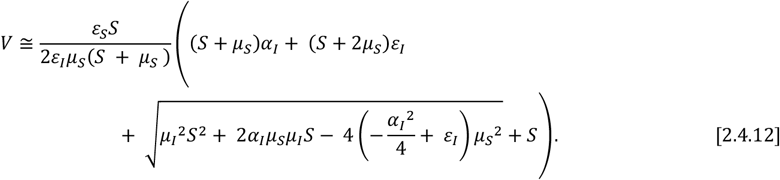

Noting that V + P + S = 1, one can express P as function of S, using P = 1 – V – S where V is defined as in **Eqs. 2.4.12**. These two equations parametrically express the post-steady state dynamics of the fully competitive inhibition scheme in the (V, P, S) space where *S* ∈ [0,1] acts as the parameter. When *P* ≅ 0 in the pre-steady state regime which can be achieved by setting *ε*_*S*_ → 0 along with the conditions that (*η*_*S*_, *η*_*I*_) → (0,0), then **Eqs. 2.2.7-8** can be approximated in the (X, Q, I) and (Y, Q, I) spaces as follows.

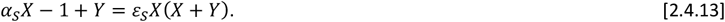

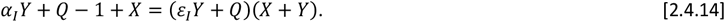

Upon the substitution of *Q* = 1 − *ε*_*I*_*Y* − *I* in this equation,

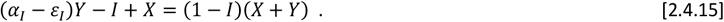

**Eq. 2.4.15** can be derived from **Eq. 2.4.14**, by using the conservation relationship U/ρ + Q + I = 1 where *U* = *ρε*_*I*_ *Y*. Upon solving **Eqs. 2.4.13-15** for (X, Y) and then converting Y into U, one finds the following expressions for the post-steady state reaction velocity in the (U, I) space under the conditions that (*η*_*S*_, *η*_*I*_, *P*) → (0,0,0).

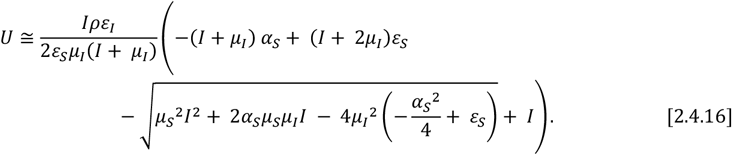

Noting that U/ρ + Q + I = 1, one can express Q as function of I, using Q = 1 – U/ρ – I where U is defined as in **Eqs. 2.4.16**. These two equations parametrically express the post-steady state dynamics in the (U, Q, I) space where *I* ∈ [0,1] acts as the parameter.

**Case III**. When (*η*_*S*_, *η*_*I*_, *ε*_*S*_, *ε*_*I*_) → 0, then one finds that *S* ≅ 1 − *P, I* ≅ 1 − *Q* and (*V, U*) ≅ (0,0) in the pre-steady state regime from which one can derive the following refined form of sQSSA approximations from **Eqs. 2.2.7-8**. Firstly, by setting (*η*_*S*_, *η*_*I*_, *ε*_*S*_, *ε*_*I*_) → (0,0,0,0) in **Eqs. 2.2.7-8**, one obtains the following set of equations.

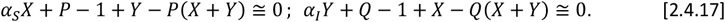

Upon solving this system of equations for (X, Y) and then transforming them into the respective velocities (V, U) using **Eq. 2.2.9**, one obtains the following post-steady state approximations in the (V, P, Q) and (U, P, Q) spaces.

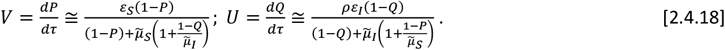

Similarly, upon the substitution of *S* ≅ 1 − *P* and *I* ≅ 1 − *Q*, **Eqs. 2.4.18** can be rewritten in the (V, S, I) and (U, S, I) spaces as follows.

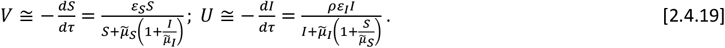

Here 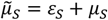. **Eqs. 2.4.17-19** are similar to **Eqs. 2.4.1** where *μ*_*S*_ and *μ*_*I*_ are replaced with 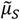 and 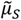. Upon applying the stationary reactant assumptions (S, I) = (1, 1) on **Eqs. 2.4.19** one obtains the **refined** form of sQSSAs. We will show in the later section that this refined form of sQSSAs can accurately predict the reaction velocities (V, U) over wide parameter ranges. Upon dividing the expression of V by the expression of U in **Eqs. 2.4.18-19**, one can obtain the following differential equation corresponding to the (P, Q) and (S, I) spaces under the conditions that (*η*_*S*_, *η*_*I*_, *ε*_*S*_, *ε*_*I*_) → (0,0,0,0).

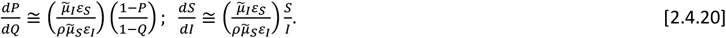

In **Eqs. 2.4.20** which are valid only in the post-steady state regimes, the initial condition in the (P, Q) space will be *P* = 0 at *Q* = 0. Similarly, the initial condition in the (S, I) space will be *S* = 1 at *I* = 1. We define *f*_*S*_ = *v*_*max*_⁄(*K*_*MS*_ + *e*_0_) and *f*_*I*_ = *u*_*max*_⁄(*K*_*MI*_ + *e*_0_) as the acceleration factors with respect to the conversion dynamics of substrate and inhibitor into their respective products (P, Q). Now let us define the critical control parameter δ as follows.

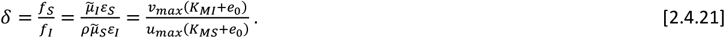

Upon solving **Eqs. 2.4.20** with the given initial conditions and using the definition of δ, one obtains the following integral solutions in the (S, I) and (P, Q) spaces [38, 39].

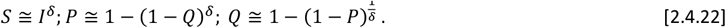

Here *δ* is the critical parameter which measures the relative speed by which enzyme-substrate and enzyme-inhibitor complexes attain their steady state. The expression for *δ* given by **Eq. 2.4.21** is a refined one compared to those definitions given in Refs. [38, 39] and straightforwardly one can show that 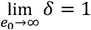. The expression for V in **Eqs. 2.4.19** in terms of S and I along with the expression for 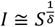 from **Eqs. 2.4.22** parametrically describe the post steady state dynamics of fully competitive enzyme kinetics in the (V, S, I) space where *S* ∈ [0,1] acts as the parameter. Similarly, expression for V in terms of P and Q as given in **Eqs. 2.4.18** along with the expression for Q that is given in **Eqs. 2.4.22** parametrically describe the post steady state dynamics in (V, P, Q) space where *P* ∈ [0,1] acts as the parameter. Upon substituting the expression for Q in terms of P obtained from **Eqs. 2.4.22** into the right-hand side of **Eqs. 2.4.18** and noting that *S* ≅ (1 − *P*) and *I* ≅ (1 − *Q*) when (*ε*_*S*_, *ε*_*I*_) → 0, so that 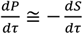 and 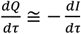, one can obtain the following approximate differential equations corresponding to the (S, τ) and (I, τ) spaces under the conditions that (*η*_*S*_, *η*_*I*_, *ε*_*S*_, *ε*_*I*_) → 0.

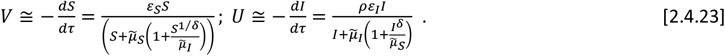

Solutions to the variable separated ODEs given in **Eqs. 2.4.23** for the initial conditions (S, I) = (1, 1) at *τ* = 0 in the (S, τ) and (I, τ) spaces can be implicitly written as follows.

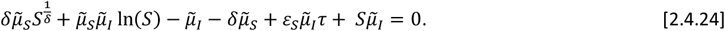

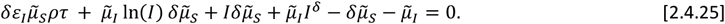

When S < 1, then one finds that 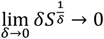 and 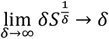and the nonlinear algebraic equation **Eq. 2.4.24** can be inverted for *S* under various limiting conditions of δ as follows.

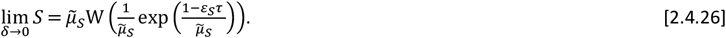

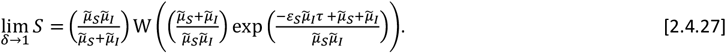

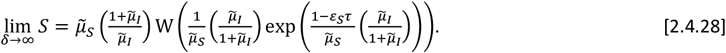

Here **W(Z)** is the **Lambert W** function which is the solution of W exp(W) = Z for W [40-42]. Similarly, when I < 1 then one finds that that 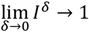and 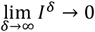 and the evolution of inhibitor level with time can be derived from **Eq. 2.4.25** under various values of δ as follows.

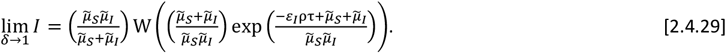

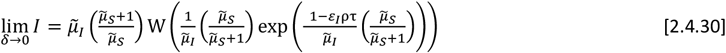

When *δ* → ∞, then *I* → 1 and one finds the following approximate asymptotic expression.

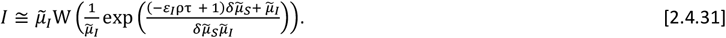

Expressions similar to **Eqs. 2.4.23-31** were proposed earlier to obtain the kinetic parameters from the substrate depletion curves of the fully competitive inhibition scheme [39]. **Eqs. 2.4.23-31** are valid only under the conditions that (*η*_*S*_, *η*_*I*_, *ε*_*S*_, *ε*_*I*_) → (0,0,0,0). In such scenarios, the right-hand sides of V and U in **Eqs. 2.4.23** can be expanded around *δ* ≅ 1 in a Taylor series as follows.

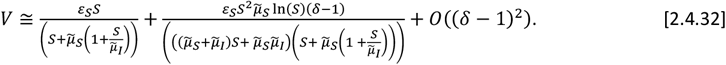

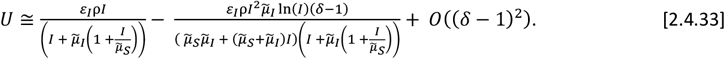

Clearly, **Eqs. 2.4.32-33** reduce to the sQSSA forms given in **Eqs. 2.4.1** only when *δ* → 1. When *δ* ≠ 1, then the enzyme-substrate-inhibitor system will exhibit complex dynamics with multiple steady states. This results in an enormous amount error in the sQSSA based parameter estimations from the experimental datasets.

**Case IV**: When (*η*_*S*_, *η*_*I*_, *P, Q*) → (0,0,0,0) so that *S* ≅ (1 − *ε*_*S*_*X*), *I* ≅ (1 − *ε*_*I*_ *Y*) in the pre-steady state regime of **Eqs. 2.2.7-8**, then one can arrive at the total quasi steady state approximations (tQSSAs) [43, 44]. We will derive explicit expressions for tQSSA in the later sections. From **Eqs. 2.4.23**, we can conclude that these are the approximate trajectories of fully competitive enzyme kinetic systems over post-steady state regime in the (V, S, I) and (U, S, I) spaces strictly either under the conditions (*ε*_*S*_, *ε*_*I*_) → (0,0), *η*_*S*_ = *η*_*I*_, *κ*_*S*_ = *κ*_*I*_and *ρ* = 1 apart from the requirement that (*η*_*S*_, *η*_*I*_, *ε*_*S*_, *ε*_*I*_) → (0,0,0,0).

#### 2.4.1. Exact steady state solutions

When the steady state timescales associated with the enzyme-substrate and enzyme-inhibitor complexes are different from each other, then **Eqs. 2.2.7-9** will not have a common steady state solution with respect to both the enzyme-substrate and enzyme-inhibitor complexes. In such scenarios, one can derive exact steady state velocities as follows. Let us assume that the steady state in the (V, P, S) space occurs at τ_CP_ where (V, S, P, U, I, Q) = (V_CP_, S_CP_, P_CP_, U_CP_, I_CP_, Q_CP_) and in the (U, Q, I) space it occurs at τ_CQ_ where (V, S, P, U, I, Q) = (V_CQ_, S_CQ_, P_CQ_, U_CQ_, I_CQ_, Q_CQ_). Noting the fact that at τ_CP_,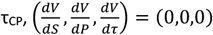 and 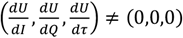 (**Figs. 3A-B**) and one can derive the following expression from **Eq. 2.3.3** by setting 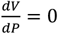 and using the conservation laws.

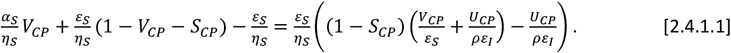

**FIGURE 3.**
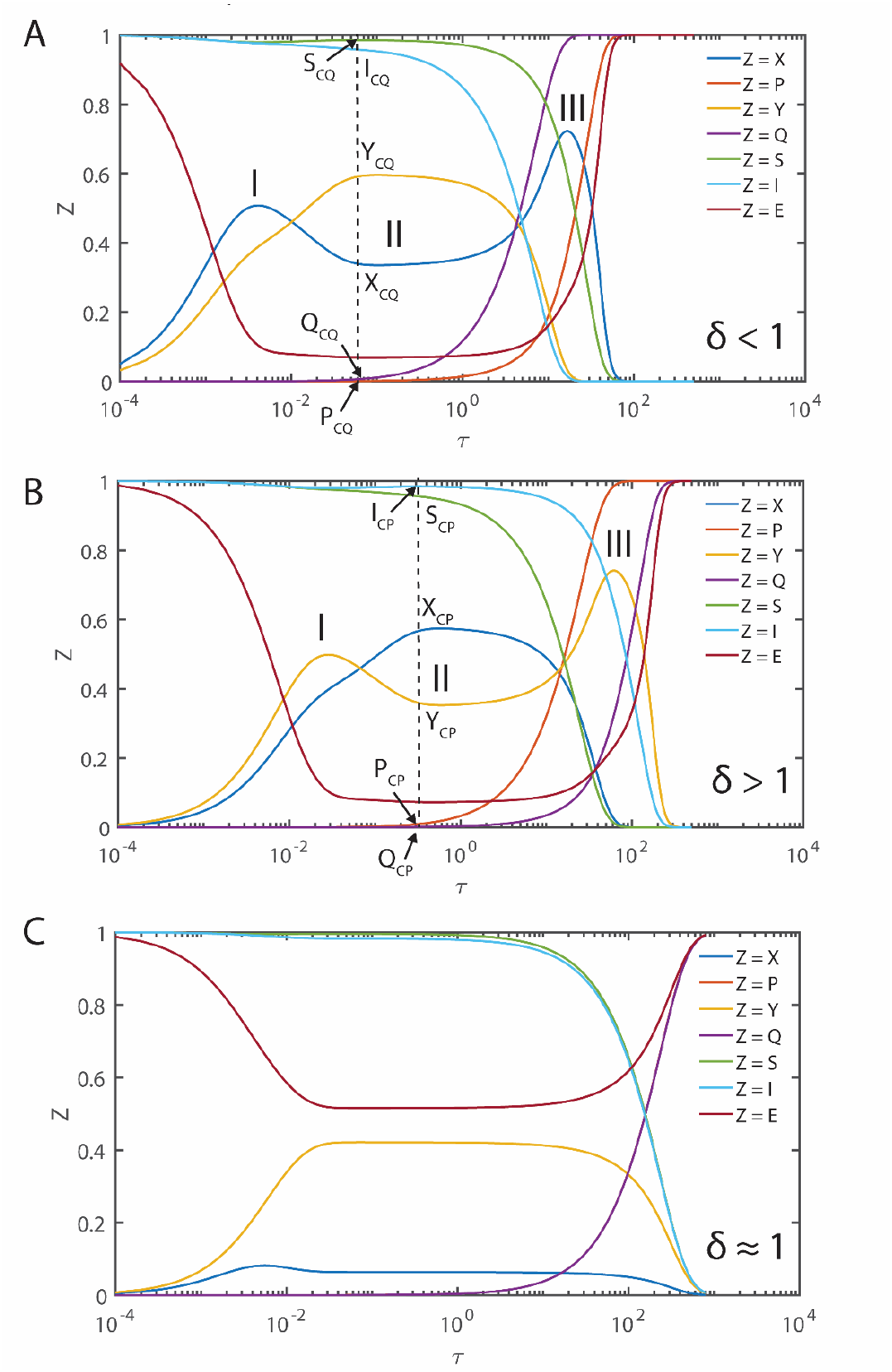
Trajectories of the enzyme kinetics with fully competitive inhibition at different values of δ. The initial conditions for the simulation of **Eqs. 2.2.7-9** are set as (*S, I, E, X, Y, P, Q*) = (1,1,1,0,0,0,0) at τ = 0. **A**. Here the settings are *η*_*S*_ = 0.002, *ε*_*S*_ = 0.04, *κ*_*S*_ = 0.2, *η*_*I*_ = 0.01, *ε*_*I*_ = 0.06, *κ*_*I*_ = 0.1 and ρ = 3.333, σ = 1, δ = 0.1405, ϒ = 3. When δ < 1 and the steady state timescale of the enzyme-substrate complex is lower than the enzyme-inhibitor complex i.e., τ_CS_ < τ_CI_, then the evolution of enzyme-substrate complex shows a bimodal type curve with respect to time. Particularly, when σ = 1 and δ > 1 or δ < 1, the temporal evolution of the enzyme-substrate and enzyme-inhibitor complexes show a complex behavior with multiple steady states. Single steady state with respect to Y occurs at (Y_CQ_, Q_CQ_, I_CQ_) and the corresponding non-steady state values in (V, P, S) space are (V_CQ_, P_CQ_, S_CQ_). **B**. Here the simulation settings are *η*_*S*_ = 0.02, *ε*_*S*_ = 0.06, *κ*_*S*_ = 0.1, *η*_*I*_ = 0.003, *ε*_*I*_ = 0.04, *κ*_*I*_ = 0.2 and ρ = 0.225, σ = 1, δ = 9, ϒ = 0.33. When δ > 1 and the steady state timescales of enzyme-substrate complex is higher than the enzyme-inhibitor complex i.e. τ_CS_ > τ_CI_, then the evolution of enzyme-inhibitor complex shows a bimodal type curve with respect to time. The single steady state with respect to X occurs at (X_CP_, S_CP_, P_CP_) and the corresponding non-steady state values in the (U, I, Q) space are (U_CP_, I_CP_, Q_CP_). **C**. Here the simulation settings are *η*_*S*_ = 0.02, *ε*_*S*_ = 0.06, *κ*_*S*_ = 8.1, *η*_*I*_ = 0.003, *ε*_*I*_ = 0.04, *κ*_*I*_ = 1.2 and ρ = 0.225, σ = 1, δ = 1.013, ϒ = 4.5.

Upon solving this equation for V_CP_, one finds the following expression.

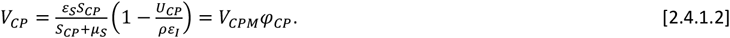

Here (V_CP_, S_CP_, P_CP_) are the steady state values in the (V, P, S) space at τ_CP_. In **Eq. 2.4.1.2**, 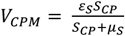 is the standard Michaelis-Menten type velocity term and 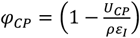 is the inhibitor dependent modifying factor. The corresponding non-steady state values in the (U, Q, I) space are (U_CP_, Q_CP_, I_CP_). Similarly, one obtains the following steady state equation from **Eq. 2.3.4** by setting 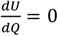 and using the conservation laws of (U, Q, I) space for the enzyme-inhibitor complex at the time point τ_CQ_ where 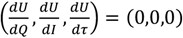 and 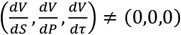.

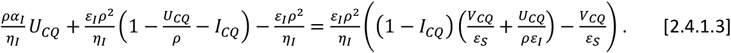

Upon solving this equation for U_CQ_, one finds the following expression.

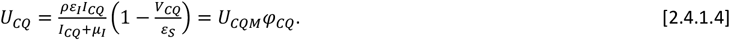

Here (U_CQ_, I_CQ_, Q_CQ_) are the corresponding steady state values in the (U, Q, I) space with respect to the enzyme-inhibitor complex at τ_CQ_ and (S_CQ_, P_CQ_, V_CQ_) are the corresponding non-steady state values in the (V, P, S) space. In **Eq. 2.4.1.4**, 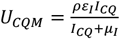 is the standard Michaelis-Menten type velocity term and 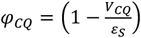 is the substrate dependent modifying factor. However, to find V_CP_ and U_CQ_ which are the exact steady state velocities, one needs to know S_CP_, U_CP_, I_CQ_ and V_CQ_. When the steady state timescales corresponding to the enzyme-substrate and enzyme-inhibitor complexes are the same, then S_CP_ = S_CQ_, U_CP_ = U_CQ_, I_CQ_ = I_CP_ and V_CQ_ **=** V_CP_ as shown in **Fig. 3C** and subsequently **Eqs. 2.4.1.1-4** reduce to the standard QSSA **Eqs. 2.4.1**.

#### 2.4.2. Complexity of the steady states

When *δ* = 1, then the approximate post-steady state reaction velocities under the conditions that (*η*_*S*_, *η*_*I*_, *ε*_*S*_, *ε*_*I*_) → (0,0,0,0) can be given by **Eqs. 2.4.23** which are monotonically increasing (and decreasing) functions of S and I. Approximate steady state velocities (V, U) can be obtained from **Eqs. 2.4.23** by asymptotic extrapolation as (*S, I*) → (1,1) which is the stationary reactant assumption. When *δ* ≠ 1, then **Eqs. 2.4.23** will exhibit a turn over type behavior upon increasing (*S, I*) from (0,0) towards (1,1) with extremum points at which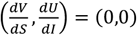. This dynamical behavior is demonstrated in **Figs. 3**. This means that there are at least two different points at which 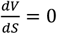 or 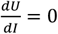 within (0,0) < (S, I) < (1, 1) depending on the value of *δ* and sQSSAs given by **Eqs. 2.4.23** are valid only when *δ* = 1. When *δ* < 1, then the reaction velocity associated with the enzyme-substrate complex will show two different steady state regions at which 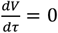(so that 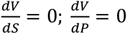 in the (V, S) and (V, P) spaces respectively) as demonstrated in **Fig. 3A**. The velocity expressions given in **Eqs. 2.4.23** with (*S, I*) → (1,1) approximately represent the first transient steady state in the (V, S) space. The approximate prolonged secondary steady state corresponding to the enzyme-substrate dynamics can be obtained by solving 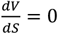 for S where V is given as in **Eqs. 2.4.23** as follows.

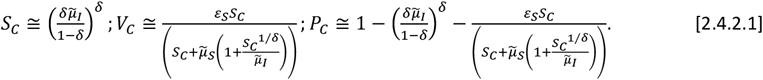

One can obtain P_C_ using the conservation relationship V_C_ + S_C_ + P_C_ = 1. When *δ* > 1, then the reaction velocity associated with the enzyme-inhibitor complex will show two different steady state regions at which 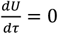 (so that 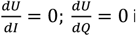 in the (U, I) and (U, Q) spaces respectively) as demonstrated in **Fig. 3B**. The velocity expressions given in **Eqs. 2.4.23** with (*S, I*) → (1,1) approximately represent the first transient steady state in the (U, I) space. One can obtain the approximate prolonged secondary steady state velocity corresponding to the enzyme-inhibitor complex U_C_ and the inhibitor concentration I_C_ by solving 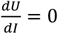 for I where U is given as in **Eqs. 2.4.23** as follows.

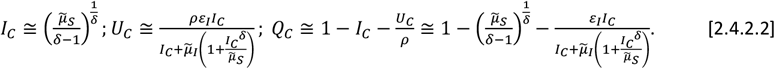

One can obtain Q_C_ using the conservation relationship U_C_/ρ + Q_C_ + I_C_ = 1. Here one should note that **Eqs. 2.4.2.1** will not be valid when *δ* ≥ 1 and **Eqs. 2.4.2.2** will not be valid when *δ* ≤ 1 since the steady state values can be negative or complex under such conditions. A common steady state can occur only when *δ* = 1 as demonstrated in **Fig. 3C** and **Eqs. 2.4.23**. Remarkably, for the first time in the literature we report this phenomenon and none of the earlier studies on the fully and partial competitive inhibition captured this complex dynamical behavior.

### 2.5. Solutions under coupled and uncoupled conditions

For the general case, the approximate steady state timescales corresponding to the enzyme-substrate and enzyme-inhibitor complexes can be obtained as follows. Using the scaling transformations 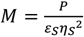 and 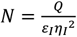, **Eqs. 2.3.1-2** can be rewritten in the following form of Murugan equations [10, 37] with appropriate initial conditions.

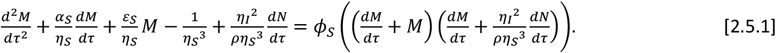

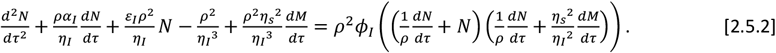

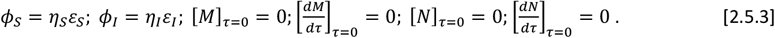

Here *ϕ*_*S*_ and *ϕ*_*I*_ are the ordinary perturbation parameters which multiply the nonlinear terms. **Eqs. 2.5.1-2** along with the initial conditions given in **Eqs. 2.5.3** completely characterize the dynamical aspects of the fully competitive enzyme inhibition scheme. **Eqs. 2.5.1-3** are the central equations of this paper from which we will derive several approximations for the pre- and post-steady state regimes under various set of conditions.

#### 2.5.1. Approximate solutions under coupled conditions

When (*ϕ*_*S*_, *ϕ*_*I*_) → (0,0), then **Eqs. 2.5.1**-**2** become coupled linear system of ordinary differential equations as follows.

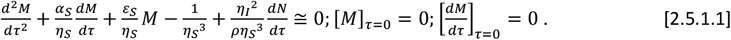

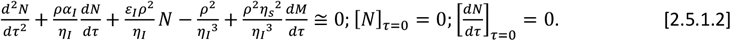

**Eqs. 2.5.1-3** are derived here for the first time in the literature. We denote **Eqs. 2.5.1-3** as Murugan type II equations [37] and the φ-approximations given by **Eqs. 2.5.1.1-2** are exactly solvable. Interestingly, these equations can be rewritten in terms of fourth order uncoupled ODEs with constant coefficients both in the (M, τ) and (N, τ) spaces as follows (see **Appendix A** for details). In the (M, τ) space, one can straightforwardly derive the following results.

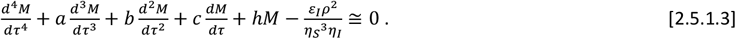

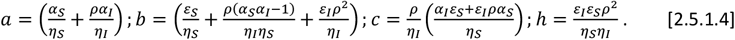

Upon obtaining the solution for the (M, τ) space, one can straightforwardly obtain the expression corresponding to the (N, τ) space as follows.

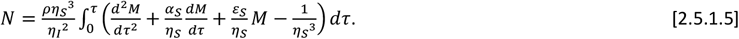

The first two initial conditions corresponding to the fourth order uncoupled ODE given by **Eqs. 2.5.1.3** can be written as follows.

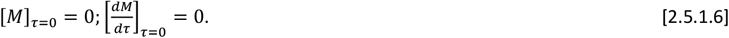

Other two initial conditions directly follow from the initial conditions corresponding to N.

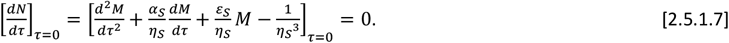

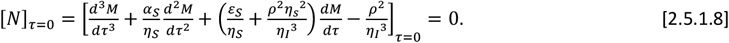

Similar to **Eq. 2.5.1.3**, one can also derive the following solution set corresponding to (N, τ) space.

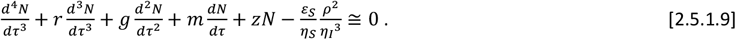

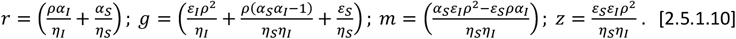

Upon obtaining the solution for the (N, τ) space, one can directly obtain the expression corresponding to the (M, τ) space as follows.

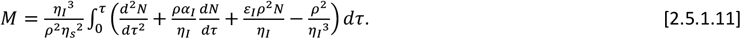

The initial conditions corresponding to the fourth order uncoupled ODEs given by **Eqs. 2.5.1.9** can be written as follows.

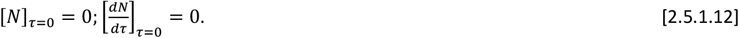

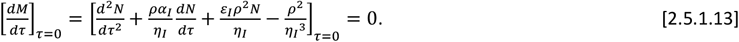

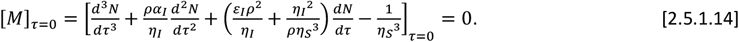

Solution to **Eqs. 2.5.1.1-2** can be obtained either by solving **Eqs. 2.5.1.3-8** or **Eqs. 2.5.1.9-14**. The detailed expressions for the solution are given in **Appendix A**. Upon obtaining solutions in the (M, τ) and (N, τ) spaces, one can revert back to (P, τ) and (Q, τ) spaces using the scaling transformations 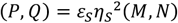 from which one can obtain the parametric expressions for the trajectories in the (V, P, S), (U, I, Q), (V, I, S), (U, I, S), (V, P, Q), (U, P, Q) and (V, U) spaces using appropriate mass conservation relationships where *τ* acts as the parameter.

#### 2.5.2. Approximate solutions under uncoupled conditions

When 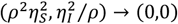 along with the conditions of ϕ-approximations as (*ϕ*_*S*_, *ϕ*_*I*_) → (0,0), then **Eqs. 2.5.1-2** can be approximated by the following uncoupled set of ODEs.

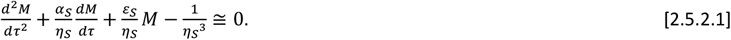

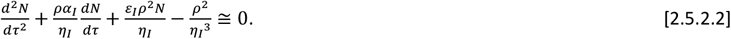

The conditions 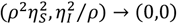 will be true when (*k*_3_⁄*k*_1_*s*_0_, *k*_2_⁄*k*_*d*_ *i*_0_) → (0,0). Here the initial conditions are 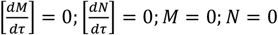 at τ = 0. Upon reverting these equations back into the (P, Q, τ) space, one obtains the following set of uncoupled ODEs along with the corresponding initial conditions.

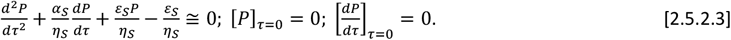

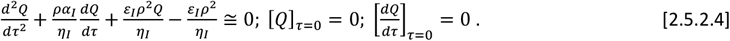

Along with the conditions that 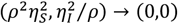, the uncoupled **Eqs. 2.5.2.3-4** are valid only (a) when (*ϕ*_*S*_, *ϕ*_*I*_) → 0 so that there is no competitive inhibition kinetics or (b) the dissociation rate constants of both the enzyme-substrate and enzyme-inhibitor complexes are high enough to uncouple the competitive kinetics i.e., (*κ*_*S*_, *κ*_*I*_) → (∞, ∞). Both these conditions will lead to the approximation *EE* = (1 − *X* − *Y*) ≅ 1. We will show later that these conditions decrease the mismatch between the steady state timescales of the enzyme-substrate and enzyme-inhibitor complexes. Upon solving these ODEs with the given initial conditions, we can derive the approximate expressions for the dynamics of the competitive inhibition scheme (P, Q, V, U, S, I) as follows [10, 37].

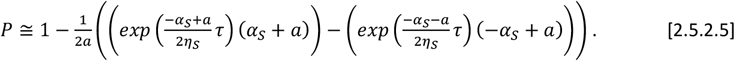

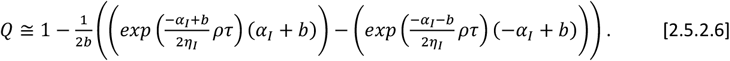

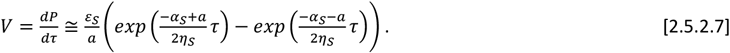

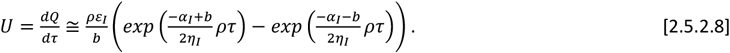

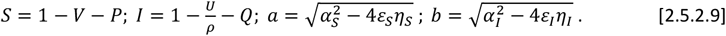

The expressions given in **Eqs. 2.5.2.5-9** for (V, P, S, τ) space will be valid only when 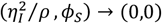 and those expressions given for (Q, U, I, τ) space will be valid only when 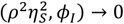. Upon solving 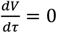 and 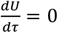 for τ in where (V, U) are given as in **Eqs. 2.5.2.7-8**, one can obtain the following approximations for the steady state timescales corresponding to substrate and inhibitor conversion dynamics. When 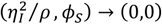, then one finds that,

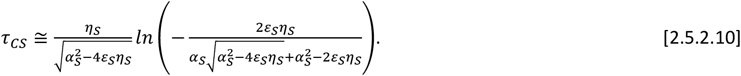

When 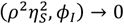, then one finds that,

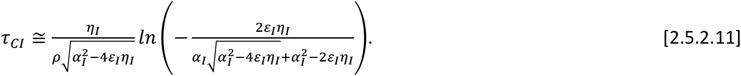

Here *τ*_*CS*_ and *τ*_*CI*_ are the approximate timescales at which the steady states with respect to the enzyme-substrate and enzyme-inhibitor complexes occur under uncoupled conditions. Upon substituting the expression for *τ*_*CS*_ into the expressions for (V, P, S) given in **Eqs. 2.5.2.5-9** one can obtain the corresponding steady state values (V_C_, P_C_, S_C_). In the same way, upon substituting the expression for *τ*_*CI*_ into the expressions for (U, Q, I) one can obtain the corresponding steady state values (U_C_, Q_C_, I_C_). Clearly, the condition *τ*_*CS*_ ≅ *τ*_*CI*_ is critical for the occurrence of a common steady state with respect to the reaction dynamics of both the enzyme-substrate and enzyme-inhibitor complexes. When *τ*_*CI*_ > *τ*_*CS*_, then the substrate depletion with respect to time will show a typical non-monotonic trend since the inhibitor reverses the enzyme-substrate complex formed before time *τ*_*CI*_ in the pre-steady state regime. In the same way, when *τ*_*CI*_ < *τ*_*CS*_ then the inhibitor depletion will show a non-monotonic behavior since the substrate reverses the enzyme-inhibitor complex formed before time *τ*_*CS*_ in the pre-steady state regime. These phenomena eventually introduce significant amount of error in various QSSAs. Under uncoupled conditions i.e., when 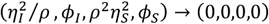, then one can rewrite the uncoupled approximations given in **Eqs. 2.5.2.3-4** over (V, P) and (U, Q) spaces as follows.

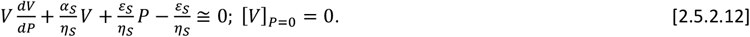

Noting that 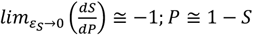, one can obtain the ODE corresponding to the (V, S) space as follows.

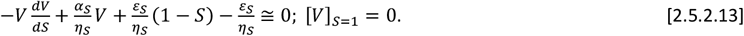

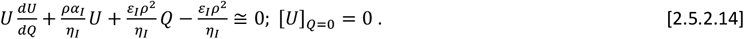

Noting that 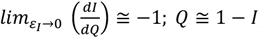, one can obtain the ODE corresponding to the (U, I) space as follows.

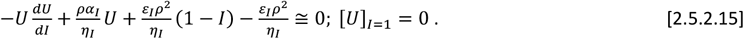

Upon considering only the linear, uncoupled portions in the (V, P) space as given in **Eqs. 2.5.2.12**, one obtains the following approximations for V and S as functions of P.

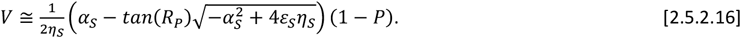

Using the conservation laws one finds the following.

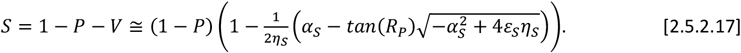

Here R_P_ is the solution of the following nonlinear algebraic equation.

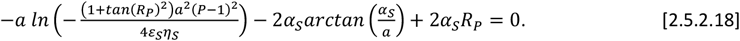

In this equation, 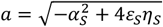. These approximate equations parametrically describe the dynamics of the enzyme catalyzed substrate conversion in the entire regime of (V, P, S) space from (V, P, S) = (0, 0, 1) to (V, P, S) = (0, 1, 0) including the steady states (V_C_, P_C_, S_C_) that occurs at 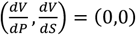. Here P acts as the parameter. Further, upon solving 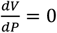 in **Eq. 2.5.2.16** one obtains the steady state concentration of the product of substrate as follows.

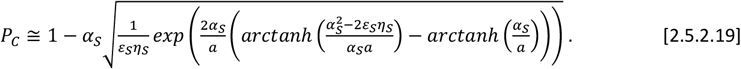

In this equation, 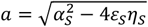. Upon substituting the expression of P_C_ into the expressions of V and S as given in **Eqs. 2.5.2.16-17**, one can obtain the steady state expressions for V_C_ and S_C_. In the same way, upon considering only the linear, uncoupled portions in the (U, Q) space as given in **Eq. 2.5.2.14**, one obtains the following approximations for U and I as functions of Q.

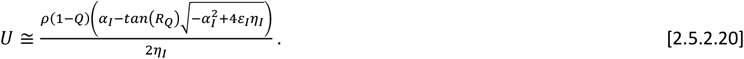

Using the conservation laws, one finds the following.

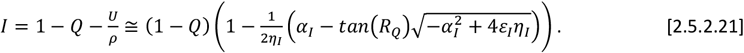

Here R_Q_ is the solution of the following nonlinear algebraic equation.

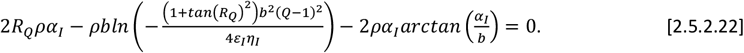

In this equation, 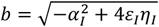. These approximate equations parametrically describe the dynamics of the enzyme catalyzed inhibitor conversion in the entire regime of (U, Q, I) space from (U, Q, I) = (0, 0, 1) to (U, Q, I) = (0, 1, 0) including the steady states (U_C_, Q_C_, I_C_) that occurs at 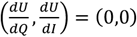. Here Q acts as the parameter. Further, upon solving 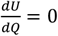 in **Eq. 2.5.2.20** one obtains the steady state level of the product of inhibitor as follows.

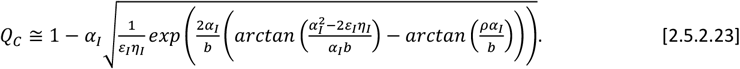

Here the term *b* is defined as in **Eq. 2.5.2.22**. Upon substituting the expression of Q_C_ for Q into the expressions of U and I as given in **Eqs. 2.5.2.20-21**, one can obtain the steady state expressions for U_C_ and I_C_.

### 2.6. Approximate pre-steady state solutions

Using the scaling transformation 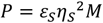 one can rewrite the set of coupled nonlinear ODEs corresponding to the fully competitive inhibition scheme given in **Eqs. 2.5.1** and **2.2.11** in the (M, Y, Q, τ) space as follows.

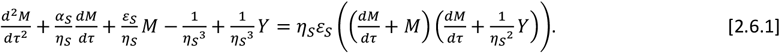

Noting that 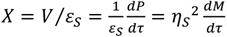, one finds from **Eq. 2.2.11** that,

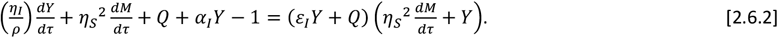

Here the initial conditions are 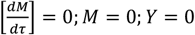. Using these equations one can derive the pre-steady state expressions associated with the enzyme-substrate complex under various conditions as follows.

**Case I**: When (*ε*_*I*_, *η*_*I*_, *ε*_*S*_) → (0,0,0), then one finds that *I* = (1 − *ε*_*I*_ *Y* − *Q*) ≅ 1 − *ε*_*I*_ *Y* since *Q* ≅ 0 in the pre-steady state regime. Under such conditions one can arrive at the approximation for Y from **Eq. 2.6.2** as 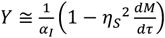. Upon substituting this expression for Y in **Eq. 2.6.1** and using the variable transformation 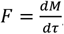, one finally arrives at the following approximate ODEs corresponding to the pre-steady state regimes in the (M, τ) and (F, M) spaces.

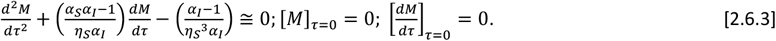

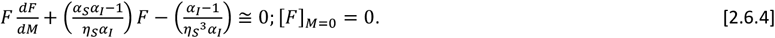

**Case II**: When (*ε*_*I*_, *η*_*I*_, *ϕ*_*S*_) → (0,0,0) in **Eqs. 2.6.1-2**, then following the same arguments as in **Eqs. 2.6.3-4**, one finds the following refined approximations in the (M, τ) and (F, M) spaces.

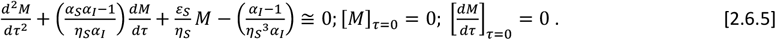

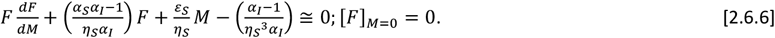

We will discuss the solutions to **Eqs. 2.6.5-6** in the later section in detail. **Eq. 2.6.3** is a linear second order ODE with constant coefficients that is exactly solvable. Upon solving the nonlinear ODE given in **Eq. 2.6.4** along with the initial condition, one arrives at the following approximate integral solution under the conditions that (*ε*_*S*_, *η*_*I*_, *ε*_*I*_) → 0 in the (F, M) space.

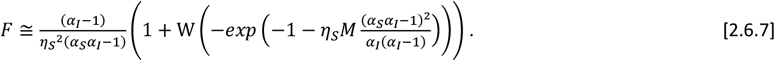

Upon solving **Eqs. 2.6.3-4** with the given initial conditions as in **Eq. 2.6.7** and then reverting back to (V, P), (V, S) and (V, τ) spaces using the transformations (*P, V*) = *ε*_*S*_ *η*_*S*_^2^(*M, F*) and using the conservation relationships, we arrive at the following approximate solutions under the conditions that (*ε*_*S*_, *η*_*I*_, *ε*_*I*_) → (0,0,0). In the (V, τ) space one finds the following result.

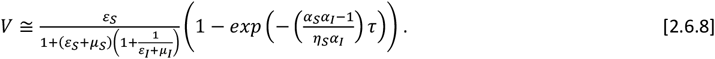

In the (V, P) space the approximate solution becomes as follows.

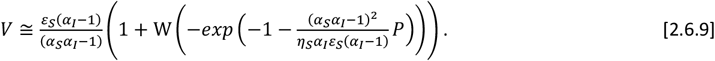

Upon solving **Eq. 2.6.4** implicitly in the (F, M) space and then reverting back to (V, P) space using the transformation (*V, P*) = *ε*_*S*_ *η*_*S*_^2^(*F, M*) and substituting P = 1 – V – S before the inversion of (**Appendix B**) the hitherto obtained implicit expression in terms of Lambert W function, one obtains the following pre-steady state solution in the (V, S) space under the conditions that (*ε*_*S*_, *η*_*I*_, *ε*_*I*_) → (0,0,0).

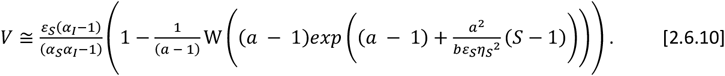

The parameters *a* and *b* in **Eq. 2.6.10** are defined as follows.

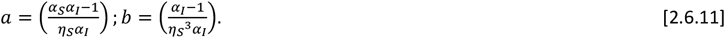

By expanding **Eq. 2.6.10** in a Taylor series around S = 1, one finds that *V* ≅ 1 − *S* + O((*S* − 1)^2^) which means that *P* ≅ 0 in the pre-steady state regime. It is also interesting to note that all the rajectories in the (V, S) space will be confined inside the triangle defined by the lines *V* ≅ 1 − *S*, V = 0 and S = 0. When P or τ becomes sufficiently large, then **Eqs. 2.6.8-9** asymptotically converge to the following limiting value that is close to the steady state reaction velocity under the conditions that (*ε*_*I*_, *η*_*I*_, *ε*_*S*_) → (0,0,0). We denote this approximation as V_2_.

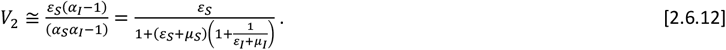

We will show in the later sections that this approximation works very well in predicting the steady state reaction velocities over wide ranges of parameters. One can also arrive at **Eq. 2.6.12** under the conditions that (*ε*_*I*_, *η*_*I*_, *ε*_*S*_, *η*_*S*_) → (0,0,0,0) as in the refined sQSSA expression given in **Eqs. 2.4.19**. In terms of original variables, this equation can be written as follows.

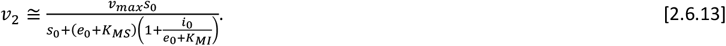

Similar to **Eqs. 2.6.1-2**, using the transformation *Q* = *ε*_*I*_ *η*_*I*_^2^*N*, one can rewrite **Eqs. 2.5.2** and **2.2.7** as the following coupled system of ODEs.

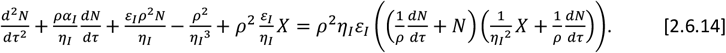

Noting 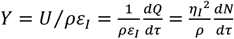, one finds from **Eq. 2.2.10** that,

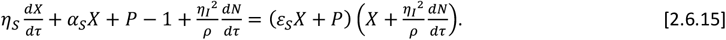

Here the initial conditions are 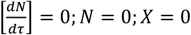 at τ = 0. Using these equations one can derive the pre-steady state expressions associated with the enzyme-inhibitor complex under various conditions.

**Case III**: When (*ε*_*S*_, *η*_*S*_, *ε*_*I*_) → (0,0,0), then *S* = (1 − *ε*_*S*_*X* − *P*) ≅ 1 − *ε*_*S*_*X* and *P* ≅ 0 in the pre-steady state regime and one finds from **Eq. 2.6.15** that 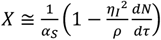. Upon substituting this expression of X into **Eq. 2.6.14**, setting (*ε*_*S*_, *η*_*S*_, *ε*_*I*_) → 0 and using the transformation 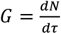 in **Eqs. 2.6.14-15** one can derive the following approximations in the (N, τ) and (G, N) spaces.

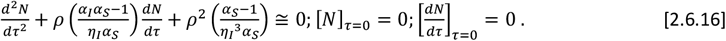

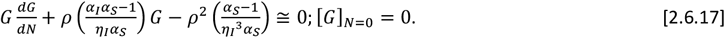

**Case IV**: When (*ε*_*S*_, *η*_*S*_, *ϕ*_*I*_) → (0,0,0), then following the same arguments with respect to **Eqs. 2.6.5-6**, one finds the following approximations in the (N, τ) and (G, N) spaces.

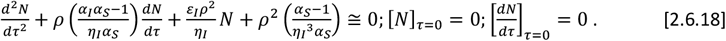

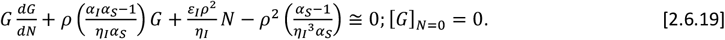

We will discuss the solutions to **Eqs. 2.6.18-19** in the later section in detail. **Eq. 2.6.16** is a second order linear ODE with constant coefficients that is exactly solvable. Upon solving the nonlinear ODE given in **Eq. 2.6.17** along with the initial condition, one can arrive at the following integral solution in the pre-steady state regime in the (G, N) space.

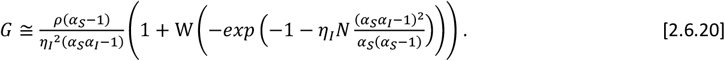

Upon solving **Eqs. 2.6.16-17** with the given initial conditions and then reverting back to the (U, Q), (U, I) and (U, τ) spaces using the transformations (*Q, U*) = *ε*_*I*_*η*_*I*_^2^(*N, G*) one finds the followingapproximate solutions to **Eqs. 2.6.16-17** under the conditions that (*ε*_*S*_, *η*_*I*_, *ε*_*I*_) → (0,0,0). In the (U, τ) space one finds the following result.

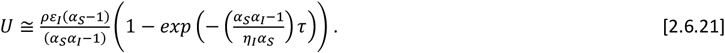

In the (U, Q) space the approximate solution becomes as follows.

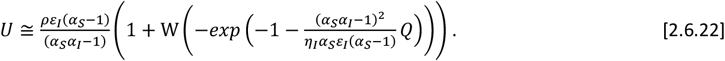

Upon solving **Eq. 2.6.17** implicitly in the (G, N) space and then reverting back to (U, Q) space using the transformations (*Q, U*) = *ε*_*I*_ *η*_*I*_^2^(*N, G*) and substituting Q = 1 – U/ρ – I before the inversion in terms of Lambert W function, one obtains the following pre-steady state solution in the (U, I) space under the conditions that (*ε*_*S*_, *η*_*I*_, *ε*_*I*_) → (0,0,0).

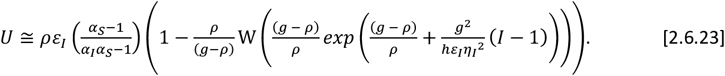

The terms *g* and *h* in **Eq. 2.6.23** are defined as follows.

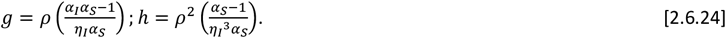

Upon expanding the right-hand side of **Eq. 2.6.23** in a Taylor series around I = 1, one finds that *U* ≅ *ρ*(1 − *I*) + O((*I* − 1)^2^) which means that *Q* ≅ 0 in the pre-steady state regime where **Eqs. 2.6.20-24** are valid. When Q or τ becomes sufficiently large, then **Eqs. 2.6.21** asymptotically converges to the following limiting value that is close to the steady state value of U under the conditions that (*ε*_*S*_, *η*_*S*_, *ε*_*I*_) → (0,0,0). We denote this approximation as U_2_. We will show in the later sections that this approximation works very well over wide ranges of parameter values.

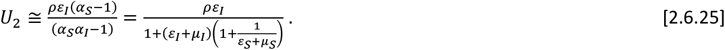

In terms of original variables, this equation can be written as follows.

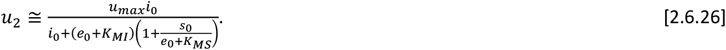

This equation is similar to the refined sQSSA given in **Eqs. 2.4.19** that was derived under the conditions that (*ε*_*I*_, *η*_*I*_, *ε*_*S*_, *η*_*S*_) → (0,0,0,0).

#### 2.6.1. Steady state timescales

Under the conditions that (*ε*_*S*_, *η*_*I*_, *ε*_*I*_) → (0,0,0) and (*ε*_*S*_, *η*_*S*_, *ε*_*I*_) → (0,0,0) as in **Eqs. 2.6.8** and **2.6.21** and noting that these approximate pre-steady state velocities (V, U) approach the steady state values as *ττ* → ∞, one finds the following approximate steady state timescales associated with the substrate and inhibitor conversion dynamics under coupled conditions.

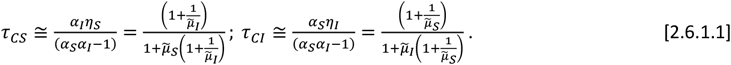

In terms of original variables, **Eqs 2.6.1.1** can be written as follows.

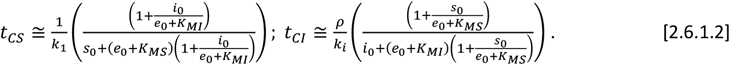

**Eqs. 2.6.1.2** are similar to the equations derived in Ref. [39] (see Eqs. 34-35 in this reference) for the steady state timescales. However, the numerator terms were set to unity for *ρ* = 1 and also e_0_ was not added with (*K*_*MS*_, *K*_*MI*_) in their expressions. When *e*_0_ → ∞, then (*t*_*CS*,_*t*_*CI*_) → (0,0) is a reasonable observation from **Eqs. 2.6.1.2**. Further, those approximate expressions suggested in Ref. [39] for *t*_*CS*_ and *t*_*CI*_ predicted that when (*i*_0_, *s*_0_) → (∞, ∞), then (*t*_*CS*_, *t*_*CI*_) → (0,0). However, when *i*_0_ → ∞, then the probability of binding of substrate with the enzyme will be decreased. As a result, when s_0_ is fixed, then *t*_*CS*_ will increase asymptotically towards a limiting value as *i*_0_ → ∞. Similarly, when *s*_0_ → ∞, then the probability of binding of inhibitor with the enzyme will be decreased towards a minimum. As a result, when i_0_ is fixed, then *t*_*CI*_ will increase asymptotically towards a limiting value as *s*_0_ → ∞. In this context, **Eqs. 2.6.1.2** correctly predict the following limiting behaviors of the steady state timescales.

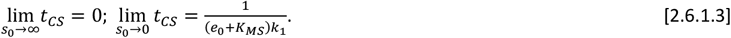

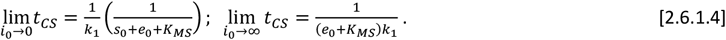

Here one should note that *s*_0_ → 0 or *i*_0_ → ∞ will have similar limiting behavior on *t*_*CS*_. This is reasonable since setting *i*_0_ → ∞ will eventually decreases the binding probability of substrate with the enzyme to a minimum. Similarly, one also finds the following limiting behaviors of *t*_*CI*_.

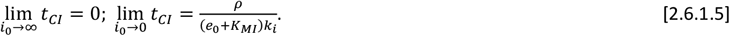

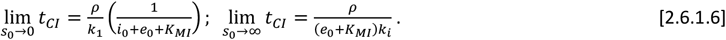

Here one should note that *i*_0_ → 0 or *s*_0_ → ∞ will have similar limiting behavior on *t*_*CI*_ since setting *s*_0_ → ∞ will eventually decreases the binding probability of inhibitor with the enzyme. **Eqs. 2.6.1.3-6** should be interpreted only in the asymptotic sense since setting (s_0_, i_0_) = (0, 0) will eventually shuts down the respective catalytic channels. **Eqs. 2.6.1.1** clearly suggest that a common steady state between enzyme-substrate and enzyme-inhibitor complexes can occur only when *α*_*I*_*η*_*S*_ ≅ *α*_*S*_ *η*_*I*_ or explicitly when the ratio 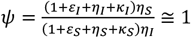. When *η*_*S*_ ≅ *η*_*I*_, then the condition *ψ* ≅ 1 can be achieved by simultaneously setting large values for any one of the parameters (*ε*_*S*_, *κ*_*S*_) in the numerator part and any one of the parameters (*ε*_*I*_, *κ*_*I*_) from the denominator part so that their ratio *ψ* tends towards one. For example, one can consider setting (*ε*_*I*_, *κ*_*S*_) → ∞ or a combination (*ε*_*S*_, *κ*_*I*_) → ∞ and so on. Under such conditions, the error in the estimation of the kinetic parameters using sQSSAs will be at minimum. In general, the condition for the minimal error in sQSSA can also be derived from *ψ* ≅ 1 in the following form.

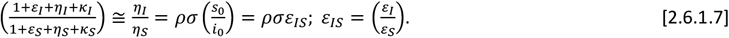

Upon considering the connection between the parameters (*ρ, γ, σ*) as given in **Eqs. 2.2.13** and noting that (*ρ, σ*) ≅ (1,1) for most of the substrate-inhibitor pairs, the required conditional **Eqs. 2.6.1.7** can be rewritten upon setting *ρσ* = 1 as follows.

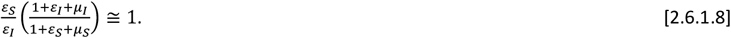

In terms of original variables this conditional equation **Eqs. 2.6.1.8** can be written as follows.

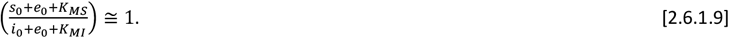

In most of the *in vitro* quasi-steady state experiments, one sets larger values for (*s*_0_, *i*_0_) than (*e*_0_, *K*_*MS*_, *K*_*MI*_) so that the left-hand side of **Eq. 2.6.1.9** tends towards unity which is essential (but not sufficient) condition to minimize the error of such QSSAs as suggested [38, 39] by most of the earlier studies in a slightly different form as *ρ* ≅ 1, *σ* ≅ 1 and *ε*_*IS*_ ≅ 1. Further, **Eqs. 2.6.1.8-9** will work only when the condition *ρσ* = 1 is true.

### 2.7. Minimization of error in the quasi steady state approximations

The essential conditions required to minimize the error in various sQSSAs of fully competitive inhibition scheme with stationary reactants assumption can be obtained as follows. The sQSSAs (**Eqs. 2.4.19**) describe only the post steady state regime in the (V, S, I) and (U, S, I) spaces and approximate the pre-steady state region with the asymptotic velocities corresponding to (S, I) → (1,1). To extract the enzyme kinetic parameter, one generally uses **Eqs. 2.4.19** with stationary reactant assumptions (S, I) = (1, 1). From pre-steady state approximations given in **Eqs. 2.6.8, 2.6.21** and the refined sQSSAs given in **Eqs. 2.4.19** with (s,I)→(1,1), one can define the error estimate in the pre-steady state regime as follows [37].

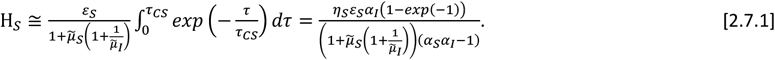

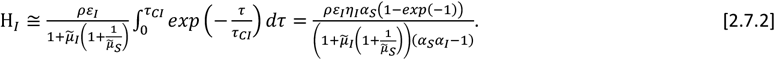

Here *τ*_*CS*_and τ_*CI*_ are defined as in **Eqs. 2.6.1.1**, *H*_*S*_ and *H*_*I*_ are the overall errors in the refined sQSSAs on enzyme-substrate and enzyme-inhibitor complexes respectively. Upon solving 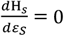 for *ε*_*S*_and 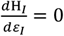 for *ε*_*I*_ after substitution of the approximate expressions for τ_*CS*_ and τ_*CI*_ from **Eqs. 2.6.1.1**, and expanding the terms 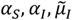 and 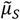, one obtains the following expressions for *ε*_*S,max*_ and *ε*_*I,max*_ at which the errors due to sQSSAs attain maxima.

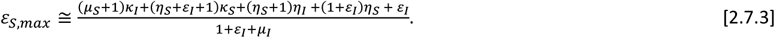

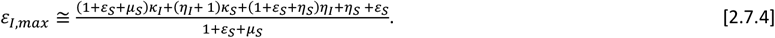

Clearly, the following generalized conditions i.e., *ε*_*S*_≪*ε*_*S,max*_and *ε*_*I*_≪*ε*_*I,max*_are essential to minimize the error in sQSSAs (refined sQSSA given in **Eqs. 2.4.19** with (s,I)→(1,1)) associated with the enzyme-substrate and enzyme-inhibitor dynamics since (*η*_*S*,_ *η*_*I*,_ *ε*_*s*,_ *ε*_*I*_)→(0,0,0,0) are the preconditions for the validity of the standard QSSAs. One can write these sufficient conditions (we denote them as **E**_**1**_ and **E**_**2**_) explicitly as follows.

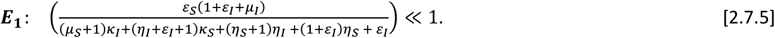

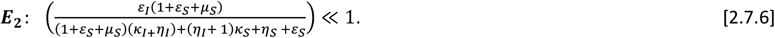

We will show later that there are strong correlations between **Eqs. 2.7.5-6** and the corresponding observed overall errors in the estimation of (V, U) using the refined sQSSA methods.

### 2.8. Approximate time dependent solutions

From **Eqs. 2.6.5-6** one can derive the refined expressions for the reaction velocity V and product P under the conditions that (*ε*_*I*_, *η*_*I*,_ *ϕ*_*S*_)→(0,0,0) as follows.

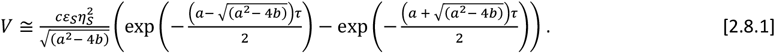

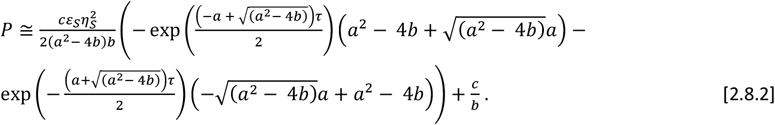

Here the parameters *a, b* and *c* are defined as follows.

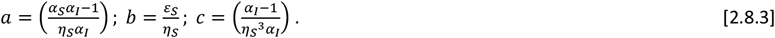

From **Eqs. 2.6.18-19** one can derive the refined expressions for the reaction velocity U and product Q under the conditions that (*ε*_*s*,_ *η*_*S*,_ *ϕ*_*I*_)→(0,0,0) as follows.

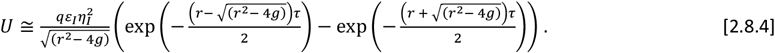

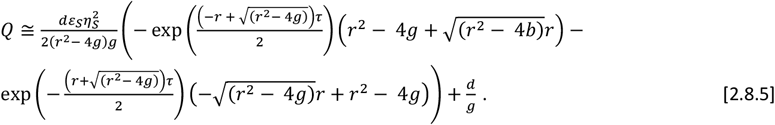

Here the terms *r, g* and *d* are defined as follows.

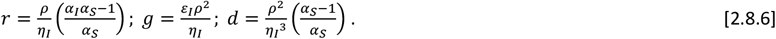

Clearly, one can conclude from **Eqs. 2.8.1** and **2.8.4** that there exist four different timescales viz. two in the pre-steady state regime and two corresponding to the post steady state regimes corresponding to enzyme-substrate and enzyme-inhibitor conversions. We denote them as (τ_*S*1_, τ_*S*2_, τ_*I*1_, τ_*I*2_). From **Eqs. 2.8.1** and **2.8.4** one can define these timescales as follows.

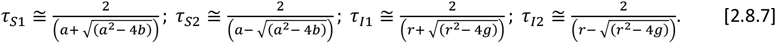

The terms *a, b, c, r, g* and *d* are defined as in **Eqs. 2.8.3** and **2.8.6**. Here τ_*S*1_ and τ_*S*2_ are the pre-steady state and post steady state timescales corresponding to enzyme-substrate dynamics and τ_*I*1_ and τ_*I*2_ are the pre-steady state and post steady state timescales associated with the enzyme-inhibitor dynamics. The errors in various QSSAs will decrease when the timescale separation ratios tend towards zero.

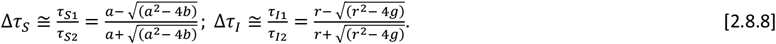

**Eqs. 2.8.1-7** can approximately describe the dynamics of fully competitive enzyme kinetics scheme over the entire (V, U) space in the parametric form when the conditions associated with **Eqs. 2.8.1** and **Eqs. 2.8.4** are true.

### 2.9. Partial competitive inhibition

The differential rate equations corresponding to the Michaelis-Menten type partial competitive inhibition **Scheme B** can be written as follows.

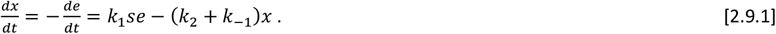

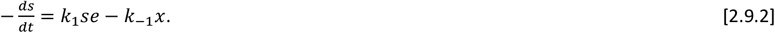

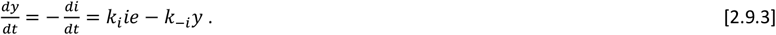

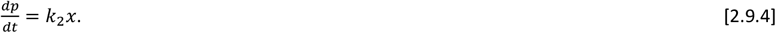

Here the dynamical variables (s, e, x, y, i, p) denote respectively the concentrations (M) of substrate, enzyme, enzyme-substrate complex, enzyme-inhibitor complex and inhibitor. Further k_1_ and k_i_ are the respective bimolecular forward rate constants (1/M/s) and k_-1_ and k_-i_ are the respective reverse rate constants (1/s), k_2_ is the product formation rate constant (1/s). Here the initial conditions are (s, e, x, y, i, p) = (s_0_, e_0_, 0, 0, i_0_, 0) at t = 0. The mass conservation laws are *e* = *e*_0_ − *x* − *y* ; *s* = *s*_0_ − *x* − *p* ; *i* = *i*_0_ − *y*. When *t*→*∞*, then the reaction trajectory ends at (s, e, x, y, i, p) = (0, e_∞_, 0, y_∞_, i_∞_, s_0_) where (e_∞_, y_∞_, i_∞_) are the equilibrium concentrations of free enzyme, enzyme-inhibitor complex and free inhibitor. The steady state of the enzyme-substrate complex occurs at the time point 0 *< t*_*CS*_*< ∞* when 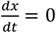 and the steady state of the enzyme-inhibitor complex occurs at the time point 0 *< t*_*Cs*_ *< ∞* when 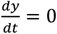 where one also finds from **Eq. 2.9.3** 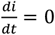. Similar to the scaling transformations in **Eqs. 2.2.7-9, Eqs. 2.9.1-4** can be reduced to the following set of dimensionless equations.

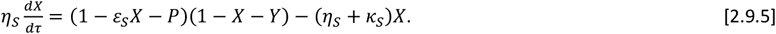

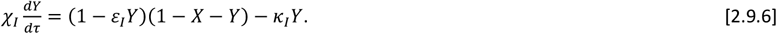

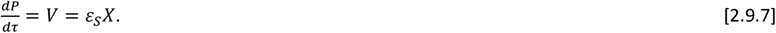

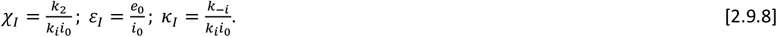

Here (S, E, X, Y, I, P) ∈ [0,1]. The mass conservation relations in the dimensionless form can be written as *I*= (1 − *ε*_*I*_*Y*), E + X + Y = 1 and V + P + S = 1. Other dimensionless parameters are defined similar to the case of fully competitive inhibition scheme as given in **Eqs. 2.2.2-6**. Here (*χ*_*I*,_ *η*_*S*_) are singular perturbation parameters and (*ε*_*I*,_ *ε*_*s*,_ *κ*_*s*,_ *κ*_*I*_) are the ordinary perturbation parameters. The initial conditions in the dimensionless space are (S, E, X, Y, I, P) = (1, 1, 0, 0, 1, 0) at τ = 0. When *τ*→*∞*, then the reaction ends at (S, E, X, Y, I, P) = (0, E_∞_, 0, E_∞_, I_∞_, 1) where the terms (E_∞_, Y_∞_, I_∞_) are the final equilibrium concentrations of the free enzyme, enzyme-inhibitor complex and free inhibitor. When (*χ*_*I*,_ *η*_*S*_)→(0,0), then **Eqs. 2.9.5-6** can be equated to zero and solved for (X, Y). Under these conditions, upon converting X into the velocity using *V* = *ε*_*S*_*X* as given in **Eq. 2.5.7** one obtains the following sQSSA results.

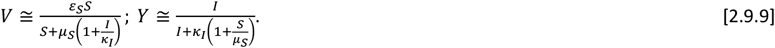

Upon applying the stationary reactants assumptions (*S, I*)→(1,1) under the condition that (*η*_*S*,_ *ε*_*s*,_ *ε*_*I*_, *χ*_*I*_)→(0,0,0,0), one finally obtains the following steady state expressions.

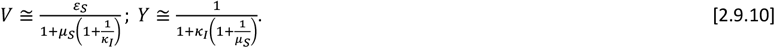

In terms of original variables, the sQSSA reaction velocity V defined in **Eqs. 2.9.10** can be written as follows.

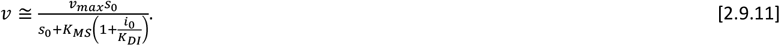

**Eq. 2.9.11** is generally used to estimate the kinetic parameters from the experimental datasets on the partial competitive inhibition scheme using double reciprocal plotting methods where the effective *K*_*MS*_increases linearly with i_0_. Upon substituting the conservation law *I*= (1 − *ε*_*I*_*Y*) in **Eqs. 2.9.9** for Y and subsequently solving the resulting quadratic equation for Y, one obtains the following post-steady state approximations in the (V, S) and (Y, S) spaces under the conditions that (*χ*_*I*_, *η*_*S*_)→(0,0).

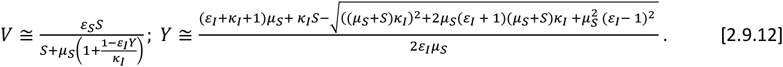

By setting *I*= (1 − *ε*_*I*_*Y*) (where Y is expressed as a function of S as given in **Eq. 2.9.12**) in the expression of V in **Eqs. 2.9.9**, one obtains the post steady state approximation in the (V, S) space. Using the mass conservation laws, one can directly obtain the post-steady state approximation for *P* = 1 − *V* − *S*. Post-steady state approximation in the (I, S) space can be expressed in a parametric form where *S∈* [0,1] acts as the parameter. One can obtain the exact equilibrium values *I*_*∞*_ and *Y*_*∞*_ by setting *S*→0 in **Eqs. 2.9.12** as follows.

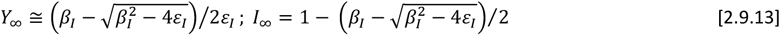

Here we have defined *β*_*I*_= 1 + *ε*_*I*_+ *κ*_*I*_. In **Eq. 2.9.13**, one finds from the mass conservation law that *I*_*∞*_ = 1 − *ε*_*I*_*Y*_*∞*_.

#### 2.9.1. Variable transformations

Using the transformation 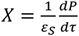 along with the other scaling transformations given in **Eqs. 2.2.7-9, Eqs. 2.9.5-7** can be reduced to the following set of coupled nonlinear second order ODEs in the (P, Y, τ) space.

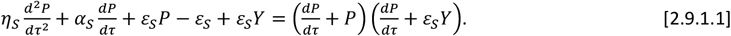

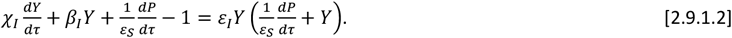

Here the initial conditions are 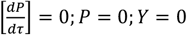 at τ = 0. **Eqs. 2.9.1.1-2** can be transformed into the (V, Y, P) and (V, Y, S) spaces using 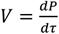 and mass conservation laws as follows.

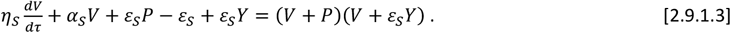

Upon substituting *P* = 1 − *V* − *S*in this equation one finds that,

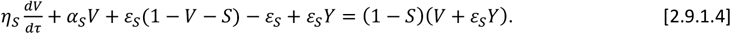

Upon the substitution of 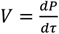 in **Eq. 2.9.1.2** one finds that,

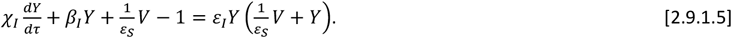

In this equation *β*_*I*_is defined as follows.

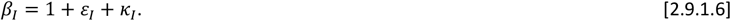

Here the initial conditions are *V* = 0; *P* = 0; *Y* = 0 at τ = 0. **Eqs. 2.9.1.1-6** completely characterize the partial competitive inhibition in the (V, Y, P) and (V, Y, S) spaces from which we will derive the following approximations.

#### 2.9.2. Post-steady state approximations

**Case I**: When (*η*_*S*,_ *χ*_*I*,_ *ε*_*s*,_ *ε*_*I*_)→0, then using X = V/ε_S_, **Eqs. 2.9.1.3-5** can be approximated in the (X, Y, P) space as follows.

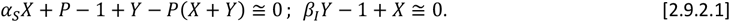

Upon solving **Eqs. 2.9.2.1** for (X, Y) and noting that *S ≅* 1 − *P, I ≅* 1 in such conditions, we obtain the following results similar to **Eqs. 2.4.1** related to the fully competitive inhibition scheme in the (V, S, I) and (Y, S, I) spaces.

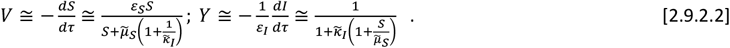

Here we have defined 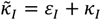. The expression for V in **Eqs. 2.9.2.2** is similar to the one in **Eqs. 2.9.9** where *μ*_*S*_and *κ*_*I*_are replaced with 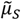 and 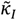 and I = 1 in the definition of Y. The post steady state approximation in the (S, I) space can be expressed in a parametric form using *I*= 1 − *ε*_*I*_*Y*. Here Y is given in terms of S as in **Eq. 2.9.2.2** where *S∈* [0,1] acts as the parameter. Upon applying the transformation 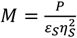 in **Eqs. 2.9.1.1-2**, one finally arrives at the following set of coupled nonlinear second order ODEs in the (M, Y, τ) space.

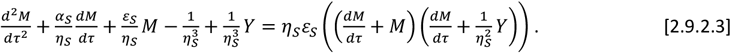

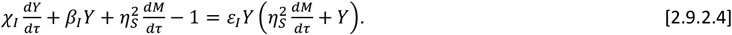

Here the initial conditions are 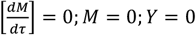 at τ = 0. **Eqs. 2.9.2.3-4** are the central equations corresponding to the Michaelis-Menten type enzyme kinetics with partial competitive inhibition. Using **Eqs. 2.9.2.3-4**, one can derive several approximations under various limiting conditions as follows.

**Case II:** When (*χ, ε, ϕ*_*S*_)→0, then from **Eq. 2.9.2.4** one finds that 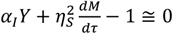 which results in the approximation 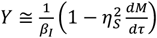. Upon substituting this approximation for Y in **Eqs. 2.9.2.3**, and setting ϕ_*S*_= *η*_*S*_*ε*_*S*_→0, one arrives at the following second order linear ODE corresponding to the (M, *τ*) space.

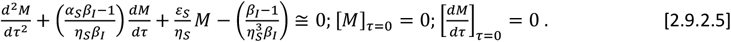

Upon solving this second order ODE for M with respect to the given initial conditions and then reverting back to the velocity V using the transformation 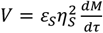, one can obtain the following expression in the (V, *τ*) space under the conditions that (*ε*_*I*_, *χ*_*I*_, ϕ_*S*_)→(0,0,0).

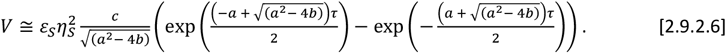

The terms *a, b* and *c* in **Eq. 2.9.2.6** are defined as follows.

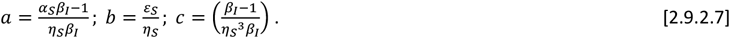

Upon solving *dV/d*τ = 0 for τ in **Eq. 2.9.2.6**, one can obtain the following expression for the steady state timescale associated with the enzyme-substrate complex.

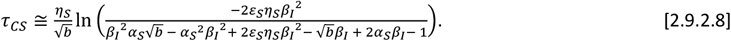

In this equation, *b* is defined as follows.

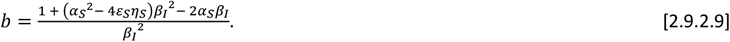

#### 2.9.3. Pre-steady state approximations

**Case III:** When *η*_*S*_→0, then one obtains the uncoupled equation in the (Y, τ) space from **Eq. 2.9.2.4** as 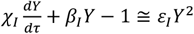. The integral solution of this first order nonlinear ODE with the initial condition [*Y*] _*τ* =0_ = 0 can be written as follows.

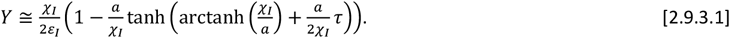

In this equation 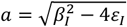. **Eq. 2.9.3.1** suggests the following the timescale associated with the enzyme-inhibitor complex to attain the steady state under the conditions that *η*_*S*_→0.

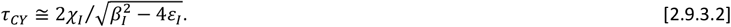

Clearly, **Eqs. 2.9.1.3-5** can have common steady states only when τ_*Cs*_ = τ_*CS*_. We will show in the later sections that when τ_*Cs*_ *>* τ_*CS*_, then the substrate depletion with respect to time will show a typical non-monotonic trend since the inhibitor reverses the enzyme-substrate complex which is formed before time τ_*Cs*_ in the pre-steady state regime with respect to the enzyme-substrate complex. This phenomenon introduces significant amount of error in the QSSAs. Using the transformation 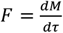, **Eqs. 2.9.2.3-4** can be rewritten in the (F, Y, M, τ) space with the initial conditions F = 0 at M = 0 and Y = 0 at τ = 0 as follows.

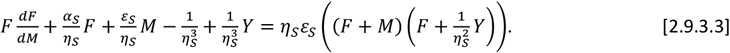

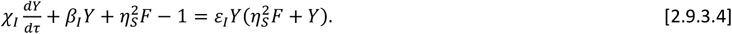

**Case IV:** When the conditions (*ε*_*S*,_ *χ*_*I*,_ *ε*_*I*_)→0 are true, then one can derive the approximate differential rate equations governing the pre-steady state dynamics in the (F, M) and (F, Y) spaces from **Eqs. 2.9.3.3-4** as follows.

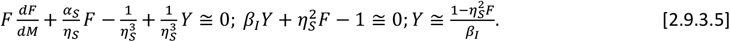

The initial condition corresponding to **Eqs. 2.9.3.5** is F = 0 at M = 0. Upon substituting the expression of Y into the differential equation corresponding to the (F, Y, M) space given in **Eqs. 2.9.3.5**, one finally obtains the following approximation.

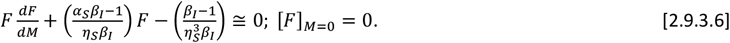

Upon solving **Eq. 2.9.3.6** with the given initial conditions, and then reverting back to (V, P) space using the transformations 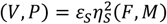, one finally obtains the following expressions for the pre-steady state regime in the (V, P) space under the conditions that (*ε*_*s*,_ *χ*_*I*_, *ε*_*I*_)→0.

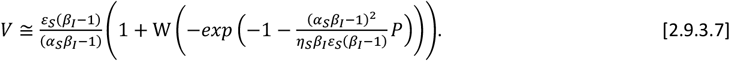

To obtain the expression for the (V, S) space one needs to first solve **Eqs. 2.9.3.6** implicitly in the (V, P) space. Then upon inverting the solution for V in terms of Lambert W function after the substitution of the mass conservation law P = 1 – V – S as demonstrated in **Appendix B**, one finally arrives at the following expression.

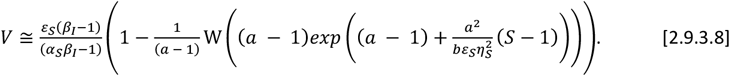

In this equation, the terms *a* and *b* are defined as follows.

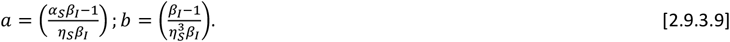

By expanding the right-hand side of **Eq. 2.9.3.8** in a Taylor series around S = 1, one finds that *V ≅* 1 − *S*+ *O*((*S*− 1)^2^) which means that *P ≅* 0 in the pre-steady state regime. One can directly translate the (V, S) space approximation given in **Eq. 2.9.3.8** into the pre-steady state of (Y, S) space under the conditions that (*ε, χ, ε*)→0 using **Eqs. 2.9.3.5** that results in 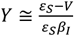. But there is a mismatch in the required initial condition for Y in this expression i.e., Y = 0 at S = 1. Particularly, **Eqs. 2.9.3.5** sets the initial condition for Y as 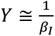 at S = 1 (at which V = 0 and therefore F = 0) that is inconsistent since *Y ≠* 0 at τ = 0 or S = 1. Detailed numerical analysis of the (Y, S) space trajectories suggests an approximate expression as 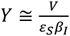 for the pre-steady state regime under the conditions that (*ε*_*s*,_ *χ*_*I*,_ *ε*_*I*_)→(0,0,0). Explicitly, one can write this approximation derived from numerical analysis as follows.

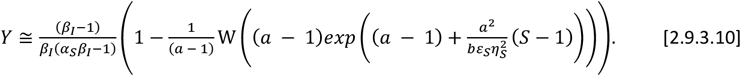

Here *a* and *b* are defined as in **Eqs. 2.9.3.9**. By expanding **Eq. 2.9.3.10** in a Taylor series around S = 1, one finds that 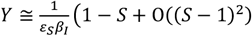. Using **Eq. 2.9.3.10** and the conservation law *I*= (1 − *ε*_*I*_*Y*), one can express I as a function of S in the pre-steady state regime of (I, S) space under the condition that (*ε*_*s*,_ *χ*_*I*,_ *ε*_*I*_)→0. Similarly, in (F, τ) space the differential equation **Eq. 2.9.3.6** can be written as follows.

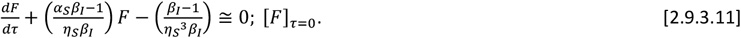

Upon solving **Eq. 2.9.3.11** with the given initial condition and then reverting back to (V, τ) space using the transformation 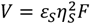, one obtains the following integral solution corresponding to the reaction velocity in the pre-steady state regime.

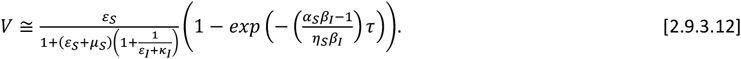

This equation at τ→*∞* along with **Eq. 2.9.3.2** suggest the following expressions for the steady state timescale associated with the enzyme-substrate and enzyme inhibitor complexes under the conditions that (*ε*_*I*_, *χ*_*I*,_ *ε*_*S*_)→(0,0,0).

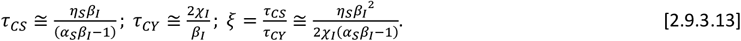

Clearly, common steady states between enzyme-substrate and enzyme-inhibitor complexes can occur only when the ratio 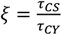 in **Eqs. 2.9.3.13** becomes as *ξ ≅* 1 at which the error associated with various QSSAs will be at minimum. Upon solving the minimum error condition *ξ ≅* 1 for *β*_*I*_, one obtains the following two possible roots.

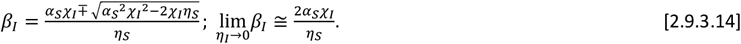

When *η*_*I*_→0, then from **Eqs. 2.9.3.13-14** one finds the following expression for *ξ*.

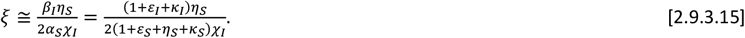

When *η*_*S*_*≅* 2*χ*_*I*_, then **Eq. 2.9.3.15** suggests that the condition *ξ ≅* 1 can be achieved by simultaneously settings sufficiently large values for any one of the parameters in the numerator (*ε*_*I*_, *κ*_*I*_) and any one of the parameters in the denominator (*ε*_*s*,_ *χ*_*I*_, *κ*_*S*_). When P, S and τ become sufficiently large, then **Eqs. 2.9.3.7, 2.9.3.12** asymptotically converge to the following limiting value that is close to the steady state reaction velocity V associated with the partial competitive enzyme kinetics described in **Scheme B** in the limit (*ε*_*s*,_ *χ*_*I*_, *ε*_*I*_)→(0,0,0). We denote this approximation as V_3_. We will show in the later sections that this approximation works very well in predicting the steady state reaction velocity of the partial competitive inhibition **Scheme B** over wide ranges of parameters.

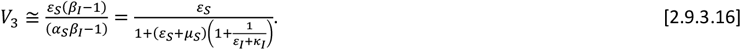

In terms of original variables, **Eq. 2.9.3.16** can be written as follows.

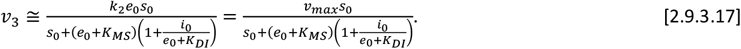

#### 2.9.4. Error in the standard QSSA of the partial competitive inhibition scheme

Similar to **Eqs. 2.7.1-2**, the overall error in the refined form of sQSSA with stationary reactant assumption as given in **Eqs. 2.9.2.2** with *S≅* 1 corresponding to the partial competitive inhibition can be computed using the pre-steady state approximation given in **Eqs. 2.9.3.12** as follows [37].

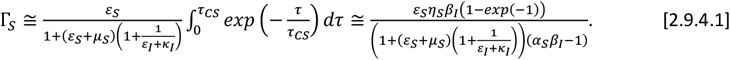

Here *τ*_*CS*_ is defined as in **Eqs. 2.9.3.13**. Upon solving 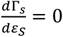 for *ε*_*S*_, one obtains the following expression for *ε*_*S,max*_at which the error in the standard QSSA attains maximum. This means that the sufficient condition to minimize such error will be given as follows.

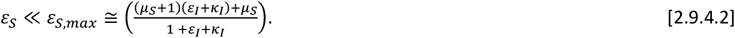

Inequality in **Eq. 2.9.4.2** (we denote this by E_3_) can be explicitly written in the following form.

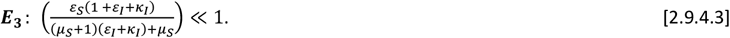

**Case IV**: When (*η*_*S*,_ *χ*_*I*_)→0, then **Eqs. 2.9.1.3-4** reduce to the following form in the (V, Y, P) and (V, Y, S) spaces.

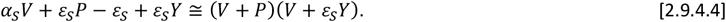

Upon using the conservation law *P* = 1 − *V* − *S*in **Eq. 2.9.4.4**, one obtains the following.

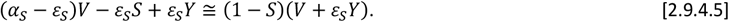

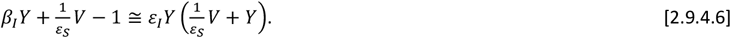

Upon solving **Eqs. 2.9.4.4-6** for (V, Y), one obtains the following expressions for V, P, Y and I in terms of S under the conditions that (*η*_*S*,_ *χ*_*I*_)→(0,0).

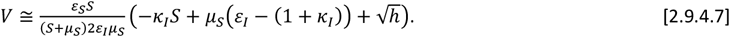

In this equation 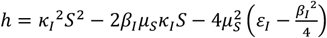. We will show later that **Eq.2.9.4.7** can predict the post-steady state reaction velocity much better than **Eqs. 2.9.2.2** in the (V, S) space. Noting that V + P + S = 1, one can derive the approximate expression for P in terms of S under the conditions that (*η*_*S*,_ *χ*_*I*_)→(0,0) as follows.

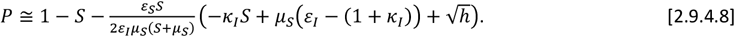

**Eqs. 2.9.4.7-8** can be used to generate trajectories in the post-steady state of (V, P), (V, P, S) and (P, S) spaces in the parametric form where *S∈* [0,1] acts as the parameter. Upon solving **Eqs. 2.9.4.4-6** for Y, the post-steady state approximation in the (Y, S) space can be written as follows.

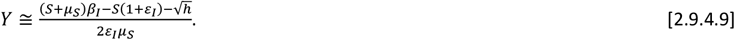

This equation is more refined one than **Eqs. 2.9.12**. Noting that *I*= (1 − *ε*_*I*_*Y*), one can obtain the following approximate expression for the inhibitor concentration in terms of S corresponding to the post-steady state of (I, S) space.

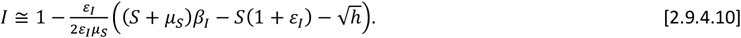

In **Eqs. 2.9.4.8-10**, *h* is defined as in **Eq. 2.9.4.7**. Solutions obtained under the conditions that (*η*_*S*,_ *χ*_*I*_)→(0,0) can approximate the original trajectory in the (V, S) space very well only in the post-steady state regime. By setting *S*→0 in **Eqs. 2.9.4.9-10**, one obtains the exact equilibrium values of (Y, I) similar to **Eqs. 2.9.12** as follows.

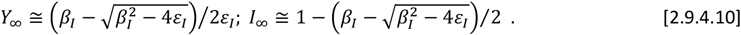

In terms of original variables, the steady state velocity approximation corresponding to the stationary reactant assumption *S*→1 can be written from **Eqs. 2.9.4.7** as follows.

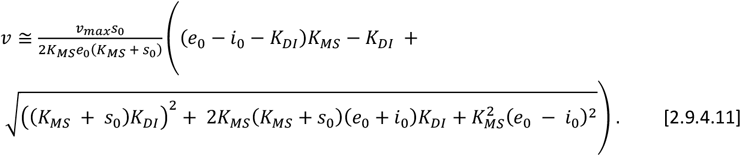

Upon inserting the experimental values of e_0_ and *i*_0_ in to this equation one can directly extract the values of *K*_*Ms*,_ *K*_*DI*_and *υ*_*max*_from the data on velocity versus total substrate concentrations using non-linear least square fitting methods.

#### 2.9.5. Steady state substrate and inhibitor levels

Similar to earlier studies [37], one can approximate the steady state substrate concentration by finding the intersection point between the pre- and post-steady state approximations in the (V, S) and (U, I) spaces. In case of partial competitive inhibition under the conditions that (*η*_*S*,_ *χ*_*I*,_ *ε*_*s*,_ *ε*_*I*_)→0, the steady state substrate level S_C_ can be obtained by finding the intersection point between the pre-steady state approximation given by **Eqs. 2.9.3.8** and the post steady state approximation given by **Eqs. 2.9.2.2** in the (V, S) space as follows.

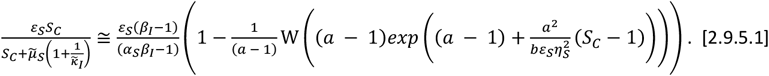

In this equation, left hand side is the post-steady state approximation and the right-hand side is the pre-steady state approximation, *S*_*C*_ is the intersection point that approximates the steady state substrate concentration and, a and b are defined as in **Eqs. 2.9.3.9**. Upon expanding the right-hand side of **Eq. 2.9.5.1** in a Taylor series around S_C_ = 1 and ignoring second and higher order terms one finds the following equation for the intersection between pre- and post-steady state solution in the (V, S) space.

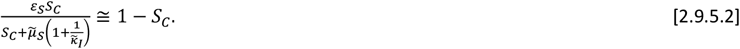

Upon solving this quadratic equation for S_C_ one obtains the following approximation for the steady state substrate level.

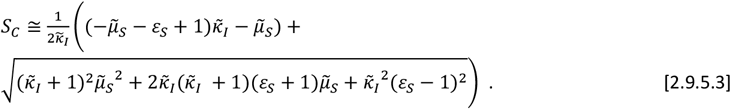

By substituting this expression of S_C_ into the post-steady state approximation given in **Eq. 2.9.2.2**, one can obtain the following expression for the steady state velocity V_C_.

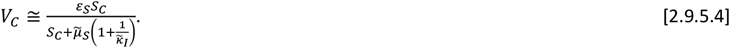

**Eqs. 2.9.5.1-4** are valid only under the conditions that (*η*_*S*,_ *χ*_*I*,_ *ε*_*s*,_ *ε*_*I*_)→0. One can also substitute the approximate value of S_C_ obtained from **Eq. 2.9.5.3**, into **Eq. 2.9.4.7** to obtain a refined steady state velocity as follows.

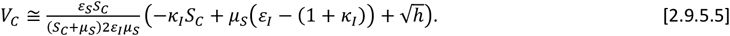

In this equation 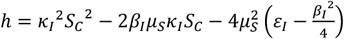 where S_C_ is defined as in **Eq. 2.9.5.3**. More accurate value of S_C_ can be obtained from the intersection point of the post-steady state velocity given by **Eqs. 2.9.4.7** and the pre-steady state velocity given by **Eq. 2.9.3.8**. Particularly, the intersection point is the real root of the following cubic equation that is valid under the conditions that (*η*_*S*,_ *χ*_*I*_)→(0,0).

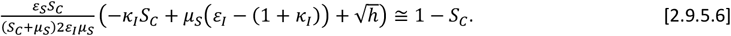

In this equation **Eq. 2.9.3.8** is approximated as *V ≅* 1 − *S*. In case of fully competitive inhibition scheme, when *δ ≠* 1 then the second prolonged steady state levels of (S, I) are given by **Eqs. 2.4.2.1-2**. The approximate first transient steady state values can be obtained from the intersections of the pre- and post-steady state solutions given by **Eqs. 2.4.23** and **Eqs. 2.6.10** and **2.6.23**. Upon expanding the pre-steady state solutions given in **Eqs. 2.6.10** for (V, S) and Eqs. **2.6.23** for (U, I) space in a Taylor series around (S, I) = (1,1) and ignoring the second order terms, one finds the following equations for the intersection points (S_C_, I_C_) of the pre- and post-steady state approximations in the (V, S) and (U, I) spaces.

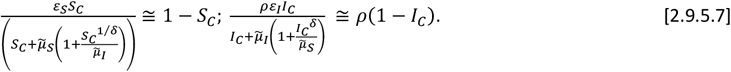

These equations are valid under the conditions that (*η*_*S*,_ *η*_*I*,_ *ε*_*s*,_ *ε*_*I*_)→0 irrespective of the values of *δ*. When *δ <* 1, then the first steady state value S_C_ in the (V, S) space can be obtained by solving the following equation for Z.

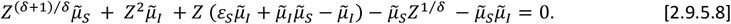

When *δ >* 1, then the primary steady state value I_C_ in the (U, I) space can be obtained by solving the following equation for R.

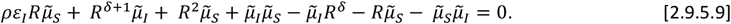

When δ = 1, then explicit expressions for the steady state substrate level can derived by solving the quadratic equation arising from **Eqs. 2.9.5.8** and choosing the appropriate root as follows.

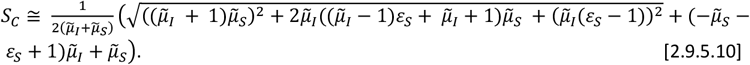

When δ = 1, then explicit expressions for the steady state inhibitor level can derived by solving the quadratic equation arising from **Eqs. 2.9.5.9** and choosing the appropriate root as follows.

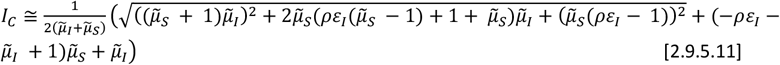

Upon substituting these S_C_ and I_C_ obtained from **Eqs. 2.9.5.8-9** in **Eqs. 2.4.23** for S and I, one can obtain accurate values of the steady state reaction velocities (V_C_, U_C_) as follows.

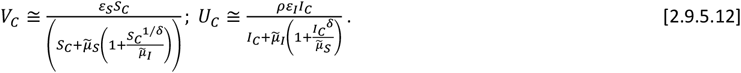

Similar to **Eq. 2.9.5.6**, accurate values of S_C_ and I_C_ can be obtained by finding the intersection points between the post-steady state expressions given in **Eq. 2.4.12** and **2.4.16** and the pre-steady state approximations *V ≅* 1 − *S*and *U ≅ ρ*(1 − *I*). When (*η*_*S*,_ *η*_*I*,_ *Q*)→(0,0,0), then using **Eq. 2.4.12**, the steady state substrate concentration can be given by the real solution of following equation.

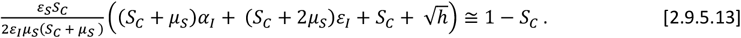

Here 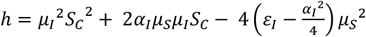. When (*η*_*S*_, *η*_*I*,_ *P*)→(0,0,0), then using **Eq. 2.4.16** the steady state inhibitor concentration *I*_*C*_ can be the given by the real solution of the following equation.

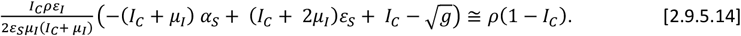

Here 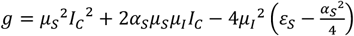. Although **Eqs. 2.9.5.13-14** can accurately predict the steady state levels of substrate S_C_ and inhibitor I_C_, one needs to perform more computations than **Eqs. 2.9.5.10-12**.

### 2.10. Total QSSA of fully competitive inhibition scheme

When the conditions (*η*_*S*,_ *η*_*I*,_ *P, Q*)→(0,0,0,0) are true so that *S≅* (1 − *ε*_*S*_*X*), *I≅* (1 − *ε*_*I*_*Y*) in the pre-steady state regime of the fully competitive inhibition scheme described by **Eqs. 2.2.7-9**, then one can arrive at the total QSSA (tQSSA) [43, 44]. In this situation, **Eqs. 2.2.7-9** can be approximated as follows.

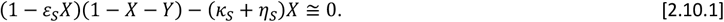

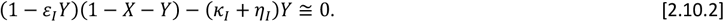

Upon solving **Eqs. 2.10.1-2** one obtains the following tQSSA expressions for V and U.

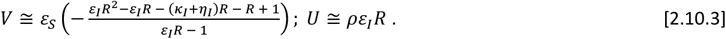

Here R is the appropriate real solution of the following cubic equation.

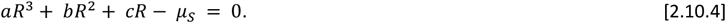

In this cubic equation, the coefficients *a, b* and *c* are defined as follows.

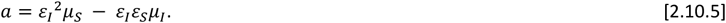

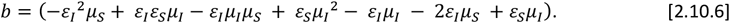

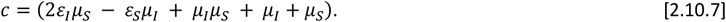

Here *μ*_*S*_= *κ*_*S*_+ *η*_*S*_ and *μ*_*I*_= *κ*_*I*_+ *η*_*I*_. The real positive root of the cubic equation **Eq. 2.10.4** can be written as follows.

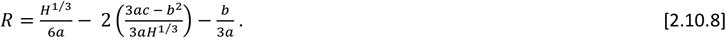

The term H in **Eq. 2.10.8** is defined as follows.

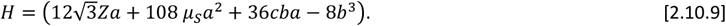

Here Z is defined as follows.

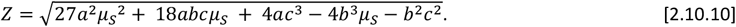

In **Eqs. 2.10.8-10**, *a, b* and *c* are defined as in **Eqs. 2.10.5-7**. In case of partial competitive inhibition, **Eq. 2.10.2** becomes in the limit (*η*_*S*,_ *η*_*I*,_ *P*)→0 as (1 − *ε*_*I*_*Y*)(1 − *X* − *Y*) − *κ*_*I*_*Y ≅* 0 and therefore the solution set given in **Eqs. 2.10.3-4** needs to be modified accordingly.

## 3. Simulation Methods

We use the following Euler iterative scheme to numerically integrate the set of nonlinear rate equations given in **Eqs. 2.2.7-9** corresponding to the fully competitive inhibition **Scheme A**.

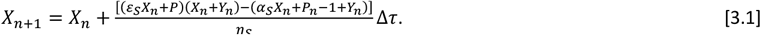

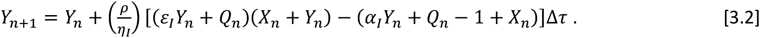

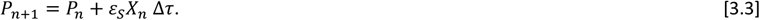

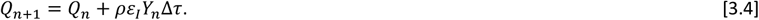

Here the initial conditions are set as (X_0_, Y_0_, P_0_, Q_0_) = (0, 0, 0, 0) at τ = 0. The trajectories of (S, I, E) can be computed from the trajectories of (X, Y, P, Q) using the mass conservation equations E = 1 – X – Y, S = 1 – *ε*_*S*_X – P and I = 1 – *ε*_*I*_Y – Q. We use the following Euler iterative scheme to numerically integrate the nonlinear rate equations given in **Eqs. 2.9.5-7** corresponding to the partial competitive inhibition **Scheme B**.

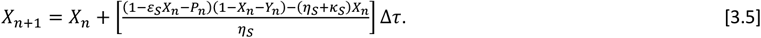

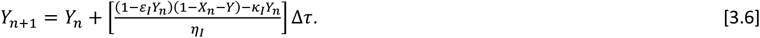

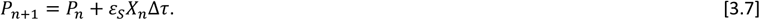

Here the initial conditions are set as (X_0_, Y_0_, P_0_) = (0, 0, 0) at τ = 0. From the trajectories of (X, Y, P) the trajectories of (S, I, E) can be computed using the mass conservation equations E = 1 – X – Y, S = 1 – *ε*_*S*_X – P and I = 1 – *ε*_*I*_Y. We further set *Δ*τ *<* 10^−5^ so that the dynamics with respect to the shortest timescale can be captured. We use the following scheme to numerically integrate the coupled nonlinear ODEs given in **Eqs. 2.5.1-2** under various parameter settings.

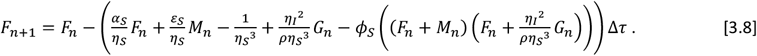

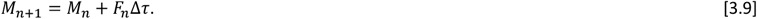

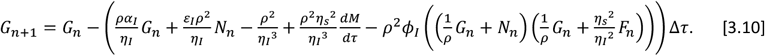

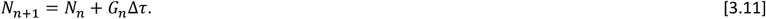

Here the initial conditions are set as (F_0_, M_0_, G_0_, N_0_) = (0, 0, 0, 0) at τ = 0. Using the transformations (*P, V*) = *ε* _*S*_*η* _*S*_^2^(*M, F*) and (*U, Q*) = *ε*_*I*_*η*_*I*_^2^(*G, N*), one can revert back to the original dynamical variables V, U, P and Q. Data in (S, I) space can be obtained using the mass conservation laws S = 1 – V – P and I = 1 – U/ρ – Q. The time at which the steady state occurs was numerically computed from the integral trajectories by looking at the time point at which the first derivative of (X, Y) with respect to τ changes the sign. The trajectories corresponding to the *ϕ-*approximations can also be computed by setting (ϕ_*s*,_ ϕ_*I*_)→(0,0) in **Eqs. 3.8-11** apart from using the integral solutions given in **Appendix A**.

## 4. Results and Discussion

Competitive inhibition of the Michaelis-Menten enzymes plays critical roles in designing drug molecules against the nodal enzymes of various pathogenic organisms. The relative efficiency of an inhibitor type drug depends on the parameters *K*_*MS*_, *v*_*max*_, *u*_*max*_ and *K*_*MI*_. Estimation of these parameters of a given enzyme-substrate-inhibitor system from the experimental data is critical for the screening of various inhibitor type drug molecules against a given enzyme of pathogen both under *in vitro* as well as *in vivo* conditions. Almost all the current experimental methods use the expression derived from the sQSSA with stationary reactant assumption i.e., approximation under the conditions that (*η*_*S*_,*η*_*I*_,*ε*_*S*_,*ε*_*I*_) → (0,0,0,0) (we denote these conditions as **C**_**1**_). Under these conditions along with the stationary reactants assumption that (*S,I*) ≅ (1,1) (condition **C**_**2**_), one can approximate the steady state reaction velocities for the fully competitive inhibition scheme as 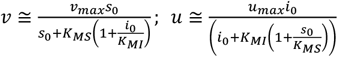as given in **Eqs. 2.4.3** and 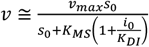 for the partial competitive inhibition scheme as given in **Eqs. 2.9.11**. Reciprocal plots of the dataset on the total substrate s_0_ and inhibitor i_0_ concentrations versus the steady state reaction velocities (*𝒱,u*) combined with linear least square fitting procedures will eventually reveal the required kinetic parameter values. Clearly, the overall error associated with this procedure can be minimized only when the inequality conditions E_1_ and E_2_ given by **Eqs. 2.7.5-6**, C_1_ and C_2_ are true. These mean that **Eqs. 2.4.3** will be valid only when conditions C_1_, C_2_, E_1_, and E_2_ are all true. In the same way, the inequality condition E_3_ given by **Eqs. 2.9.4.3** should be true to minimize the error in the sQSSA based estimation of kinetic parameters of the partial competitive inhibition scheme. In this context, we have obtained here several approximate solutions to the fully as well as partial competitive inhibition schemes over both pre- and post-steady state regimes. These phase-space approximations are summarized in **Table 2** and **Table 3** along with the respective conditions of validity.

**TABLE 2.**
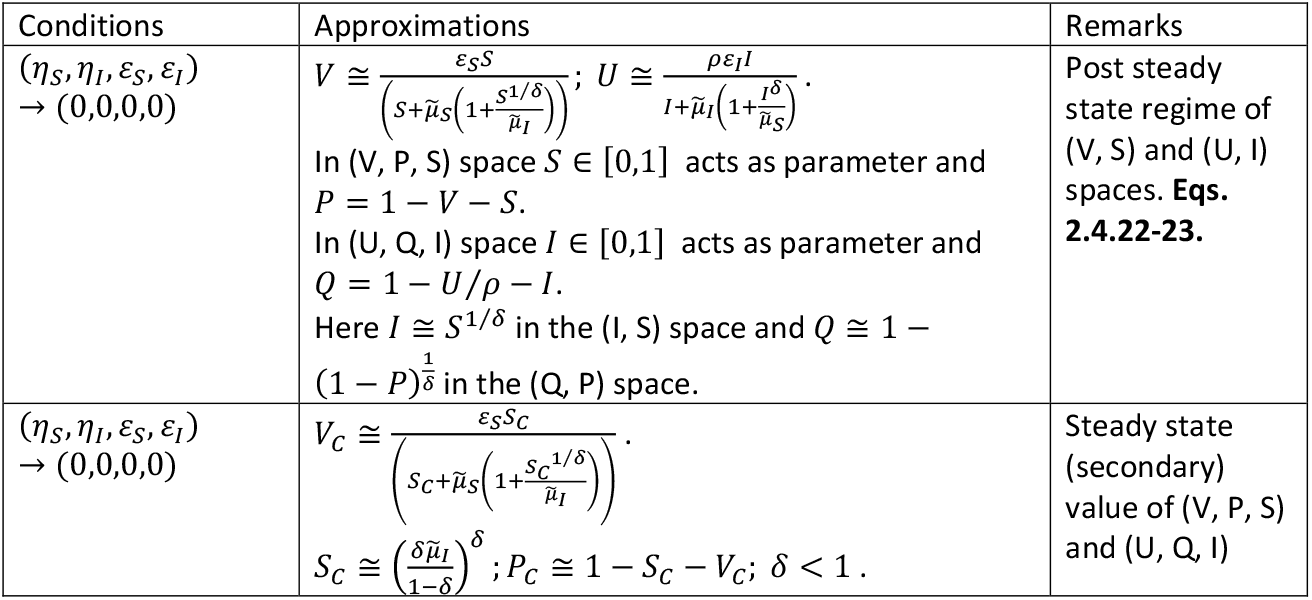
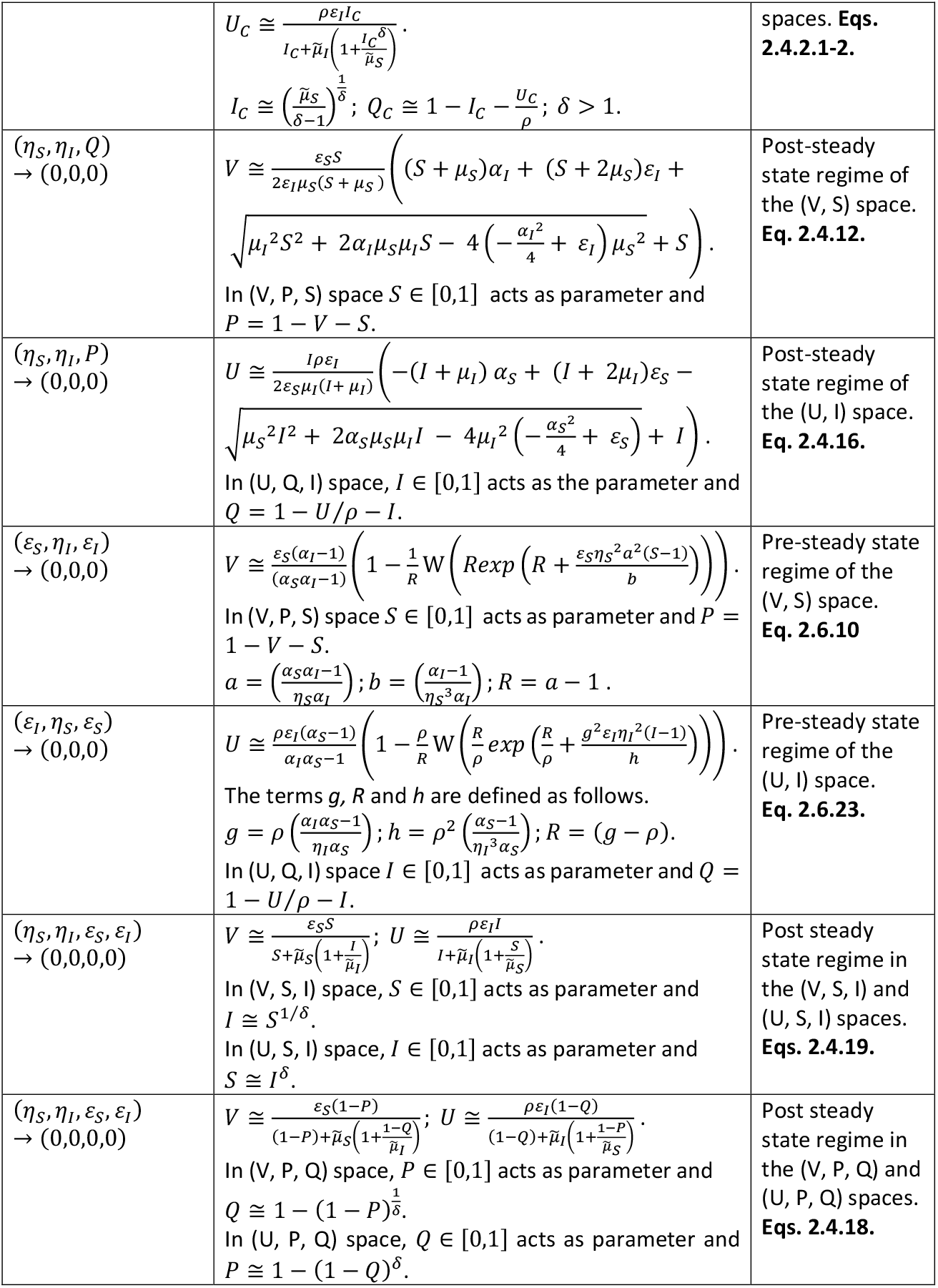
Phase-space approximations of fully competitive inhibition scheme.

**TABLE 3.**
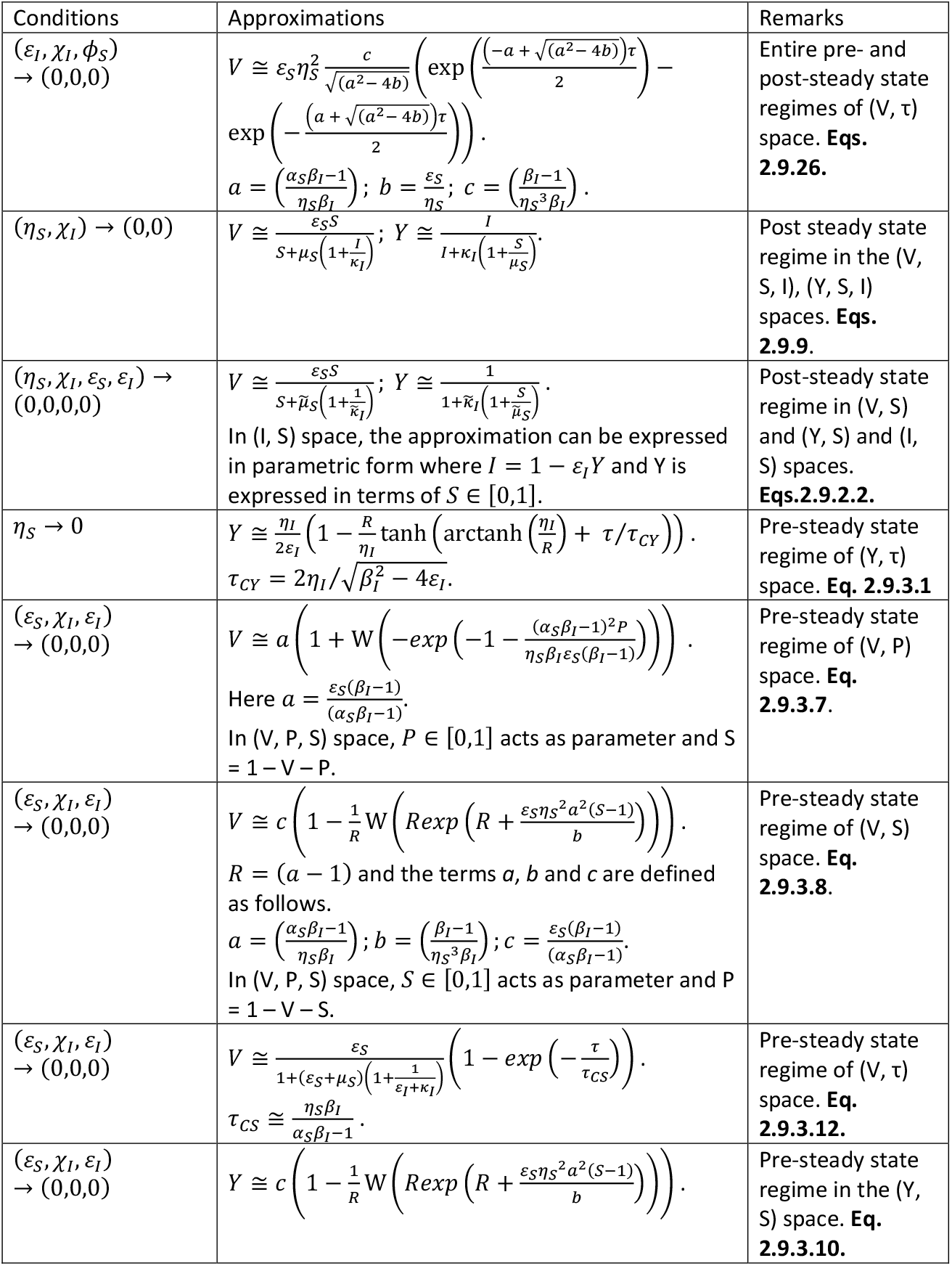
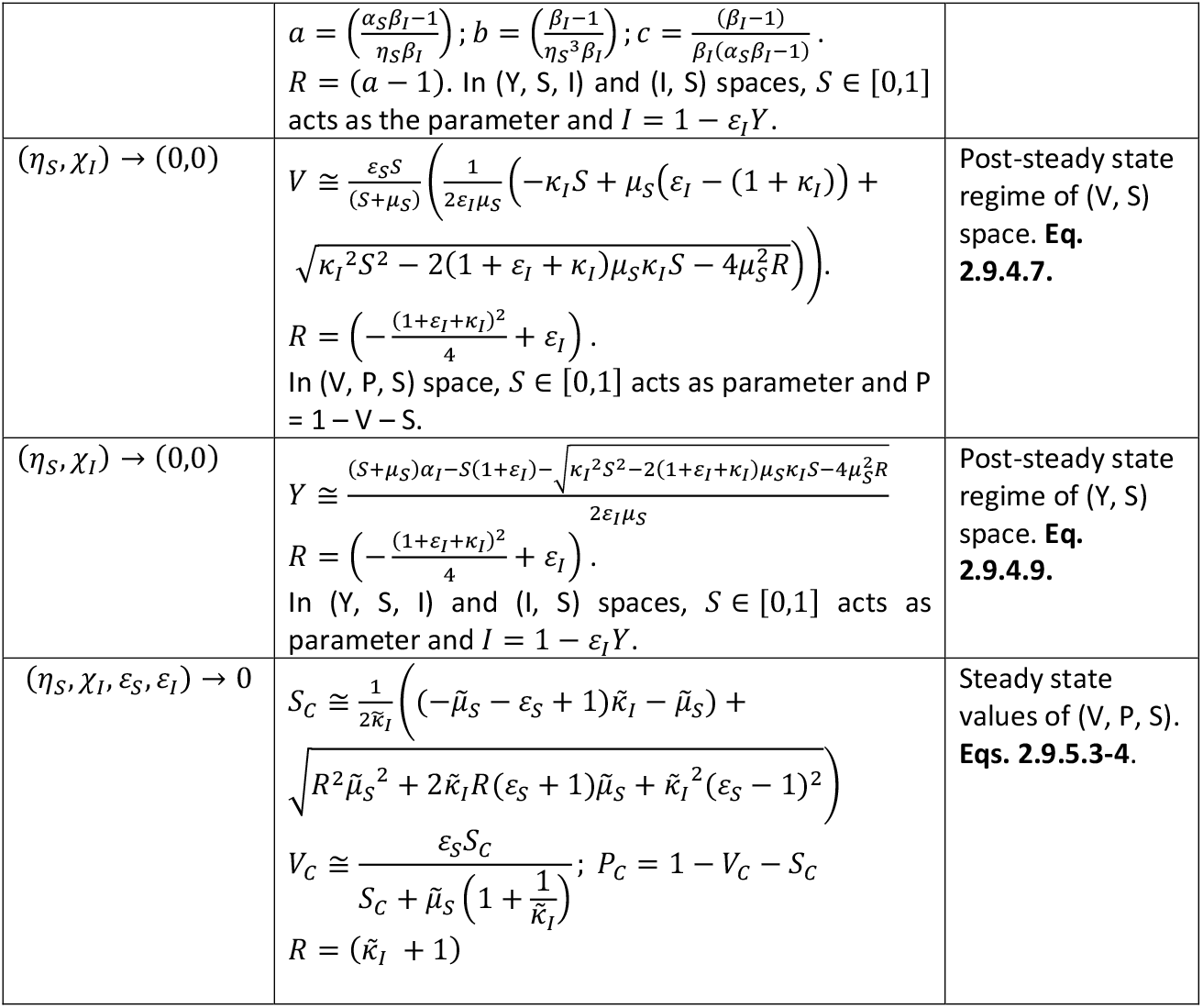
Phase-space approximations of the partial competitive inhibition scheme.

### 4.1. Refined expressions for the sQSSAs

Upon observing the asymptotic behavior of the pre-steady state regime approximations in the (V, S) and (V, P) spaces, we obtained refined expressions for the standard QSSA with stationary reactant assumption i.e., 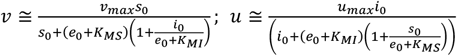 as given in **Eqs. 2.6.13** and **2.6.26** for the fully competitive inhibition scheme and 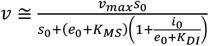 as given in **Eq. 2.9.4.17** for the partial competitive inhibition scheme. These approximations seem to be applicable even when the condition C1 is violated. Particularly, these refined expressions can approximate the steady state reaction velocities even at high values of (*ε*_*S*_,*ε*_*I*_). Upon using of these refined expressions, one can significantly reduce the error in the estimation of various enzyme kinetic parameters from the experimental datasets.

### 4.2. Error in the sQSSA of fully competitive inhibition scheme

The conditions given in C_1_ can be set to a required value under *in vitro* scenarios by manipulating the relative concentrations of the enzyme, substrate and inhibitor. Remarkably, **Eq. 2.4.2.1-2** which deals with the critical parameter *δ* clearly reveals the validity of the stationary reactant assumptions associated with the condition C_2_. From **Eq. 2.4.2.1** one can show that the prolonged secondary steady state substrate level becomes as *S*_*C*_ ≅ 1 when *δ* < 1 and *δ* → 0. This means that the enzyme-substrate complex will exhibit multiple steady states when *δ* < 1 and *δ* → 0. The deviation from the sQSSA of enzyme-substrate complex that represents the primary steady state increases with respect to decrease in *δ* and subsequently the error in the substrate conversion velocity will be negatively correlated with *δ*. Whereas, sQSSA works very well for the enzyme-inhibitor complex with single steady state when *δ* < 1 and *δ* → 0 (**Figs. 4A** and **4B**). Particularly, the approximation given in **Eq. 2.4.16** under the conditions (*η*_*S*_,*η*_*I*_,*P*) → (0,0,0) works very well in the (U, I) space as shown in **Fig. 4B**. On the other hand, from **Eqs. 2.4.1.2** one can show that the prolonged secondary steady state inhibitor level becomes as *I*_*C*_ ≅ 1 when *δ* > 1 and *δ* → ∞. This means that the enzyme-inhibitor complex will exhibit multiple steady states when *δ* > 1 and *δ* → ∞. The deviation from the sQSSA of enzyme-inhibitor complex that represents the primary steady state increases with respect to increase in *δ* and subsequently the error in the substrate conversion velocity will be positively correlated with *δ*. Whereas, sQSSA works very well for enzyme-substrate complex with single steady state when *δ* > 1 and *δ* → ∞ (**Figs. 3C**). Particularly, the approximation given in **Eq. 2.4.12** under the conditions (*η*_*S*_,*η*_*I*_,*Q*) → (0,0,0) works very well in the (V, S) space as shown in **Fig. 4C**.

**FIGURE 4.**
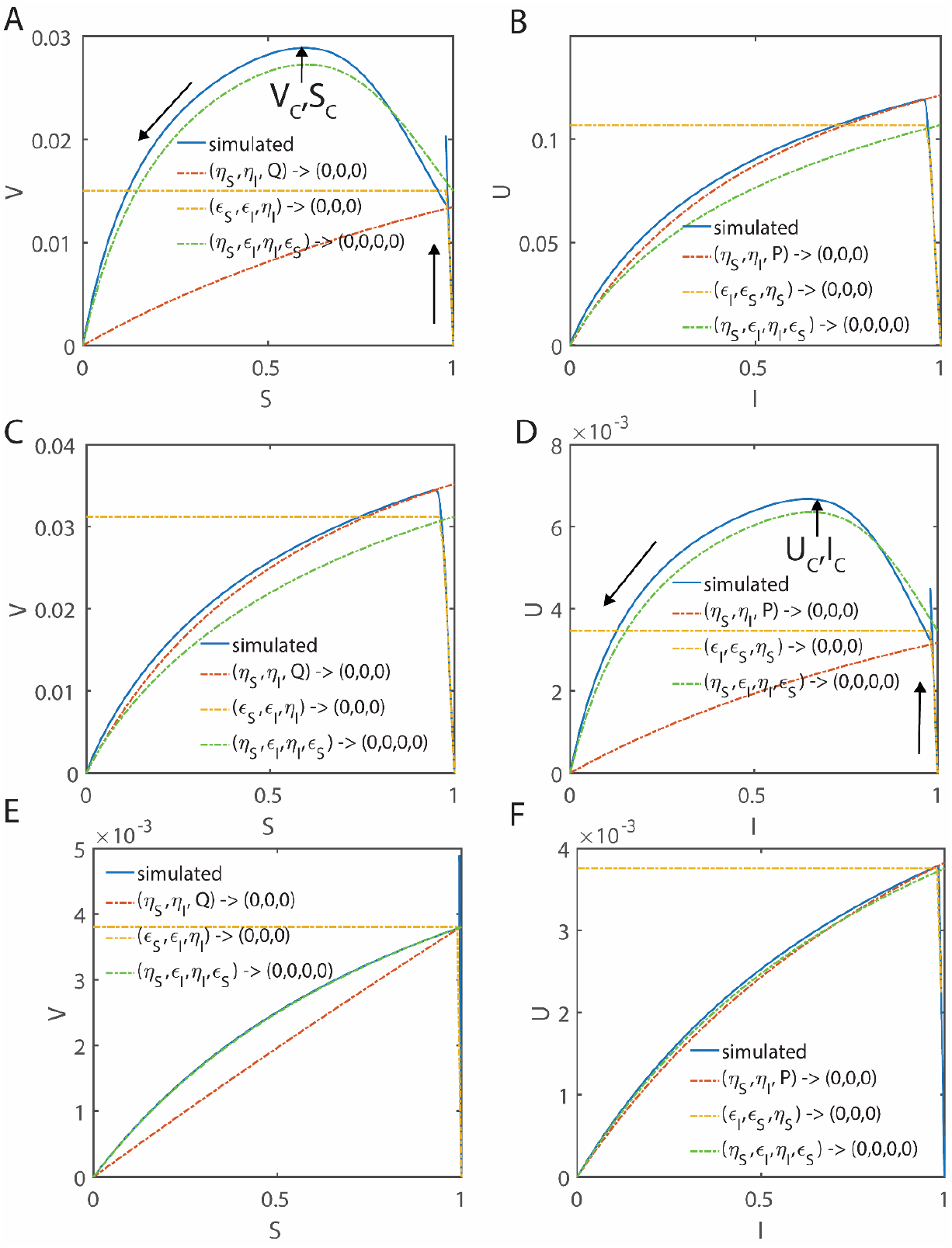
Pre-steady state and post-steady state approximations of the enzyme kinetics with fully competitive inhibition in the velocity-substrate (V, S), velocity-inhibitor spaces (U, I) at different values of δ. The phase-space trajectories start with V = 0 at S = 1 and U = 0 at I = 1, and end at V = 0 at S = 0 and U = 0 at I = 0 with maxima at the steady state. We considered the approximations of (*V, U*) under the conditions (*η*_*S*_, *η*_*I*_, *ε*_*S*_, *ε*_*I*_) → (0,0,0,0) which is the sQSSA in both (V, S) and (U, I) spaces (**Eq. 2.4.23**, refined forms of sQSSA), (*η*_*S*_, *η*_*I*_, *Q*) → (0,0,0) (**Eq. 2.4.12**) and (*η*_*I*_, *ε*_*S*_, *ε*_*I*_) → (0,0,0) (**Eq. 2.6.10**) corresponding to the post and pre-steady state regimes in the (V, S) space, (*η*_*S*_, *ε*_*S*_, *ε*_*I*_) → (0,0,0) (**Eq. 2. 6. 23**) and (*η*_*S*_, *η*_*I*_, *P*) → (0,0,0) (**Eq. 2.4.16**) corresponding to the pre and post steady state regimes of the (U, I) space. Initial conditions for the numerical simulation of **Eqs. 2.2.7-9** are set as (*S, I, E, X, Y, P, Q*) = (1,1,1,0,0,0,0) at τ = 0. **A-B**. The simulation settings are *η*_*S*_ = 0.002, *ε*_*S*_ = 0.04, *κ*_*S*_ = 0.2, *η*_*I*_ = 0.01, *ε*_*I*_ = 0.06, *κ*_*I*_ = 0.1 and ρ = 3.333, σ = 1, δ = 0.1405, ϒ = 3. **C-D**. The simulation settings are *η*_*S*_ = 0.02, *ε*_*S*_ = 0.06, *κ*_*S*_ = 0.1, *η*_*I*_ = 0.003, *ε*_*I*_ = 0.04, *κ*_*I*_ = 0.2 and ρ = 0.225, σ = 1, δ = 9, ϒ = 0.33. **E-F**. Here the settings are *η*_*S*_ = 0.02, *ε*_*S*_ = 0.06, *κ*_*S*_ = 8.1, *η*_*I*_ = 0.003, *ε*_*I*_ = 0.04, *κ*_*I*_ = 1.2 and ρ = 0.225, σ = 1, δ = 1.013, ϒ = 4.5.

When *σ* ≅ 1 and *δ* < 0 or *δ>* 0, then the steady states of enzyme-substrate-inhibitor system show a complex behavior as demonstrated in the **Section 2.4.2**. This means that the widely used sQSSA equations along with the stationary reactant assumption as given in **Eqs. 2.4.3** will be valid only when *δ*≅ 1. When *δ*< 0 or *δ*> 0, then there is a possibility of multiple steady states and **Eqs. 2.4.3** can capture only the transient first occurring steady state point. One needs to use the velocity equations given in **Eqs. 2.4.2.1-2** to capture the actual second prolonged steady state point. This phenomenon is demonstrated in **Figs. 3** and **Figs. 4**. When *σ* ≅ 1 and *δ<* 1, then the temporal evolution of the enzyme-substrate complex level will show two different time points (denoted as phases **I** and **II** in **Fig. 3A**) at which the trajectory attains maxima where 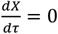 with a local minimum (denoted as phase **III** in **Fig. 3A**) in between these two maxima.

Similarly, when *σ* ≅ 1 and *δ>* 1, then the temporal evolution of the enzyme-inhibitor complex will show up two different time points at which the trajectory attains maximum where 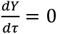 with a local minimum (**Fig. 3B**) in between these two maxima points. Common steady states corresponding to the enzyme-substrate and enzyme inhibitor complexes can occur only when *δ* ≅ 1 as demonstrated in **Figs. 3C**. One can interpret these results as follows. In most of the experimental scenarios, binding of substrate or inhibitor with the respective enzyme will be a diffusion-controlled bimolecular collision process. As a result, one can assume that *k*_1_ ≅ *k*_*i*_ since the size of the substrate and inhibitor are similar which means that *σ* ≅ 1. However, the rate of dissociation and conversion into the respective products will depend on the specific bonding and non-bonding interactions at the protein-ligand interfaces of enzyme-substrate and enzyme-inhibitor complexes. In this context, 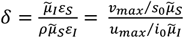 represents the cumulative effects of relative speed of binding, dissociation and conversion into the respective products of enzyme-inhibitor and enzyme-substrate complexes on the overall enzyme catalysis.

Let us define the ratios 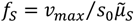 and 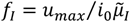 as the acceleration factors with respect to the conversion dynamics of substrate and inhibitor into their respective products. When 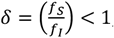, then the speed of conversion of the enzyme-inhibitor complex Y into the product Q will be faster than the conversion speed of enzyme-substrate complex X into the respective product P. The rapid turn-over of the enzyme-inhibitor complex will eventually causes dissociation of already formed enzyme-substrate complex. As a result, when *δ*< 1 and *σ* ≅ 1, then the enzyme-inhibitor complex will show a single steady state and the enzyme-substrate complex will show temporally well-separated two steady states viz. transient primary and prolonged secondary one. This secondary full-fledged steady state can occur only after the depletion of the enzyme-inhibitor complex as shown in **Fig. 3A**. On the other hand, when 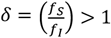 and *σ* ≅ 1, then the rapid turn-over of the enzyme-substrate complex will eventually causes dissociation of already formed enzyme-inhibitor complex. As a result, the enzyme-substrate complex will show a single steady state and the enzyme-inhibitor complex will show temporally well-separated two steady states viz. transient primary and prolonged secondary one. This secondary full-fledged steady state can occur only after the depletion of enzyme-substrate complex as shown in **Fig. 3B**. In term of original variables, 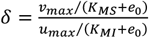 from which one obtains the limiting condition as 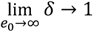 This is a reasonable observation since the fully competitive inhibition scheme will be uncoupled under such limiting condition.

The phase-space behavior of those trajectories described in **Figs. 3** under the conditions that *σ* ≅ 1 and *δ<* 1 or *δ*> 1 over (V, S) and (U, I) spaces are shown in **Figs. 4** and over (V, P) and (U, Q) spaces are shown in **Figs. 5**. When *δ*< 1, then **Figs. 4A-B** and **Figs. 5A-B** suggest that the first occurring steady state point in the evolution of enzyme-substrate complex will be a transient one and it will be observed as a spike in the (V, S) phase-space plot. As a result, we consider the second occurring prolonged steady state as the original steady state with respect to the enzyme-substrate and enzyme-inhibitor complexes. Clearly, the expression obtained from standard QSSA with the reactants stationary assumption as given by **Eqs. 2.4.3** can be used to obtain the steady state velocities only when *σ* ≅ 1 and *δ*≅ 1. When *δ<* 1, then using the standard QSSA one can obtain only the steady state velocity of the enzyme-inhibitor complex as shown in **Figs. 4B** and **5B**. Similarly, when *δ*> 1, then only the steady state velocity associated with enzyme-substrate complex can be obtained as shown in **Figs. 4C** and **5C**. When *δ*≅ 1, then one can obtain the steady state velocities associated with both the enzyme-substrate and enzyme-inhibitor complexes as demonstrated in **Figs. 4E-F** and **5E-F**.

**FIGURE 5.**
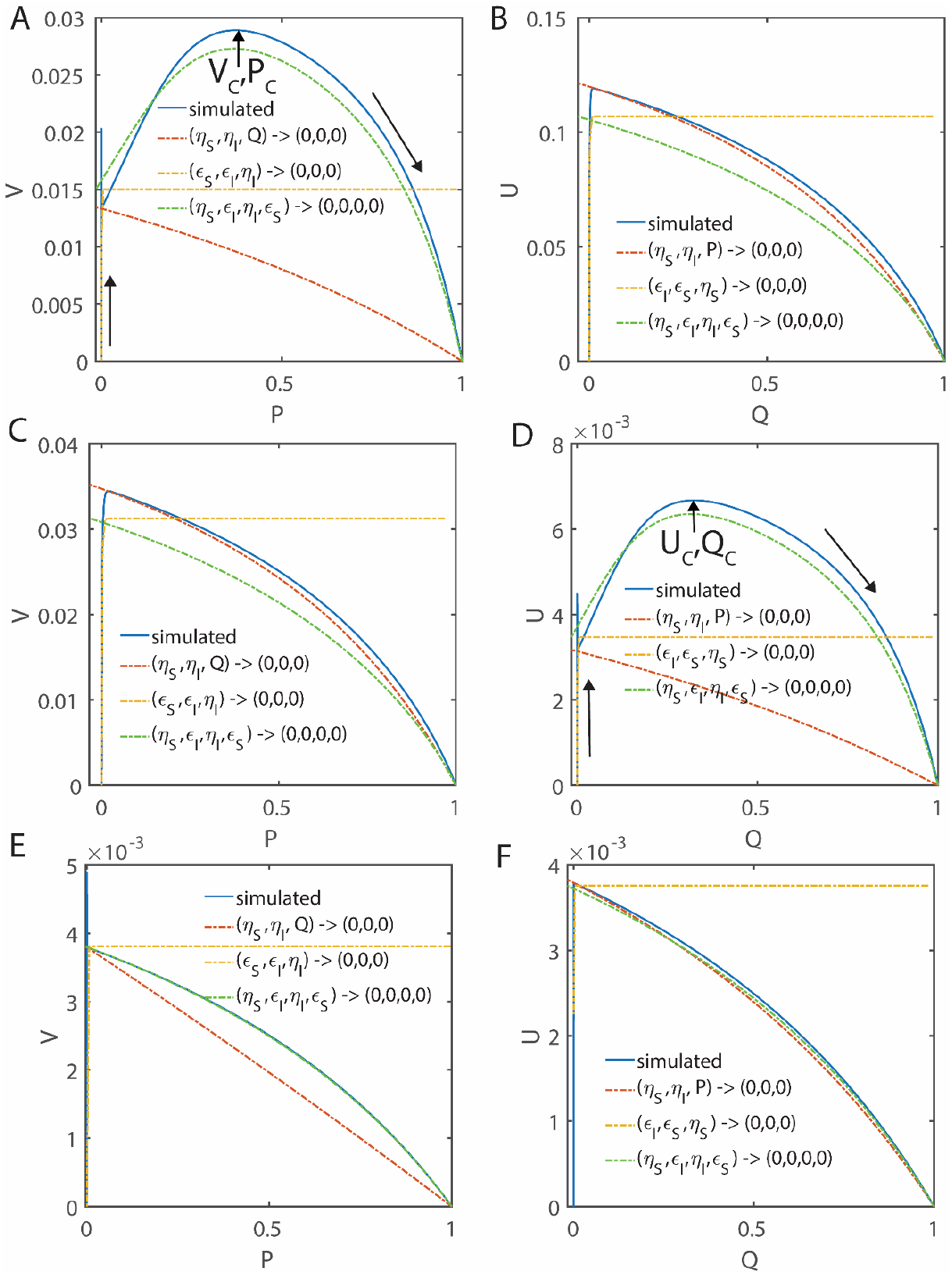
Pre-steady state and post-steady state approximations of the enzyme kinetics with fully competitive inhibition in the velocity-product of substrate (V, P), velocity-product of inhibitor spaces (U, Q) at different values of δ. The phase-space trajectories start at P = 0 and Q = 0, and end at P = 1 and Q = 1 with maxima at the steady state. We considered the approximations of (*V, U*) under the conditions that (*η*_*S*_, *η*_*I*_, *ε*_*S*_, *ε*_*I*_) → (0,0,0,0) which is the refined standard QSSA in both (V, P) and (U, Q) spaces (**Eq. 2.4.18**), (*η*_*S*_, *η*_*I*_, *Q*) → (0,0,0) (**Eq. 2.4.12**) and (*η*_*I*_, *ε*_*S*_, *ε*_*I*_) → (0,0,0) (**Eq. 2.6.10**) corresponding to the post and pre-steady state regimes in the (V, P) space, (*η*_*S*_, *ε*_*S*_, *ε*_*I*_) → (0,0,0) (**Eq. 2.6.23**) and (*η*_*S*_, *η*_*I*_, *P*) → (0,0,0) (**Eq. 2.4.16**) corresponding to the pre- and post-steady state regimes of the (U, Q) space. Using the mass conservation laws V + P + S = 1 and U/ρ + Q + I = 1, V and P can be expressed in terms of S in a parametric form and U and Q can be expressed in in terms of I in a parametric form. Initial conditions for the simulation of **Eqs. 2.2.7-9** are (*S, I, E, X, Y, P, Q*) = (1,1,1,0,0,0,0) at τ = 0. **A-B**. Simulation settings are *η*_*S*_ = 0.002, *ε*_*S*_ = 0.04, *κ*_*S*_ = 0.2, *η*_*I*_ = 0.01, *ε*_*I*_ = 0.06, *κ*_*I*_ = 0.1 and ρ = 3.333, σ = 1, δ = 0.1405, ϒ = 3. **C-D**. Here the simulation settings are *η*_*S*_ = 0.02, *ε*_*S*_ = 0.06, *κ*_*S*_ = 0.1, *η*_*I*_ = 0.003, *ε*_*I*_ = 0.04, *κ*_*I*_ = 0.2 and ρ = 0.225, σ = 1, δ = 9, ϒ = 0.33. **E-F**. Here the simulation settings are *η*_*S*_ = 0.02, *ε*_*S*_ = 0.06, *κ*_*S*_ = 8.1, *η*_*I*_ = 0.003, *ε*_*I*_ = 0.04, *κ*_*I*_ = 1.2 and ρ = 0.225, σ = 1, δ = 1.013, ϒ = 4.5.

Irrespective of the values of δ, the expressions corresponding to the pre-steady state dynamics in the (V, S) space under the conditions that (*η*_*I*_,*ε*_*S*_,*ε*_*I*)_ → (0,0,0) (**Eqs. 2.6.10-11**) and in the (U, I) space under the conditions that (*η*_*S*_,*ε*_*S*_,*ε*_*I*)_ → (0,0,0) (**Eqs. 2.6.23-24**) can approximate the simulated trajectory very well as demonstrated in **Figs. 4A-F** and **5A-F**. Interestingly, when *δ*< 1, then the approximation under the conditions that (*η*_*S*_,*η*_*I*_,*P*) → (0,0,0) that is given by **Eq. 2.4.16** accurately predicts the post-steady state reaction velocity associated with the enzyme-inhibitor complex in the (U, I) space as shown in **Fig. 4B** and **5B**. Similarly, when *δ*> 1, then the approximation under the conditions that (*η*_*S*_,*η*_*I*_,*Q*) → (0,0,0) that is given in **Eq. 2.4.12** can accurately predicts the post-steady state reaction velocity associated with the enzyme-substrate complex in the (V, S) space as shown in **Fig. 4C** and **5C**.

### 4.3. Minimization of error in the sQSSA over (V, S) and (U, I) spaces

The overall error associated with the standard QSSA with stationary reactants assumption of the fully competitive enzyme kinetics used in the literature over (V, S) and (U, I) spaces at different values of (*ε*_*S*_,*ε*_*I*)_ and other parameters are shown in **Figs. 6**. Similarly, the error characteristics of the refined form of standard QSSA are shown in **Figs. 7-8. Figs. 6** clearly demonstrate the poor performance of sQSSA given in **Eqs. 2.4.2-3** which are widely used in the literature to obtain the kinetic parameters of fully competitive inhibition systems especially when (*k*_*S*_,*k*_*I*_) ≪ (1,1). We summarize the following essential conditions for the validity of the sQSSA given in **Eqs. 2.4.2-3** and the refined form of sQSSA given in **Eqs. 2.4.19** with stationary reactant assumptions (S, I) = (1,1). Apparently, these conditions are mandatory to minimize the error in various sQSSAs.

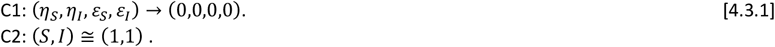

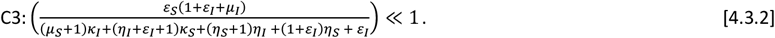

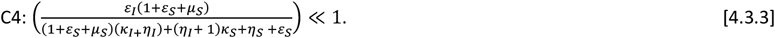

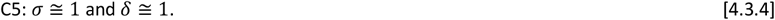

**FIGURE 6.**
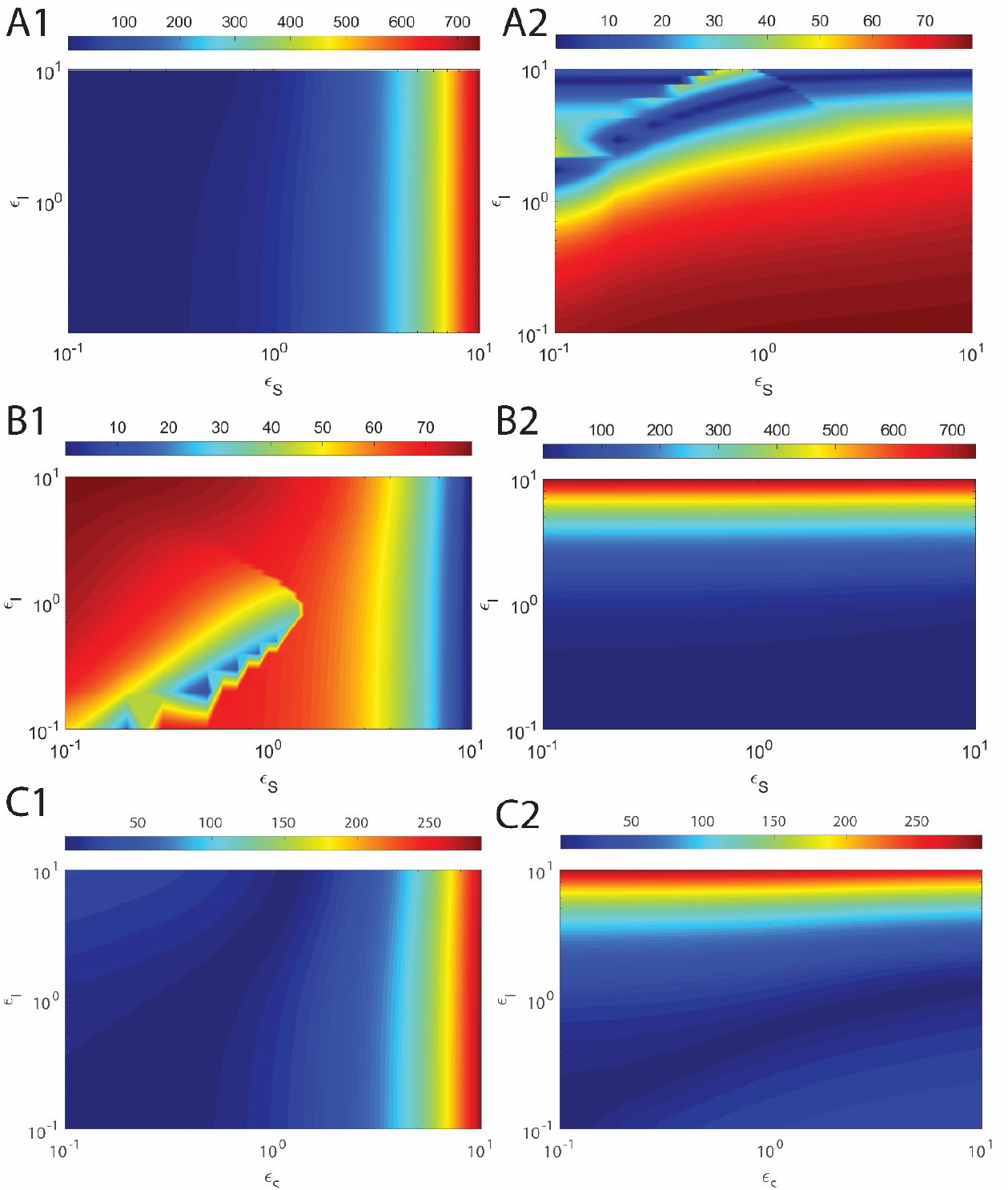
Error associated with the sQSSA with stationary reactant assumption of the fully competitive enzyme kinetics over the velocity-substrate (V, S), velocity-inhibitor spaces (U, I) at different values of *ε*_*S*_, *ε*_*I*_. Here δ will vary with respect to each iteration. We considered the error in the approximations of the reaction velocities (*V, U*) under the conditions that (*η*_*S*_, *η*_*I*_, *ε*_*S*_, *ε*_*I*_) → (0,0,0,0) and stationary reactant assumption as defined in **Eqs. 2.4.2**. The error was computed as error (%) = 100 |steady state velocities from simulation – approximated velocities| / steady state velocities from simulation. Here the simulation settings are *η*_*S*_ = 0.02, *η*_*I*_ = 0.01 and σ = 1. With these settings, upon fixing σ one finds that 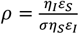 and 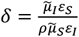 as defined in **Eqs. 2.4.22. A1, B1, C1**. Error % in the standard QSSA of V. **A2, B2, C2**. Error % in QSSA of U. **A1-2**. *κ*_*S*_ = 0.1, *κ*_*I*_ = 1. **B1-2**. *κ*_*S*_ = 1, *κ*_*I*_ = 0.1. **C1-2**. *κ*_*S*_ = 1, *κ*_*I*_ = 1.

**FIGURE 7.**
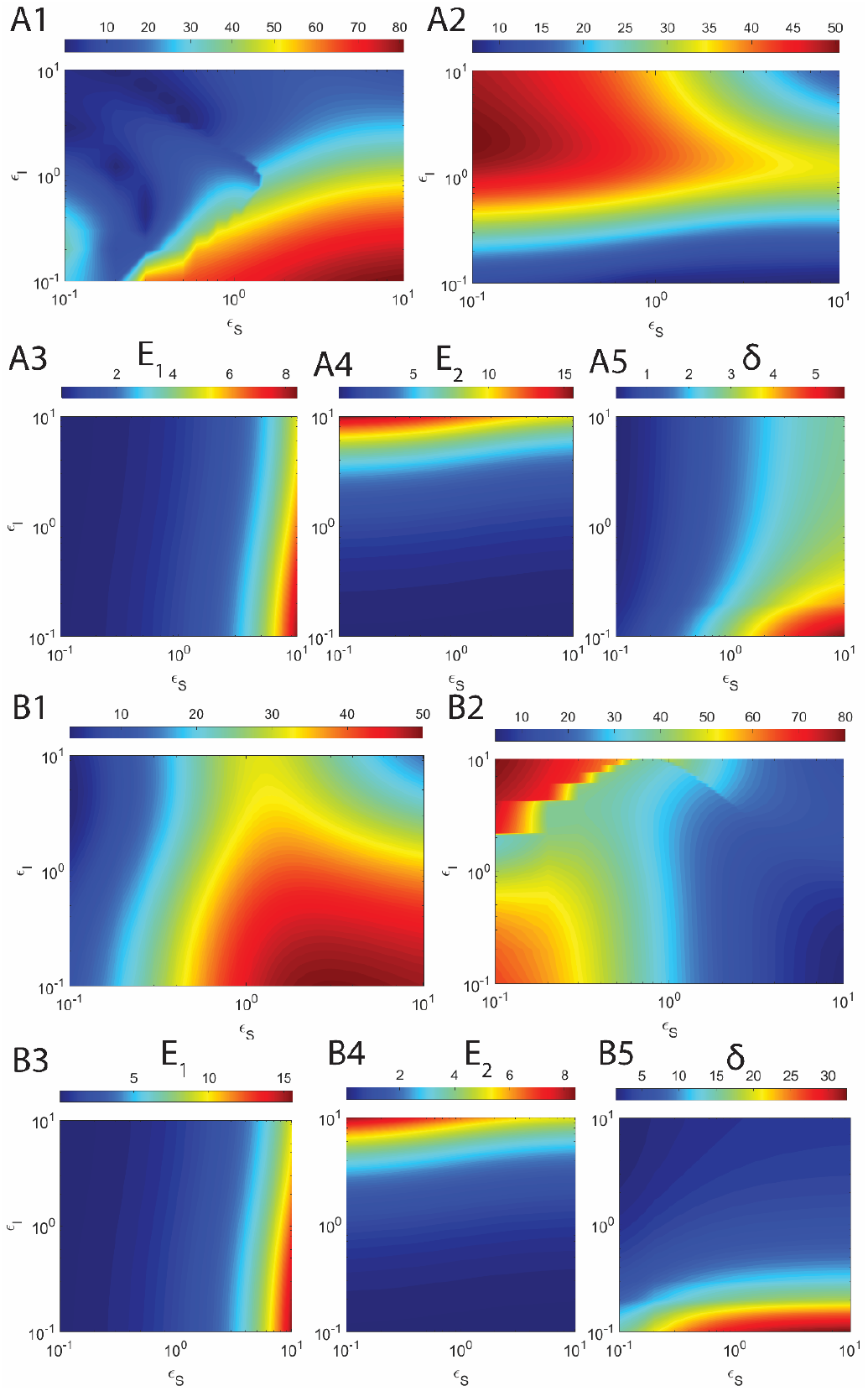
Error associated with the refined form of sQSSA with stationary reactant assumption of the fully competitive enzyme kinetics in the velocity-substrate (V, S), velocity-inhibitor spaces (U, I) at different values of *ε*_*S*_, *ε*_*I*_ under the conditions that *κ*_*S*_ ≠ *κ*_*I*_. Here E_1_, E_2_ and δ will vary with respect to each iteration. We considered the error in the approximations of the reaction velocities (*V, U*) under the conditions that (*η*_*S*_, *η*_*I*_, *ε*_*S*_, *ε*_*I*_) → (0,0,0,0) various limiting conditions as defined in **Eqs. 2.4.19** with (S, I) = (1, 1). The error was computed as error (%) = 100 |steady state velocities from simulation – approximated velocities| / steady state velocities from simulation. Here the simulation settings for **A1-5** are *η*_*S*_ = 0.02, *κ*_*S*_ = 1, *η*_*I*_ = 0.01, *κ*_*I*_ = 0.1 and σ = 1. Simulation settings for **B1-5** are *η*_*S*_ = 0.02, *κ*_*S*_ = 0.1, *η*_*I*_ = 0.01, *κ*_*I*_ = 1 and σ = 1. With these settings, upon fixing σ one finds that 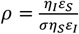 and 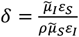 as defined in **Eqs. 2.4.22** along with the inequality conditions E_1_, and E_2_ as defined in **Eqs. 2.7.5-6. A1, B1**. Error % in the QSSA of V. **A2, B2**. Error % in the QSSA of U. **A3, B3**. E_1_ (**Eq. 2.7.5**). **A4, B4**. E_2_ (**Eq. 2.7.6**). **A5, B5**. *δ* as defined in **Eqs. 2.4.22**.

The condition C1 ensure the occurrence of similar steady state timescales associated with the enzyme-substrate and enzyme-inhibitor complexes. C2 is the stationary reactant assumption that is required to approximate the unknown steady state substrate and inhibitor levels. C3 and C4 (following from **Eqs. 2.7.5-6**) are required to minimize the deviations occurring in the pre-steady state regime. C5 is required to avoid the complex multiple steady state dynamics of enzyme-substrate and enzyme-inhibitor complexes. The error levels of the refined standard QSSA given in **Eqs. 2.4.19** are demonstrated in **Figs. 7** and **Figs. 8. Figs. 7** show the error levels under the conditions that *k*_*S*_ ≠ *k*_*I*_ and **Figs. 7** demonstrate the error levels when *k*_*S*_ = *k*_*I*_. **Figs. 7** and **Figs. 8** clearly show the error control capability of the refined QSSA expressions given by **Eqs. 2.4.19** in estimating the steady state reaction velocities over **Eqs. 2.4.2-3**. Remarkably, there is a strong correlation between the error levels in the estimated steady state reaction velocities and *δ*. The error level associated with the steady state velocity of the enzyme-substrate complex is positively correlated with *δ*. On the other hand, the error levels associated with the steady state velocity of the enzyme-inhibitor complex seem to be negatively correlated with *δ*. These means that to obtain the accurate estimate of the steady state velocity of enzyme-substrate complex using **Eqs. 2.4.3**, one needs to set *δ*> 1. To obtain the accurate estimate of the steady state velocity of the enzyme-inhibitor complex, one needs to set *δ*< 1. On the overall basis, we find that the conditions C1, C5 are critical to minimize the error in the refined standard QSSA.

**FIGURE 8.**
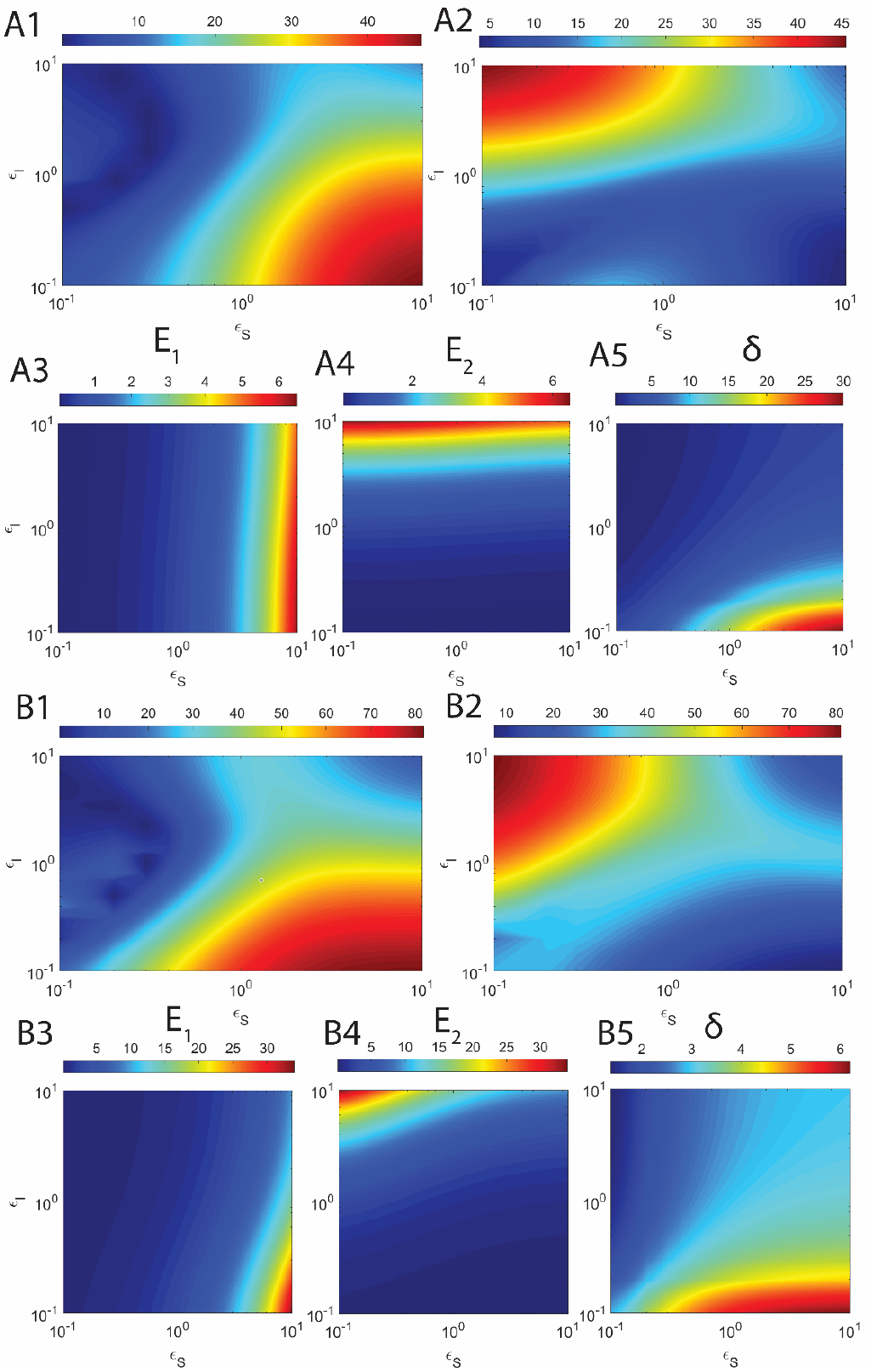
Error associated with the refined form of sQSSA with stationary reactant assumption of the fully competitive enzyme kinetics in the velocity-substrate (V, S), velocity-inhibitor spaces (U, I) at different values of *ε*_*S*_, *ε*_*I*_ under the condition that *κ*_*S*_ = *κ*_*I*_. Here E_1_, E_2_ and δ will vary with respect to each iteration. We considered the error in the approximations given in **Eqs. 2.4.19** with (S, I) = (1, 1). The error was computed as error (%) = 100 |steady state velocities from simulation – approximated velocities| / steady state velocities from simulation. Here the simulation settings for **A1-5** are *η*_*S*_ = 0.02, *κ*_*S*_ = *κ*_*I*_ = 1, *η*_*I*_ = 0.01 and σ = 1. Similar simulation settings for **B1-5** with *κ*_*S*_ = *κ*_*I*_ = 0.1. With these settings, upon fixing the value of σ one finds that 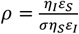 and 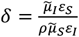 as defined in **Eqs. 2.4.22** along with the inequality conditions E_1_, and E_2_ as defined in **Eqs. 2.7.5-6. A1, B1**. Error % in QSSA of V. **A2, B2**. Error % in QSSA of U. **A3, B3**. E_1_ (**Eq. 2.7.5**). **A4, B4**. E_2_ (**Eq. 2.7.6**). **A5, B5**. *δ* as defined in **Eqs. 2.4.22**.

### 4.4. The ϕ-approximations

The performance of ϕ-approximations described by the set of coupled linear ODEs given by **Eqs. 2.5.1.1-2** over the (V, I, S) and (U, I, S) spaces is demonstrated along with the sQSSA trajectories in **Figs. 9**. These parametric solutions were generated with τ as the parameter as given in the integral solutions of **Eqs. 2.5.1.1-2** in **Appendix A**. Clearly, the ϕ-approximations accurately fit both the pre- and post-steady state regimes especially at large values of (*ε*_*S*_,*ε*_*I*_). Whereas, the sQSSAs work only in the post steady state regime. Behavior of ϕ-approximations in the (V, P, Q) and (U, P, Q) spaces are demonstrated in **Figs. 10** and the performance of ϕ-approximations in the (V, U) space at various parameter values are demonstrated in **Figs. 11**.

**FIGURE 9.**
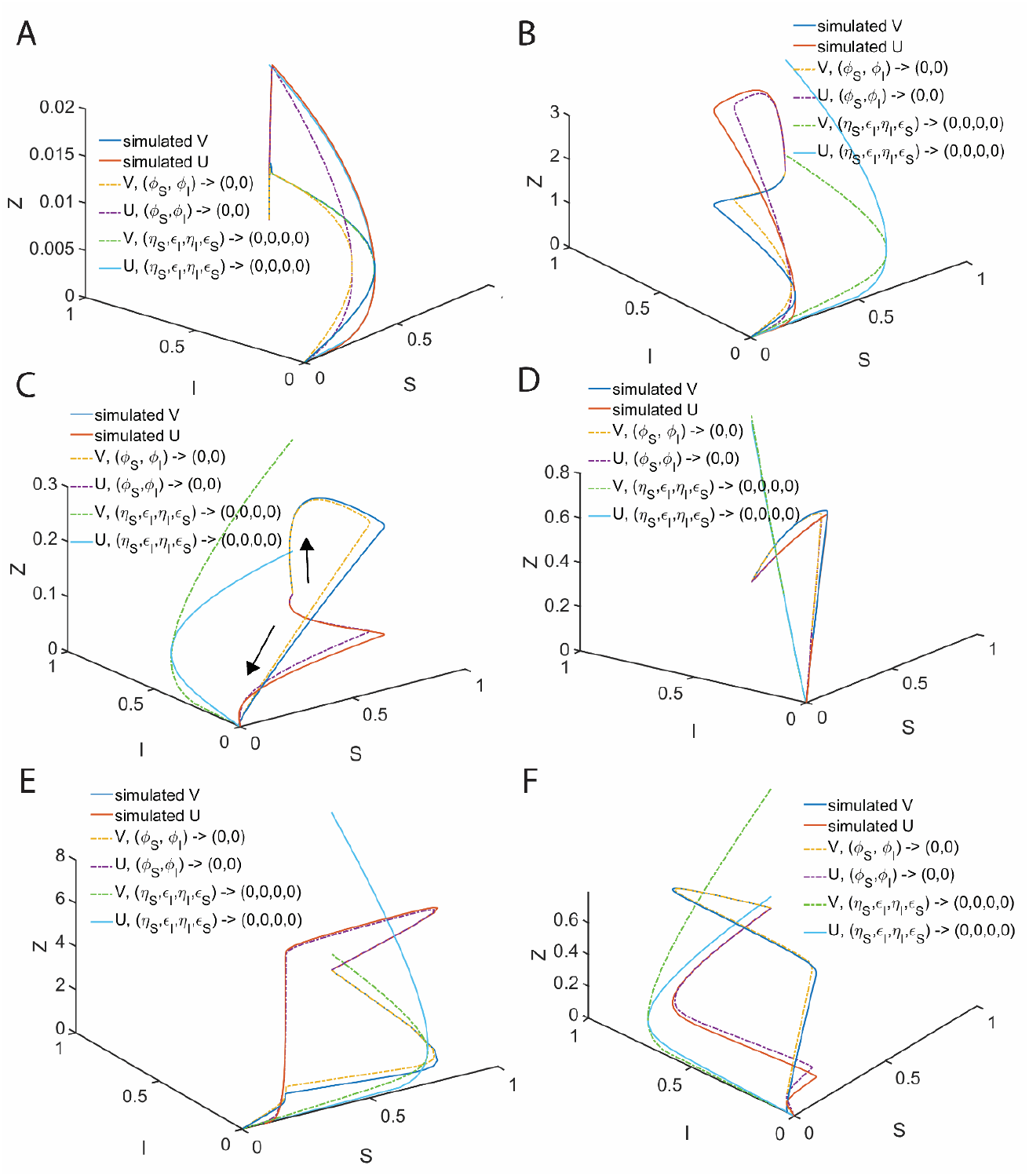
Approximate solutions in the velocity-substrate-inhibitor spaces (V, I, S) and (U, I, S). We considered the approximations of the reaction velocities (Z = V and U) under the conditions (*η*_*S*_, *η*_*I*_, *ε*_*S*_, *ε*_*I*_) → (0,0,0,0) which is the refined form of standard QSSA as given in **Eqs. 2.4.19** and **Eqs. 2.6.12** (for the relationship between S and I) and under the conditions (*ϕ*_*I*_, *ϕ*_*S*_) → (0,0) as given by the solutions of the coupled approximate linear ODEs **Eqs. 2.5.1.1-2** as given in **Appendix A** in a parametric form where τ acts as the parameter. Here the common initial conditions for the numerical simulation of **Eqs. 2.2.7-9** are (*S, I, E, X, Y, P, Q*) = (1,1,1,0,0,0,0) at τ = 0 and other simulation settings are *η*_*S*_ = 0.06, *κ*_*S*_ = 8.1, *η*_*I*_ = 0.03, *κ*_*I*_ = 1.2. σ = 1 for (**A-D**), σ = 0.1 for **E** and σ = 10 for **F. A**. *ε*_*S*_ = 0.08, *ε*_*I*_ = 0.04, ρ = 1, δ = 0.3083, ϒ = 3.375. **B**. *ε*_*S*_ = 13.8, *ε*_*I*_ = 0.4, ρ = 17.25, δ = 0.1485, ϒ = 0.1957. **C**. *ε*_*S*_ = 3.8, *ε*_*I*_ = 20.4, ρ = 0.093, δ = 3.617, ϒ = 36.24. **D**. *ε*_*S*_ = 33.8, *ε*_*I*_ = 20.4, ρ = 0.8284, δ = 1.031, ϒ = 4.074. **E**. *ε*_*S*_ = 33.8, *ε*_*I*_ = 20.4, ρ = 8.284, δ = 0.1031, ϒ = 0.4074. **F**. *ε*_*S*_ = 33.8, *ε*_*I*_ = 20.4, ρ = 0.0828, δ = 10.31, ϒ = 40.74.

**FIGURE 10.**
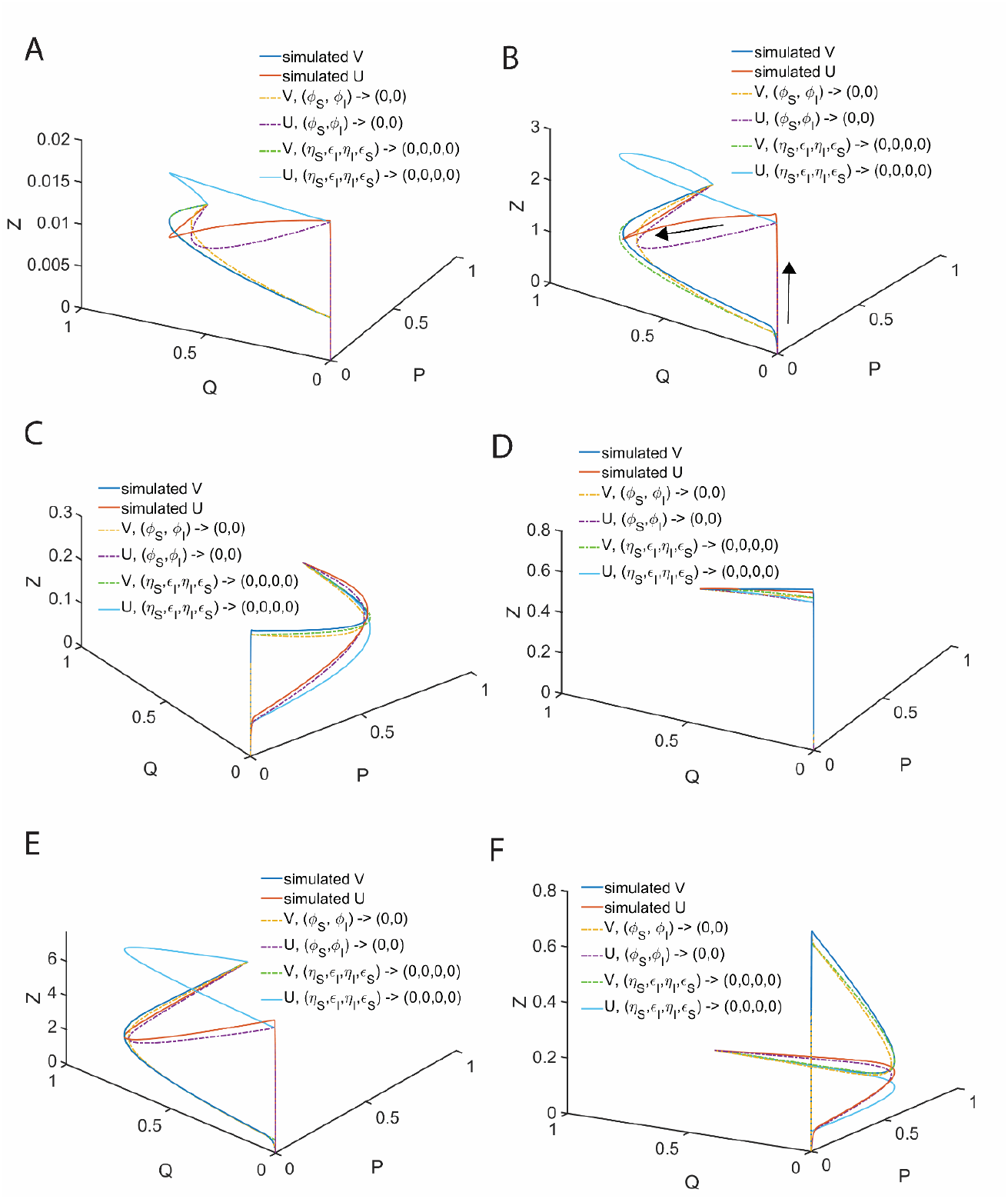
The φ-approximations of the fully competitive enzyme kinetics in the velocity-products spaces (V, P, Q) and (U, P, Q) at different values of δ. The phase-space trajectories start at P = 0 and Q = 0, and end at P = 1 and Q = 1 with maxima at the steady state. We considered the φ-approximations of (*V, U*)which are the solutions of **Eqs. 2.5.1.1-2** as given in **Appendix A** in a parametric form where τ act as the parameter and standard QSSA solutions under the conditions that (*η*_*S*_, *η*_*I*_, *ε*_*S*_, *ε*_*I*_) → (0,0,0,0) in a parametric form where S acts as the parameter as given in **Eqs. 2.4.19** and **Eqs. 2.6.12**. Common initial conditions for the numerical simulation of **Eqs. 2.2.7-9** are (*S, I, E, X, Y, P, Q*) = (1,1,1,0,0,0,0) at τ = 0 and other simulation settings are *η*_*S*_ = 0.06, *κ*_*S*_ = 8.1, *η*_*I*_ = 0.03, *κ*_*I*_ = 1.2. σ = 1 for (**A-D**), σ = 0.1 for **E** and σ = 10 for **F. A**. *ε*_*S*_ = 0.08, *ε*_*I*_ = 0.04, ρ = 1, δ = 0.3083, ϒ = 3.375. **B**. *ε*_*S*_ = 13.8, *ε*_*I*_ = 0.4, ρ = 17.25, δ = 0.1485, ϒ = 0.1957. **C**. *ε*_*S*_ = 3.8, *ε*_*I*_ = 20.4, ρ = 0.093, δ = 3.617, ϒ = 36.24. **D**. *ε*_*S*_ = 33.8, *ε*_*I*_ = 20.4, ρ = 0.8284, δ = 1.031, ϒ = 4.074. **E**. *ε*_*S*_ = 33.8, *ε*_*I*_ = 20.4, ρ = 8.284, δ = 0.1031, ϒ = 0.4074. **F**. *ε*_*S*_ = 33.8, *ε*_*I*_ = 20.4, ρ = 0.0828, δ = 10.31, ϒ = 40.74.

**FIGURE 11.**
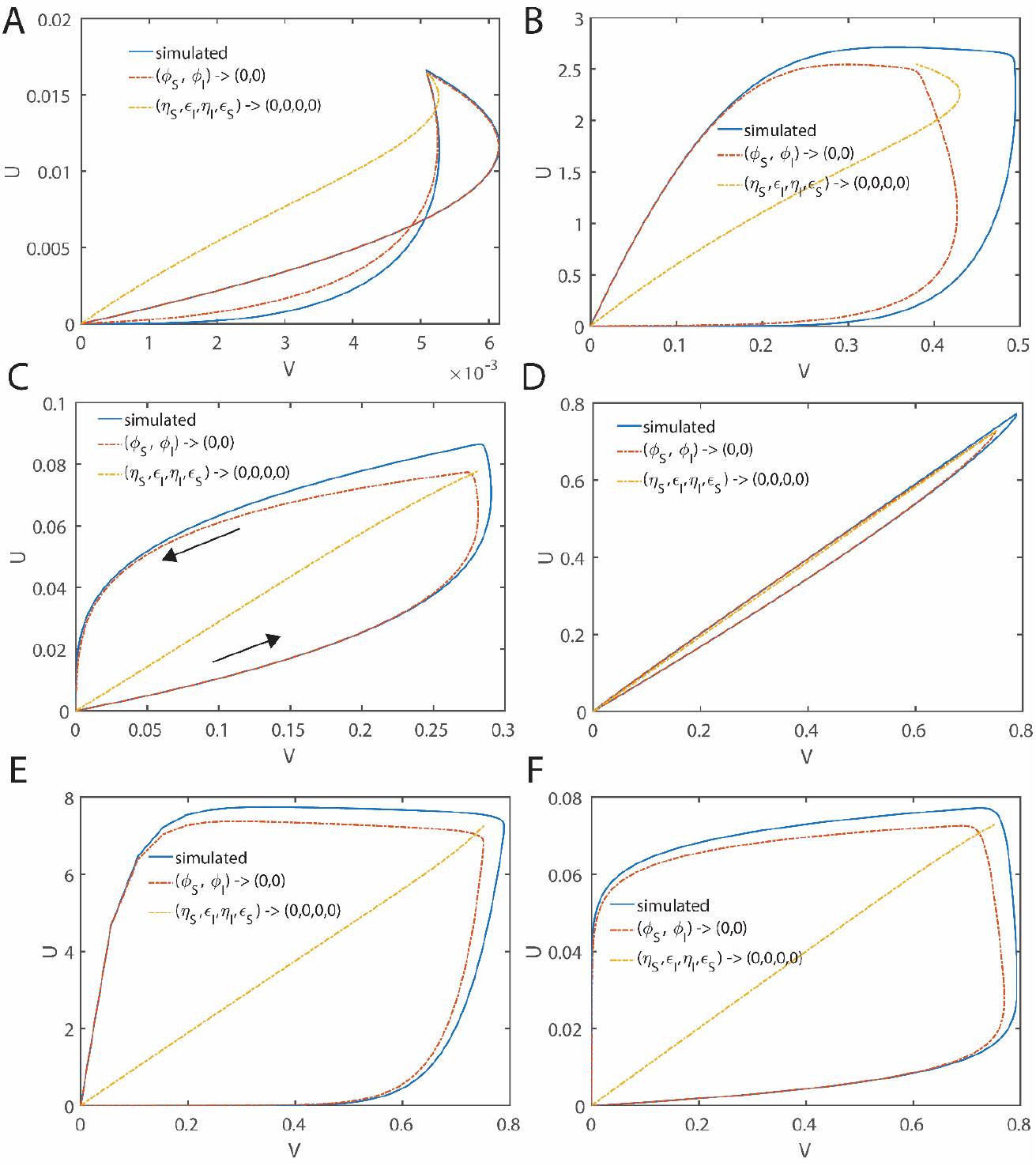
The φ-approximations of the fully competitive enzyme kinetics in the velocity spaces (V, U) at different values of δ. We considered the φ-approximations which are the solutions of **Eqs. 2.5.1.1-2** as given in **Appendix A** in a parametric form where τ act as the parameter and standard QSSA solutions for (*V, U*) under the conditions that (*η*_*S*_, *η*_*I*_, *ε*_*S*_, *ε*_*I*_) → (0,0,0,0) in a parametric form where S acts as the parameter as given in **Eqs. 2.4.19** and **Eqs. 2.4.22**. The trajectory in the (V, U) space starts at (V, U) = (0, 0) and ends at (V, U) = (0, 0). Arrow in **C** indicates the direction of the trajectory evolution. Common initial conditions for the numerical simulation of **Eqs. 2.2.7-9** are (*S, I, E, X, Y, P, Q*) = (1,1,1,0,0,0,0) at τ = 0 and other simulation settings are *η*_*S*_ = 0.06, *κ*_*S*_ = 8.1, *η*_*I*_ = 0.03, *κ*_*I*_ = 1.2. σ = 1 for (**A-D**), σ = 0.1 for **E** and σ = 10 for **F. A**. *ε*_*S*_ = 0.08, *ε*_*I*_ = 0.04, ρ = 1, δ = 0.3083, ϒ = 3.375. **B**. *ε*_*S*_ = 13.8, *ε*_*I*_ = 0.4, ρ = 17.25, δ = 0.1485, ϒ = 0.1957. **C**. *ε*_*S*_ = 3.8, *ε*_*I*_ = 20.4, ρ = 0.093, δ = 3.617, ϒ = 36.24. **D**. *ε*_*S*_ = 33.8, *ε*_*I*_ = 20.4, ρ = 0.8284, δ = 1.031, ϒ = 4.074. **E**. *ε*_*S*_ = 33.8, *ε*_*I*_ = 20.4, ρ = 8.284, δ = 0.1031, ϒ = 0.4074. **F**. *ε*_*S*_ = 33.8, *ε*_*I*_ = 20.4, ρ = 0.0828, δ = 10.31, ϒ = 40.74.

### 4.5. Partial competitive inhibition

Most of the inhibitory drug molecules and the respective experimental systems are partial competitive ones following the **Scheme B** of **Fig. 1**. These inhibitor molecules competitively bind the active site of the target enzyme against the natural substrate and form reversible dead-end complexes. Upon complete depletion of the substrate level, enzyme-inhibitor complex attains the equilibrium state as demonstrated in **Fig. 12A**. Similar to the fully competitive inhibition scheme, depending on the steady state timescales of the enzyme-substrate and enzyme-inhibitor complexes, partial completive inhibition scheme can also exhibit a complex behavior as demonstrated in **Fig. 12A**. Sample trajectories over (V, S), (Y, S), (I, S), (V, I, S) and (V, P, S) spaces are shown in **Figs. 12B-D** along with the pre- and post-steady state approximations under the conditions that (*ε*_*S*_,*X*_*I*_,*ε*_*I*_) → (0,0,0) and (*η*_*S*_,*X*_*I*)_ → (0,0) respectively. When (*η*_*S*_,*X*_*I*)_ → (0,0), then one can approximate the reaction velocity associated with the enzyme-substrate compelx as 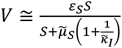 in the (V, S) space and the level of enzyme-inhibitor compelx as 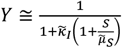 in the (Y, S) space as shown in **Eqs. 2.9.2.2**. In the derivation of these equations, we have applied stationary reactant assumption on the inhibitor as *I* ≅ 1. Upon applying the stationary reactant assumption on the substrate as *S* ≅ 1, one finally arives at 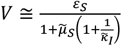 under the conditions that (*η*_*S*_,*X*_*I*_,*ε*_*S*_,*ε*_*I*)_ → (0,0,0,0). In terms of the original variables, this refined equation for the partial competitive inhbition scheme can be writen as 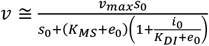 that is given in **Eqs. 2.9.3.17**. Remarkably, under the conditions that (*η*_*S*_,*X*_*I*_) → (0,0) the post-steady state (V, S) space approximation given by **Eq. 2.9.4.7** seems to be more accurate than the approximation given by **Eqs. 2.9.2.2**.

**FIGURE 12.**
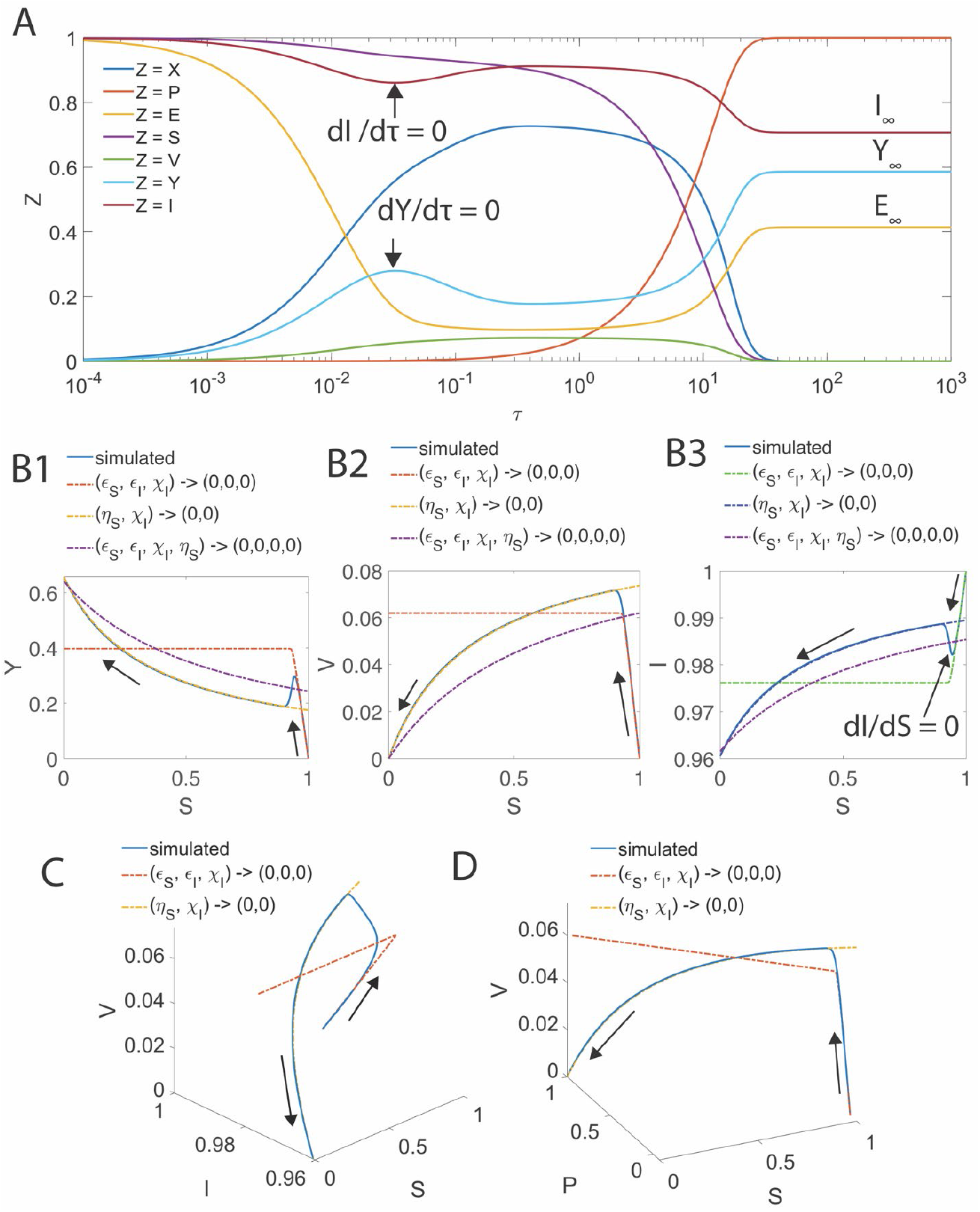
Pre- and post-steady state approximations of the enzyme kinetics with partial competitive inhibition. Here the simulation settings are *η*_*S*_ = 0.02, *ε*_*S*_ = 0.1, *κ*_*S*_ = 0.1, *χ*_*I*_ = 0.03, *ε*_*I*_ = 0.06, *κ*_*I*_ = 0.5. Common initial conditions for the numerical simulation of **Eqs. 2.9.5-7** are (*S, I, E, X, Y, P*) = (1,1,1,0,0,0) at τ = 0. Post-steady state approximations were generated under the conditions that (*η*_*S*_, *χ*_*I*_) → 0 and the pre-steady state approximations were computed under the conditions that (*ε*_*S*_, *χ*_*I*_, *ε*_*I*_) → (0,0,0). **A**. Simulation trajectories of (*S, I, E, X, Y, P*). Clearly, (*I, E, Y*) ends at the equilibrium states (*I*_∞_, *E*_∞_, *Y*_∞_) upon complete depletion of the substrate. When the steady state timescales of X and Y are different, then Y will exhibit a steady state where 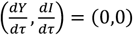. **B1-3, C** and **D**. Simulated trajectories along with the in the pre- and post-steady state approximations. Post steady state approximations under the conditions that (*η*_*S*_, *χ*_*I*_, *ε*_*S*_, *ε*_*I*_) → (0,0,0,0) were generated using **Eqs. 2.9.2.2. B1**. (Y, S) space trajectory and approximations are computed using **Eqs. 2.9.3.10** and **2.9.4.9** for the pre and post-steady state regimes respectively. **B2**. (V, S) space trajectory with approximations using **Eqs. 2.9.3.8** and **2.9.4.7** and **2.9.2.2** corresponding to the pre and post steady state regimes. **B3**. (S, I) space trajectory and approximations using the mass conservation law *I* = 1 - *ε*_*I*_ *Y* (**Eqs. 2.9.3.10** and **2.9.4.9** with *S* ∈ [0,1] as the parameter). When 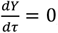, then one finds that 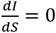, representing a local minimum in the (I, S) space. **C**. (V, S, I) space approximations (**Table 3** for parametric representations). **D**. (V, P, S) space approximations (**Table 3**).

In the sQSSA expressions which are generally used in the literature, the term *e*_0_ will not be added up to *K*_*MS*_ and *K*_*DI*_. Similar to **Eqs. 2.6.13** and **2.6.26, Eq. 2.9.3.17** will be valid over wide range of parameters (*ε*_*S*_,*ε*_*I*)_ as demonstrated in **Figs. 12B-D**. As in **Eqs. 4.3.1-5**, the error in the steady state reaction velocity **Eqs. 2.9.3.17** can be minimized using the following conditions.

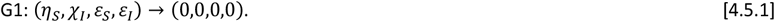

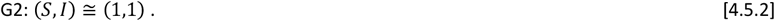

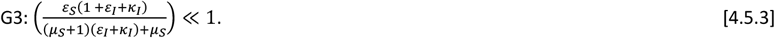

Here the condition G1 is required for the occurrence of common steady states with respect to both the enzyme-substrate and enzyme-inhibitor complexes. The condition G2 is the stationary reactant assumption that is required to approximate the unknown steady state levels of (S, I). The condition G3 ensures the occurrence of minimal error in the pre-steady state regime. The overall error in various steady state approximations of the partial competitive inhibition scheme at different parameter settings are demonstrated in **Figs. 13**. We considered the sQSSA given by **Eqs. 2.9.10**, refined form of sQSSA given by **Eqs. 2.9.3.16** and the error in **Eqs. 2.9.4.7** with S = 1. Results clearly suggest that the approximations given by **Eqs. 2.9.10** and **2.9.3.16** works well when (*k*_*I*_,*k*_*I*_) ≫ (1,1). When (*k*_*I*_, *k*_*I*_) ≪ (1,1), then the approximation given by **Eqs. 2.9.4.7** predicts the steady state velocity well.

**FIGURE 13.**
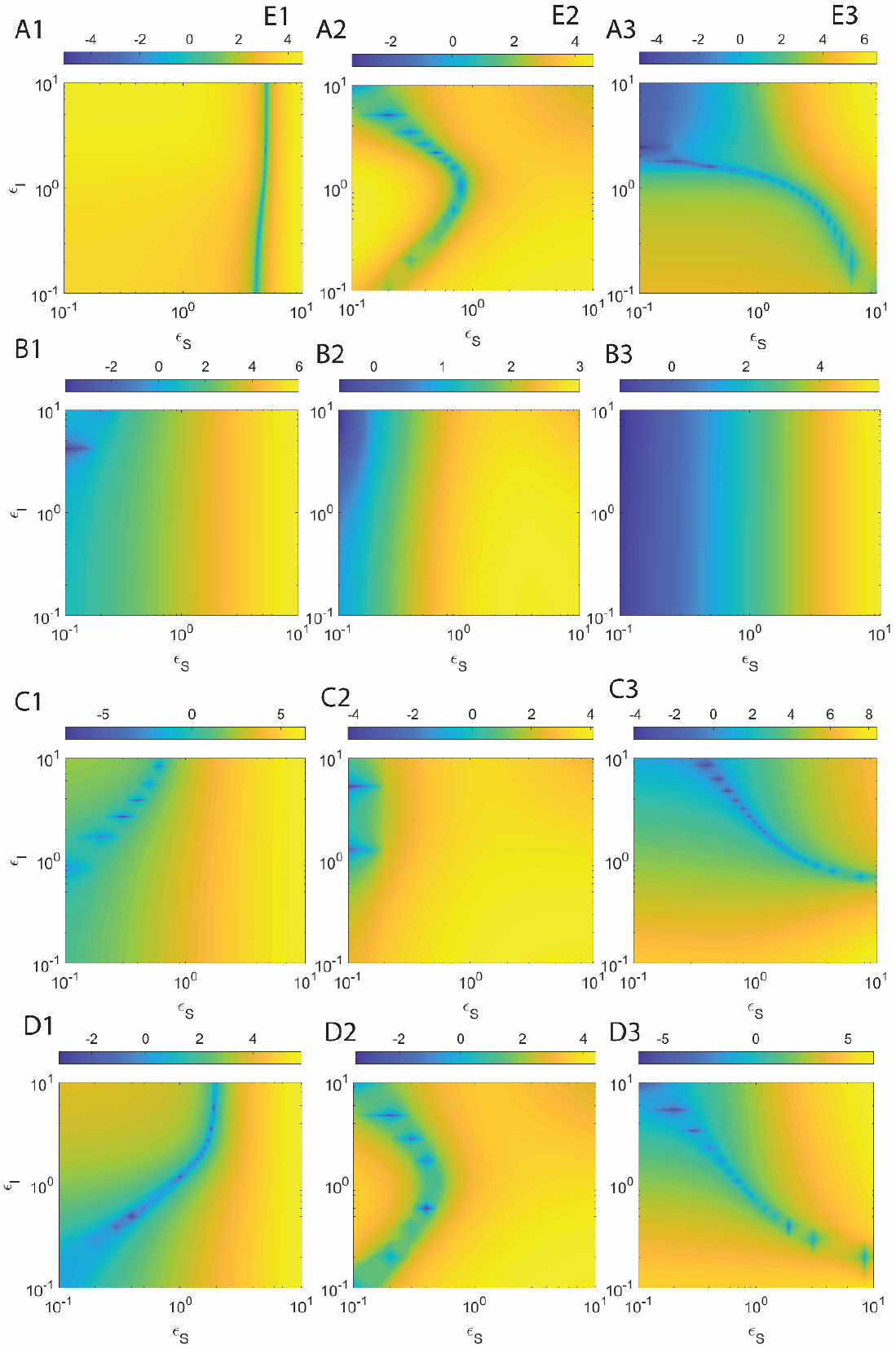
Error associated with various steady state approximations along with the stationary reactant assumption corresponding to the enzyme kinetics with partial competitive inhibition in the velocity-substrate (V, S) space at different values of *ε*_*S*_, *ε*_*I*_. Here E_1_ (A1, B1, C1 and D1, error in sQSSA given by **Eqs. 2.9.10**), E_2_ (A2, B2, C2 and D2, error in the refined form of sQSSA given in **Eqs. 2.9.3.16**) and E_3_ (A3, B3, C3 and D3, error in **Eqs. 2.9.5.5**) are logarithm of percentage errors. The computed error (%) = 100 |steady state velocities from simulation – approximated velocities| / steady state velocities from simulation. **A1-3**. *η*_*S*_ = 0.02, *κ*_*S*_ = 0.001, *κ*_*I*_ = 0.005, *η*_*I*_ = 0.01. **B1-3**. *η*_*S*_ = 0.02, *κ*_*S*_ = 1, *κ*_*I*_ = 5, *η*_*I*_ = 0.01. **C1-3**. *η*_*S*_ = 0.02, *κ*_*S*_ = 0.1, *κ*_*I*_ = 0.5, *η*_*I*_ = 0.01. **D1-3**. *η*_*S*_ = 0.02, *κ*_*S*_ = 0.1, *κ*_*I*_ = 0.1, *η*_*I*_ = 0.01.

### 4.6. Substrate-inhibitor (S, I) space dynamics

Remarkably, when the timescales associated with the steady states of enzyme-substrate (X) and enzyme-inhibitor (Y) complexes are not the same, then one can show that there exists a regime in the (I, S) space at which 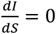 (**Figs. 14**). Since *I* = 1 − *ε Y* and *S* = 1 − *ε X* − *P* for the partial competitive inhibition scheme, one finds that 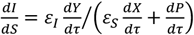 which will be zero at the steady state of Y that occurs at the time point τ_*CY*_ where 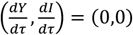. Since the steady state timescale of X is different from the steady state timescale of 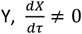 at τ_*CY*_. One should note that 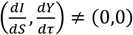 both under the conditions τ = 0 and τ → ∞ where Y approaches its equilibrium value. Building up of the product P will be a monotonically increasing function of time so that 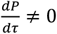 over the entire timescale regime except at τ = 0 and τ → ∞ (**Figs. 12A, 12B3**, and **14C-D**). Further, 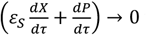 when τ → ∞ which means that when τ → ∞, then 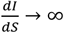 in case of partial competitive inhibition.

**FIGURE 14.**
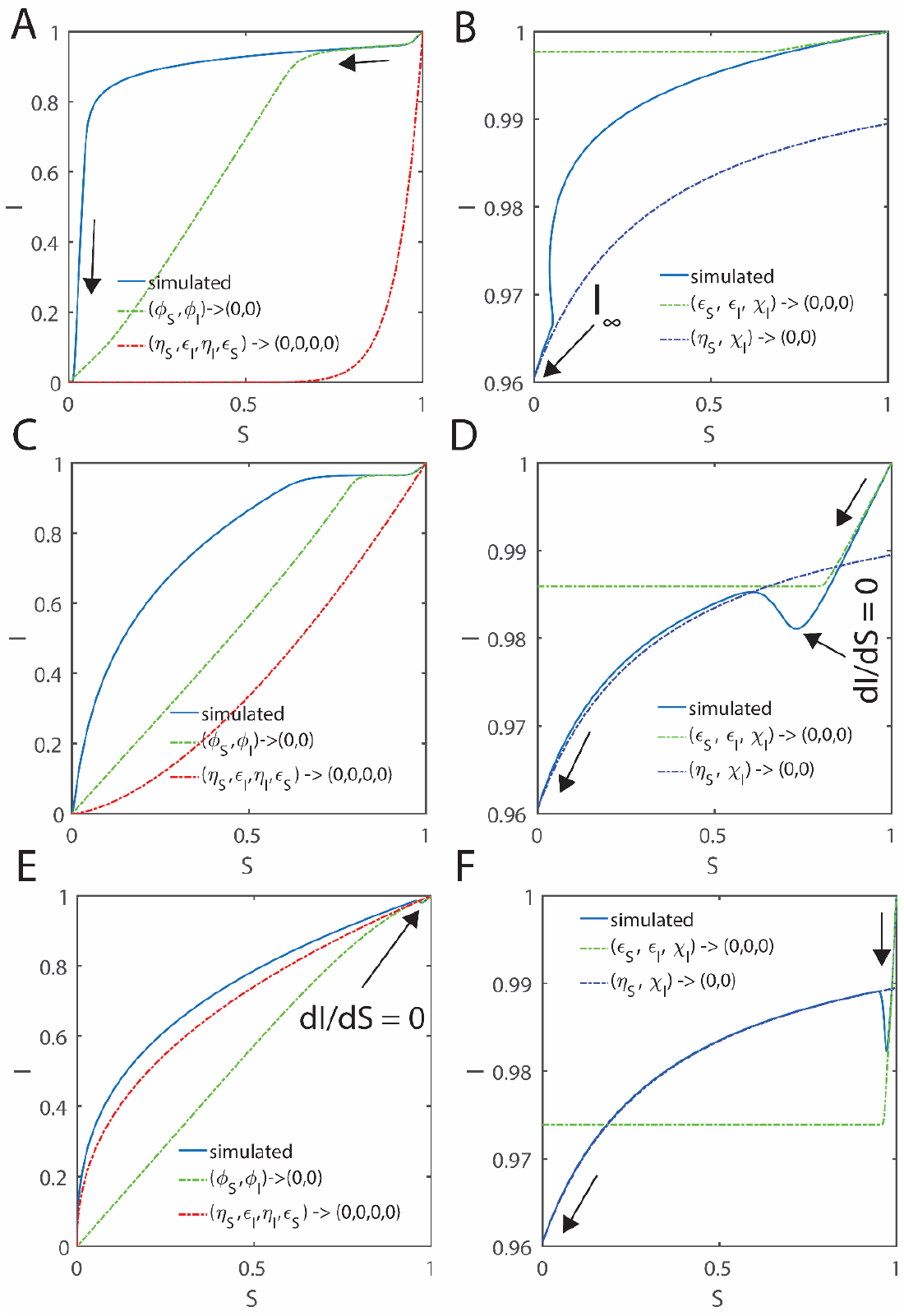
Pre- and post-steady state approximations in the inhibitor-substrate space for the enzyme kinetics with fully and partial competitive inhibition. Common initial conditions for the simulation of fully competitive inhibition **Eqs. 2.2.7-9** are (*S, I, E, X, Y, P, Q*) = (1,1,1,0,0,0,0) and for partial competitive inhibition **Eqs. 2.9.5-7** are (*S, I, E, X, Y, P*) = (1,1,1,0,0,0) at τ = 0. For the fully competitive inhibition scheme (**A, C, E** where trajectories start at (I, S) = (1,1) and end at (I, S) = (0,0)), approximations were computed using **Eqs. 2.4.22** which are valid under the conditions that (*η*_*S*_, *η*_*I*_, *ε*_*S*_, *ε*_*I*_) → (0,0,0,0) and the φ-approximations which are the solutions of **Eqs. 2.5.1.1-2** as given in **Appendix A** in a parametric form where τ act as the parameter. For the partial competitive inhibition scheme (**B, D, F** where trajectories start at (I, S) = (1,1) and end at (I,S) = (*I*_∞_,0)), post-steady state approximations were generated under the conditions that (*η*_*S*_, *χ*_*I*_) → 0 and the pre-steady state approximations were computed for (*ε*_*S*_, *χ*_*I*_, *ε*_*I*_) → (0,0,0) using the conservation law *I* = 1 - *ε*_*I*_*Y* (using **Eqs. 2.9.3.10** and **2.9.4.9** respectively with *S* ∈ [0,1] as the parameter). Common settings are *η*_*S*_ = 0.02, *κ*_*S*_ = 0.1, *η*_*I*_ = 0.03, *χ*_*I*_ = 0.03, *κ*_*I*_ = 0.5, *σ* = 1. **A-B**. *ε*_*S*_ = 5.5, *ε*_*I*_ = 0.06, *ρ* = 137.5, *δ* = 0.07, *γ* = 0.022.**C-D**. *ε*_*S*_ = 0.5, *ε*_*I*_ = 0.06, *ρ* = 12.5, *δ* = 0.634, *γ* = 0.024. **E-F**. *ε*_*S*_ = 0.05, *ε*_*I*_ = 0.06, *ρ* = 1.5, *δ* = 2.314, *γ* = 0.24.

In case of fully competitive inhibition scheme, one finds that 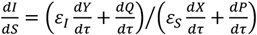 which follows from *I* = 1 − *ε*_*I*_ *Y* − *Q*. Here the product levels (*P,Q*) are monotonically increasing functions of time so that 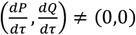 throughout the entire timescale regime except at τ = 0 and τ → ∞. Unlike the partial competitive inhibition, in case of fully competitive inhibition scheme 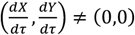 at τ = 0 and 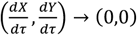 at τ → ∞. The conditions 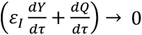 and 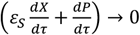 will be true when τ → ∞. This means that when τ → ∞, then 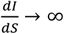 or undefined. When there is a significant mismatch in the steady state timescales, then one can still observe a time point at which 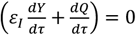 and 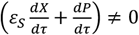 leading to 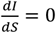 in the (I, S) space of the fully competitive inhibition scheme. Contrasting from the partial competition, the time at which 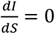 may not be equal to the steady state timescale corresponding to the enzyme-inhibitor complex. Various ways to approximate the steady state substrate and inhibitor levels described in section **2.9.5** are demonstrated in **Figs. 15**. Results clearly suggest that the pre-steady states in the (V, S) and (U, I) spaces can be well approximated by *V* ≅ 1 − *S* and *U* ≅ *ρ*(1 − *I*) and **Eqs. 2.9.5.13-14** can predict the steady state substrate level S_C_ and inhibitor level I_C_ very well. All the trajectories in the (V, S) space will be confined by the triangle formed by the lines *V* = 1 − *S*, V = 0 and S = 0. Similarly, all the trajectories in the (U, I) space will be confined by the triangle formed by the lines *U* = *ρ*(1 − *I*), U = 0 and I = 0.

**FIGURE 15.**
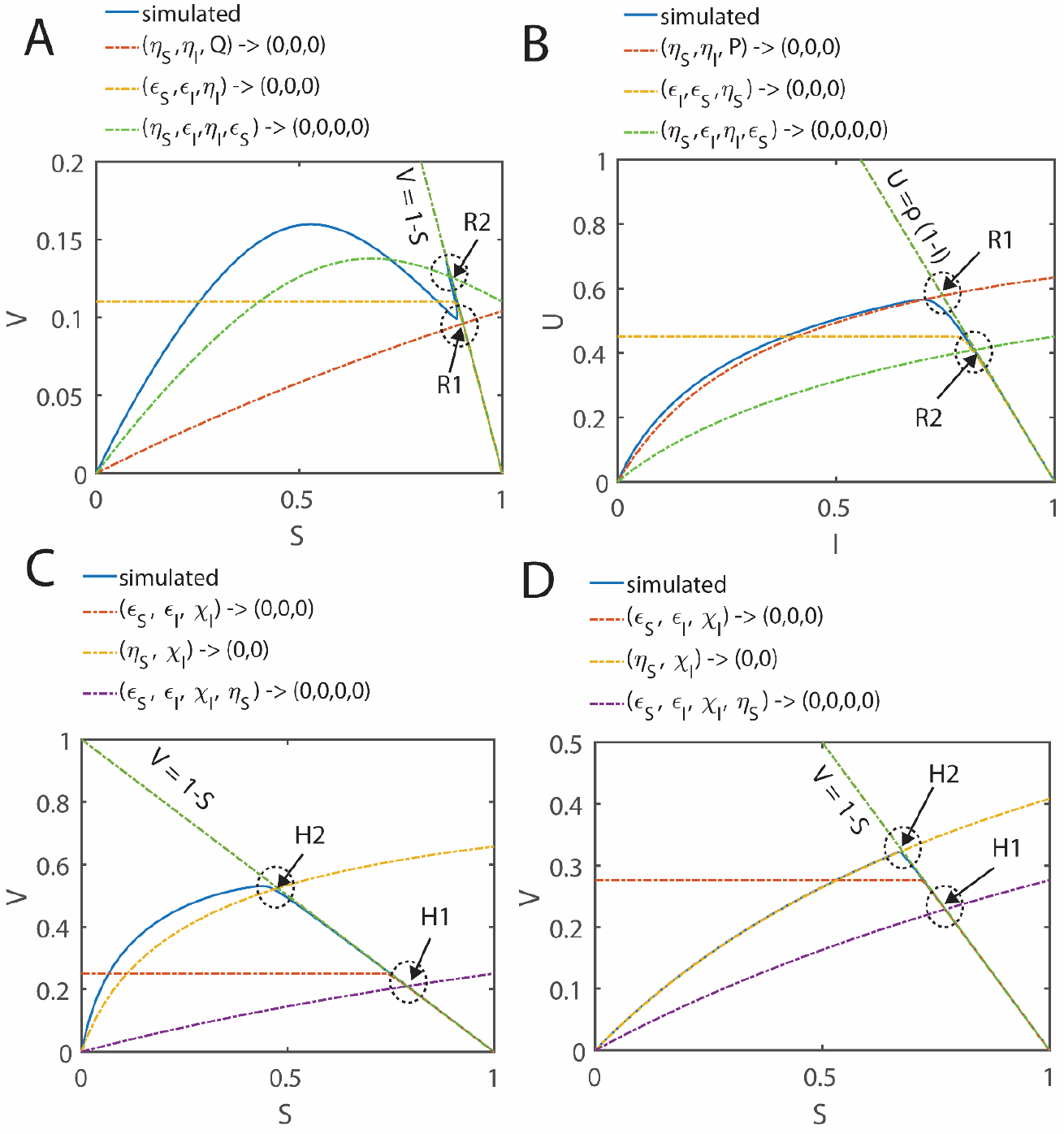
Approximation of the steady state substrate and inhibitor levels corresponding to fully (A, B) and partial competitive (C, D) schemes. Common initial conditions for the simulation of fully competitive inhibition **Eqs. 2.2.7-9** are (*S, I, E, X, Y, P, Q*) = (1,1,1,0,0,0,0) and for partial competitive inhibition **Eqs. 2.9.5-7** are (*S, I, E, X, Y, P*) = (1,1,1,0,0,0) at τ = 0. A-D clearly show that (*P, Q*) *≅* (0,0) in the pre-steady state regime so that *V ≅* 1 - *S* and *U ≅ ρ*(1 - *I*). These lines intersect the post-steady state approximations near the original steady state. In case of fully competitive inhibition, we considered the intersection (R1, R2) between the post steady state approximations under the conditions that (*η*_*S*_, *η*_*I*_, *ε*_*S*_, *ε*_*I*_) → (0,0,0,0) (**Eqs. 2.4.23**), (*η*_*S*_, *η*_*I*_, *Q*) → (0,0,0) (**Eq. 2.4.16**), (*η*_*S*_, *η*_*I*_, *P*) → (0,0,0) (**Eq. 2.4.12**) and the pre-steady state approximations under the conditions that (*η*_*S*_, *ε*_*S*_, *ε*_*I*_) → (0,0,0) (**Eq. 2.6.10**) and (*η*_*I*_, *ε*_*S*_, *ε*_*I*_) → (0,0,0) (**Eq. 2.6.23**) along with *V ≅* 1 - *S* and *U ≅ ρ*(1 - *I*). In case of partial competitive inhibition, we considered the intersections (H1, H2) between the post-steady state approximations under the conditions that (*η*_*S*_, *χ*_*I*_, *ε*_*S*_, *ε*_*I*_) → (0,0,0,0) (**Eqs. 2.9.2.2**) and (*η*_*S*_, *χ*_*I*_) → (0,0) (**Eq. 2.9.4.7**) and the pre-steady state approximations under the conditions that (*χ*_*I*_, *ε*_*S*_, *ε*_*I*_) → (0,0,0,0) (**Eq. 2.9.3.8**) along with *V ≅* 1 - *S*. The settings are as follows. **A-B**. *η*_*S*_ = 0.02, *κ*_*S*_ = 1.1, *κ*_*I*_ = 0.2, *η*_*I*_ = 0.03, *ε*_*S*_ = 0.6, *ε*_*S*_ = 0.4, σ = 1, ρ = 2.25, δ = 0.244, ϒ = 3.67. **C**. *η*_*S*_ = 0.01, *κ*_*S*_ = 0.01, *κ*_*I*_ = 0.05, *η*_*I*_ = 0.02, *ε*_*S*_ = 0.9, *ε*_*S*_ = 0.5. **D**. *η*_*S*_ = 0.01, *κ*_*S*_ = 1, *κ*_*I*_ = 5, *η*_*I*_ = 0.02, *ε*_*S*_ = 0.9, *ε*_*S*_ = 0.5.

## 5. Conclusion

Fully and the partial competitive inhibition of the Michaelis-Menten enzyme kinetics play critical role in designing drug molecules against the nodal enzymes of various harmful pathogens. Designing of such drug molecules involves screening of various substrate like small molecules which can act as potential inhibitors of the target enzymes. Estimation of various kinetic parameters associated with the competitive inhibition is essential for such comparative studies and evaluation of various potential drug candidates. The currently available standard quasi steady state approximation with stationary reactant assumption is applicable only in the post-steady state regime of the velocity-substrate-inhibitor space and it is significantly limited by the vast number of conditions of validity. Particularly, this approximation will not work when the concentration of the enzyme is equal to or higher than the substrate.

In this context, we have derived several approximations under various conditions of validity over both pre- and post-steady state regimes of the velocity-substrate-inhibitor spaces of fully and partial competitive inhibition schemes. Our detailed analysis yielded refined expressions over the currently available standard quasi steady state approximation with stationary reactants assumption. We have shown that these refined expressions are valid for wide ranges of enzyme to substrate and inhibitor ratios. Further, we have shown for the first time in the literature that the enzyme-inhibitor-substrate system can exhibit temporally well separated two different steady states with respect to both enzyme-substrate and enzyme-inhibitor complexes under certain conditions. When the total substrate and inhibitor levels are higher than the enzyme level, then one can define 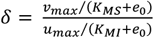 which plays critical role in controlling the phase-space dynamics associated with the relative speed of binding, dissociation and conversion into the products of the enzyme-inhibitor and enzyme-substrate complexes of the fully competitive enzyme inhibition scheme.

The ratios *f*_*S*_ = *v*_*max*_⁄(*K*_*MS*_ + *e*_0_) and *f*_*S*_ = *u*_*max*_⁄(*K*_*MI*_ + *e*_0_) are the acceleration factors with respect to the conversion dynamics of substrate and inhibitor into their respective products. When 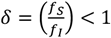, then the speed of conversion of the enzyme-inhibitor complex into the product will be faster than the speed of conversion of the enzyme-substrate complex into the respective product. As a result, the enzyme-substrate complex will exhibit multiple steady states and eventually can reach the full-fledged steady state value only after the depletion of enzyme-inhibitor complex. On the other hand, when *δ*> 1, then the enzyme-inhibitor complex will exhibit multiple steady states and eventually can reach the full-fledged steady state value only after the depletion of enzyme-substrate complex.

This complicated behavior of the enzyme-substrate-inhibitor system especially when *δ*≠ 1 poses enormous difficulties in generating consistent experimental datasets on the steady state velocities versus substrate and inhibitor levels and also introduces large amount of error in the estimation of various kinetic parameters from these datasets both in the cases of fully and partial competitive inhibitions. Remarkably, our refined expressions for the reaction velocities over enzyme-substrate-inhibitor space can control this error more significantly than the currently available standard QSSA velocity expressions.

## Appendix A

Consider the following set of coupled linear second order ODEs.

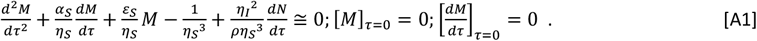

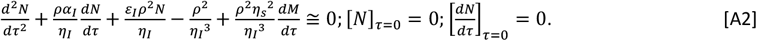

This system can be uncoupled as follows. Upon differentiating **Eqs. A2** with respect to τ, one arrives at the following third order ODE.

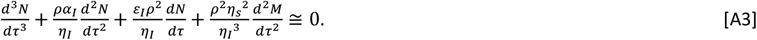

From **Eqs. A1**, one can derive the following expression.

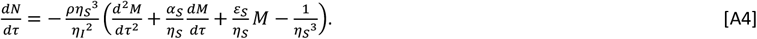

By repeated differentiation of **Eq. A4** with respect to τ, we obtain the following relationships.

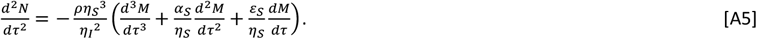

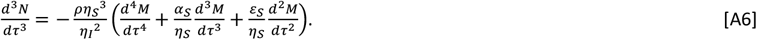

Upon substituting **Eqs. A4-A6** into **Eq. A3**, one obtains the following uncoupled fourth order ODE corresponding to (M, τ) space.

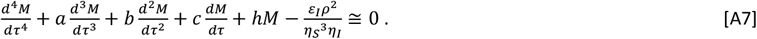

In this equation, various terms a, b, c and h are defined as follows.

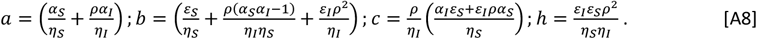

The first two initial conditions associated with **Eq. A7** are as follows.

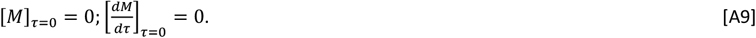

Other two initial conditions directly follow from the initial conditions corresponding to N.

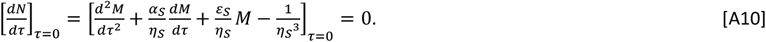

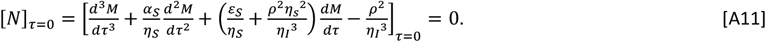

Upon obtaining the solution for the (M, τ) space, one can directly obtain the expression corresponding to the (N, τ) space as follows.

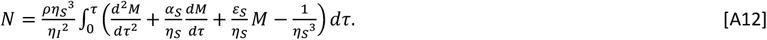

Similar to **Eqs. A3-A12**, one can also derive the following solution set. From **Eqs. A2**, one can derive the following expression.

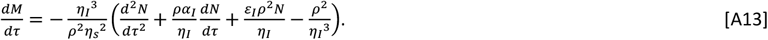

By repeated differentiation of **Eq. A13** with respect to τ, we obtain the following relationships.

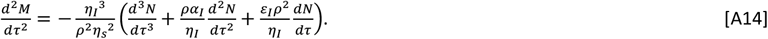

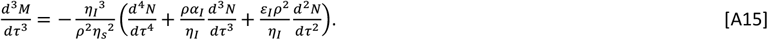

Upon differentiating **Eq. A1** with respect to τ, one obtains the following expression.

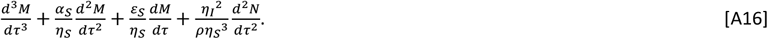

Upon substituting the derivatives from **Eqs. A13-A15** into **Eq. A16**, one obtains the following uncoupled fourth order ODE corresponding to (N, τ) space.

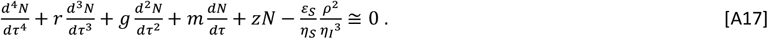

Here the parameters r, *g*, m, and z are defined as follows.

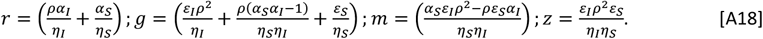

Upon obtaining the solution for the (N, τ) space one can directly obtain the expression corresponding to the (M, τ) space as follows.

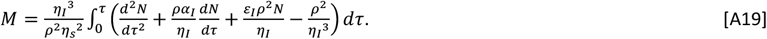

The initial conditions corresponding to the fourth order uncoupled ODE given by **Eqs. A17** can be written as follows.

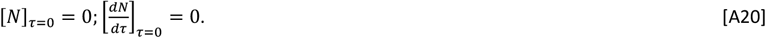

Other two initial conditions directly follow from the initial conditions corresponding to M.

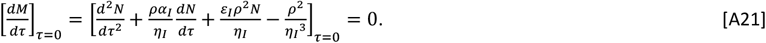

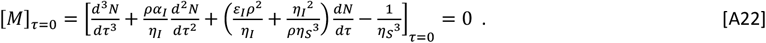

Solution to **Eqs. A1**-**A2** can be written as follows.

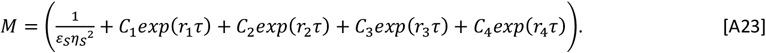

From this equation, N can be obtained using **Eq. A12**. Here r_1_, r_2_, r_3_, and r_4_ are the roots of the following fourth degree polynomial in r.

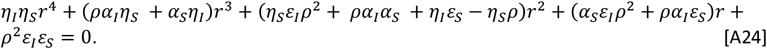

Upon using the initial conditions given by **Eqs. A9**-**11**, one can obtain the expressions for various constant terms C_1_-C_4_ in **Eqs. A23** as follows.

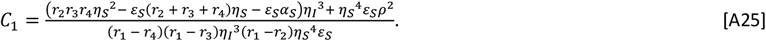

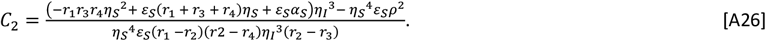

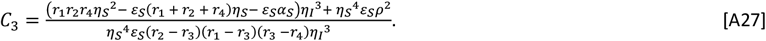

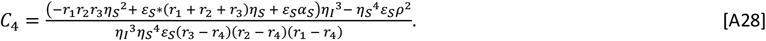

Using the transformations (*P,V*) = *ε* _*S*_ *η*_*S*_^2^(*M,F*) and (*Q,U*) = *ε* _*I*_ *η*_*I*_ ^2^(*N,G*) one can revert back to the original dynamical variables V, U, P and Q. Subsequently one finds that S = 1 – V – P and I = 1 – U/ρ – Q. Here the dynamical variables (S, I, V, U, P, Q) are all expressed over (V, S, I), (U, S, I), (V,P,Q) and (U, P, Q) spaces in the parametric form where τ acts as the parameter.

## Appendix B

Let us consider the following nonlinear first order ODE along with the initial condition.

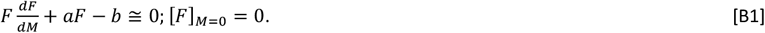

Here 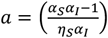 and 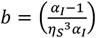. Upon solving **Eq. B1** implicitly and reverting back to (V, P) space using the transformation 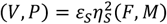, one obtains the following solution.

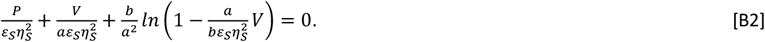

Upon substitution of the conservation law, P = 1 – V – S in **Eq. B2** and after few manipulations and rearrangements after adding (*a* − 1) term both sides of equation, one finds the following implicit solution corresponding to the (V, S) space.

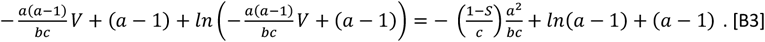

Here 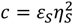. Upon exponentiating both sides of **Eq. B3**, one arrives at the form *Wexp*(*W*) = *Z* as follows.

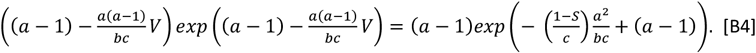

Solution to **Eq. B4** can be written in terms of Lambert W function as follows.

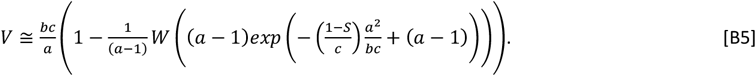

Upon inserting the corresponding values of a, b and c in **Eq. B5**, one finally arrives at the following approximation in the pre-steady state regime of the (V, S) space.

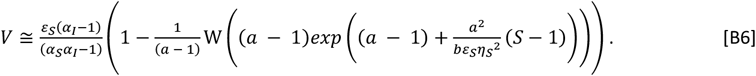

Let us consider the following properties of the Lambert’s W(Z) function.

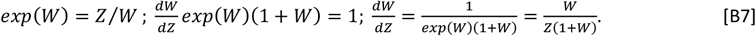

Upon expanding the right-hand side of **Eq. B5** using **Eq. B7** in a Taylor series around S = 1, noting that *WZexp*(*Z*) = *Z*, one finds the following series.

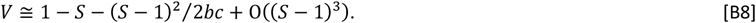

Since the amount of product building up in the pre-steady state regime will be negligible in most of the scenarios, one can ignore the second order terms in **Eq. B8** and use the pre-steady state approximation as *V* ≅ 1 − *S* in the (V, S) space.

